# OTRec: A Deep Learning Recommender for Druggable Disease–Target Prioritization

**DOI:** 10.64898/2025.12.21.695803

**Authors:** Dan Ofer, Michal Linial

## Abstract

**Objective.:** Identifying druggable disease–target associations remains a central challenge in translational medicine, limiting therapeutic discovery and repurposing. We rank, for every disease in the Open Targets Platform (OTP), every gene in the druggable genome by its likelihood of entering clinical trials.

**Methods.:** Here we present OTRec, a deep learning–based recommender system that ranks such associations at scale. Unlike approaches that rely on manually curated or aggregated evidence scores, OTRec employs a two-tower architecture to learn latent representations from 663,351 disease–target pairs. The model integrates heterogeneous inputs, including textual descriptions, ontology-derived features, and biological annotations such as tractability, Gene Ontology (GO) terms, and pathway information. We evaluated OTRec by target-disjoint cross-validation and by a release-based temporal protocol, training on OTP Release 22.02 and testing against clinical evidence added by 2025.

**Results.:** OTRec improves on the retrospective OTP association score both in temporal validation (ROC-AUC: 0.872 vs. 0.559; PR-AUC: 0.288 vs. 0.082) and in cross-validation (ROC-AUC 0.950 and PR-AUC 0.844, against 0.914 and 0.455). Across 1,455 temporal-test diseases with a later positive target, OTRec places a true target first for 46.3% of diseases, compared with 41.8% for a matched model, and reaches ROC-AUC 0.887 on cold-start targets with no 2022 clinical indication.

**Conclusion.:** We rank the 4,479 genes of the druggable genome across *∼*19,000 OTP diseases and release 214,968 previously unlinked candidate associations with score *≥*0.65, covering 4,347 diseases, including 2,322 target-orphan diseases (with no clinically associated target in OTP).

**Availability.:** Code, models, ranked predictions, and extended tables are available at https://github.com/LinialLab/OTRec, together with an interactive prediction platform https://huggingface.co/spaces/GrimSqueaker/ OTRec.

## 1. Introduction

Drug discovery is hindered by a pervasive “streetlight effect”: research concentrates on well-studied targets while large regions of the druggable genome remain under-investigated [1–3]. The Open Targets platform (OTP) integrates diverse evidence sources, including GWAS associations, somatic mutations, animal models, drug and chemical data, and text-mined literature, but the resulting association scores are intrinsically retrospective, reflecting existing knowledge rather than a target’s prospective therapeutic promise [4]. A zero or low score may signal the absence of investigation rather than a genuine lack of relevance. Moreover, OTP is built on manually designed, rule-based scoring, and such frameworks may underperform data-driven, learned approaches, at least in structure-based virtual screening [5].

We therefore developed **OTRec** (Open Targets Recommender). The task is: given a disease, rank every gene in the druggable genome by its probability of entering clinical trials, a label OTP captures across *∼*19,000 diseases. In this work, “target” denotes an OTP gene target (mapped to an Ensembl gene ID), and “drug” denotes therapeutic evidence represented in OTP via approved or clinical-stage associations linked to those targets. Importantly, OTRec ranks disease–target pairs. It does not predict a compound, direction of modulation, dose, tissue context, or safety. OTRec is designed to generalize to unobserved disease–target pairs, including targets absent from training and existing data, using disease and target annotations and attributes. Our temporal evaluation asks a prospective ranking question: can we predict *future* clinical labels?

## Statement of Significance

|  |  |
| --- | --- |
| **Problem or Issue** | Evidence-based disease–target scores are retrospective and may disadvantage poorly studied targets and rare or orphan diseases. |
| **What is Already Known** | The Open Targets Platform aggregates heterogeneous evidence into rule-based association scores. Existing OTP-based methods have largely been evaluated retrospectively, while drug–target interaction models require a compound as input. |
| **What this Paper Adds** | OTRec formulates disease–target prioritization as content-based retrieval from biomedical text, ontology, and target annotations. We introduce a release-based 2022→2025 benchmark of clinical progression, demonstrate improved disease-centric ranking and cold-start performance, and release ranked predictions, benchmark splits, code, models, and an interactive platform. |
| **Who would benefit from the new knowledge in this paper** | Researchers and drug-development teams prioritizing therapeutic targets, especially for rare, neglected, or target-orphan diseases, and researchers developing computational target-prioritization methods. |

## 2. Related Work

Disease–target prioritization is a recommendation problem over biomedical knowledge, and a distinct task from drug–target interaction (DTI) prediction.

The closest prior work on Open Targets is Han et al. [6], which trains gradient boosting on five OTP association scores plus five computed target-similarity features. Its published results use OTP Release 21.04 and a random 70/30 target split, and neither runnable code nor processed data are available, so we quote them as context rather than as a matched benchmark. Genetics-guided prioritization is a parallel approach: Duffy et al. [7] rank 19,365 genes across 399 indications from human genetic evidence, including a direction-of-effect variant. That score is target-centric and evidence-driven; OTRec learns disease-conditional rankings from text and ontology.

The DTI literature is larger, but solves a different problem: DTI models predict whether a compound binds a protein, either from a curated biomedical graph [8–10] or from SMILES and protein sequence [11, 12]. No compound, structure, or binding affinity enters OTRec’s inputs or its labels, so DTI models cannot be run on this task without redefining it. Graph methods have a further limitation: their edges record what has already been studied. We include Matrix Factorization, Node2Vec, and TF-IDF as cold-start controls. Pretrained language models (BioBERT [13], PubMedBERT [14]) supply biomedical text representations, and benchmarking suites such as Therapeutics Data Commons [15] supply standardized splits, but neither defines a disease– target prioritization task with release-separated clinical-trial-entry labels at OTP scale. We introduce this benchmark and release splits, code, and candidate predictions to enable direct replication and comparison.

## 3. Methods

### 3.1. Problem Formulation and Data

We formulate target prioritization as an implicit-feedback recommendation task: rank candidate human protein-coding genes for a given disease by likelihood of clinical relevance. We used OTP Release 25.06 for cross-validation (CV) and inference, and Release 22.02 as the historical training snapshot for temporal validation [1, 16–18]. Candidate rows are drawn from the OTP direct-association table, so every constructed pair carries at least one direct evidence source. Positive labels (*y*=1) are pairs with evidence of clinical progression (clinical trials or approved drugs) in the relevant OTP known-drug / clinical-evidence table; the remaining rows — pairs with direct evidence but no recorded clinical progression — are implicit negatives (*y*=0). The implicit-feedback formulation conflates true negatives with unlabeled positives; this is a positive–unlabeled setting, with unlabeled examples treated as negatives. The temporal split (Section 3.3) re-labels pairs whose clinical evidence appears only in a later OTP release.

Throughout, “drug” follows OTP usage and covers any therapeutic entity in the OTP drug index (*N* =18,081 entities in Release 2025.06, of which 10,956 have reached clinical development, phase 1–4). Altogether, the OTP drug index is predominantly small molecules (82%), with antibodies, proteins, gene therapies, and vaccines making up the remainder. “gene” / “target” refers to the human protein-coding subset of the Finan et al. druggable genome [19] (4,479 genes, out of 20,130 protein-coding genes in OTP). Non-coding loci catalogued by OTP (lncRNAs *N* =34,882, miRNAs *N* =1,879, pseudogenes) are excluded from both the training label set and the inference pool, mirroring the small-molecule bias of the OTP drug index.

We applied filtering criteria aligned with prior work [6]: non-specific ontology terms (“measurement”, “phenotype”, “biological process”, “cell proliferation disorder”) were excluded. We restrict training to *∼*1,500 targets with clinical associations across at least two indications. This prevents a target with one recorded indication from being learned almost entirely through that disease (stratified results in Section 4.3). The training set comprises 663,351 disease– target pairs across 12,337 diseases (Release 25.06 frame), including 2,329 positive-bearing diseases and 67,532 positives (10.18%). During inference, the model ranks the full druggable genome [19] (4,479 genes) across *∼*19,000 valid OTP diseases, including pairs with no direct evidence and hence absent from the training frame.

### 3.2. Model Architecture and Features

OTRec uses a multi-input two-tower architecture (Figure 2) that separately encodes disease and target representations, enabling the model to learn semantic and biological attributes from text and ontology structure.

**Figure 1:**
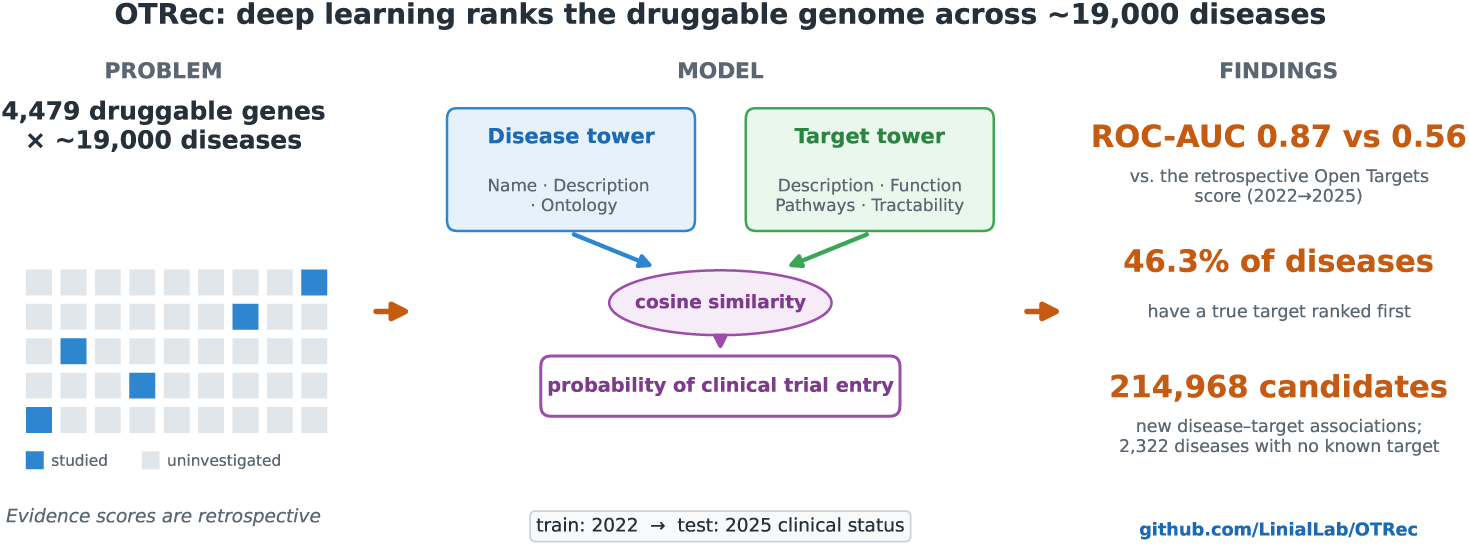
Graphical Abstract

**Figure 2:**
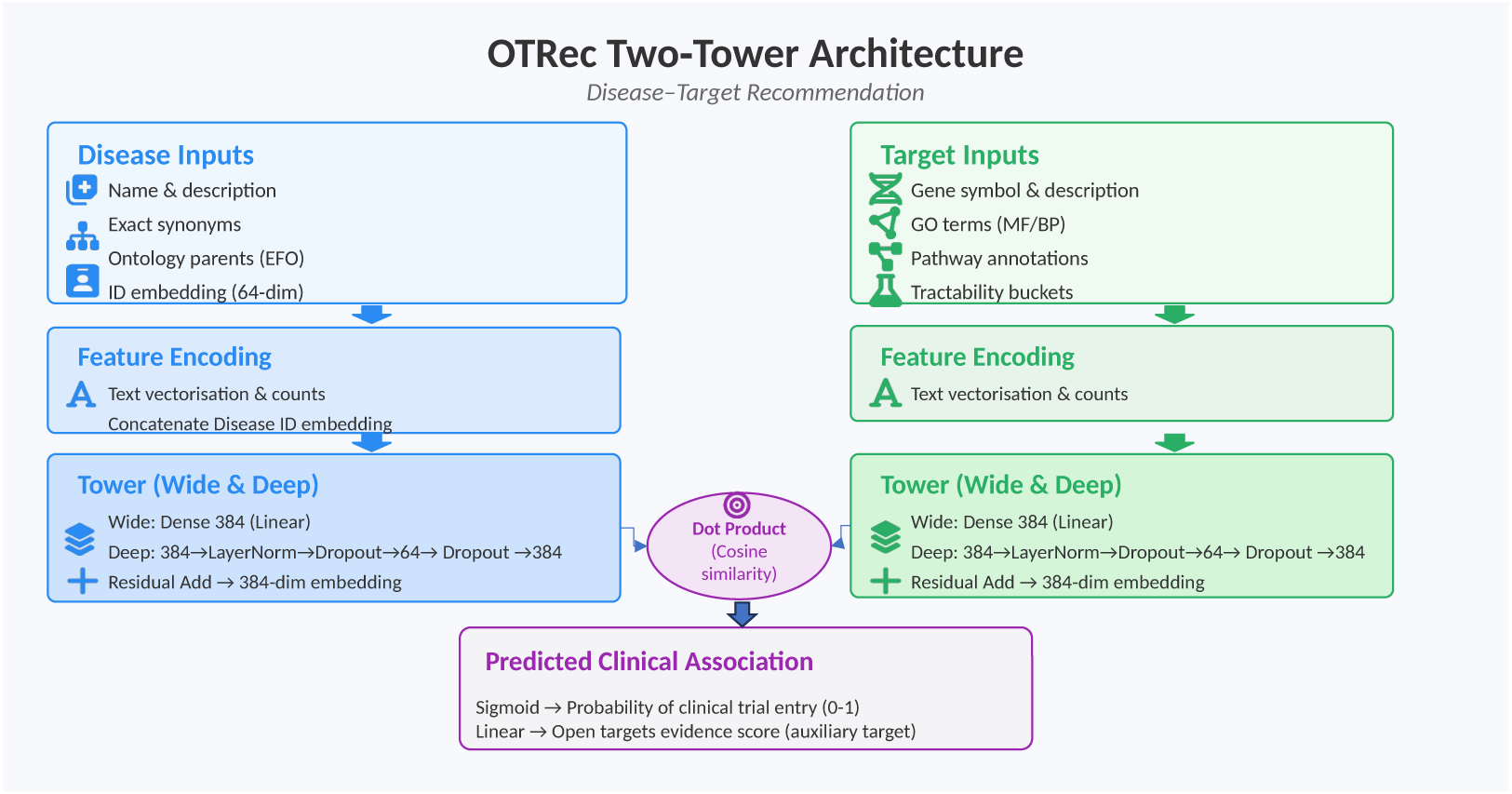
OTRec architecture. The two-tower model separately encodes disease and target features (text, ontology, tractability, gene) into 384-dimensional embeddings, compared by normalized dot product (cosine similarity) to predict clinical trial entry.

#### 3.2.1. Inputs

Each tower vectorizes its text fields with Keras [20] TextVectorization in count mode (bag-of-words). The *disease tower* uses disease names, descriptions, exact synonyms, therapeutic areas, phenotypes, and ontology parents (Experimental Factor Ontology (EFO) terms [21]), concatenated with a learned disease-ID embedding (*d*=64). The *target tower* uses gene symbol, name, synonyms, function descriptions, Gene Ontology terms [22], pathway annotations including Reactome-derived terms [23], UniProt annotations [24], and OTP target prioritization / tractability features including gnomAD-derived genetic constraint scores where available [16, 25].

#### 3.2.2. Architecture and Training

Each tower applies a residual “Wide & Deep” block [26], combining a memorization-oriented linear (“wide”) path with a generalization-oriented nonlinear (“deep”) path: a direct linear projection (*d*=384) summed with a layer-normalized nonlinear bottleneck path (Dense 384*→*64*→*384, ELU activations, Dropout 0.15–0.35). The 384-dimensional embeddings interact via normalized dot product (cosine similarity), followed by a multi-task head: a sigmoid output predicts clinical trial entry while an auxiliary linear head predicts the OTP overall association score from the same release used to construct the training split (2025.06 in target-disjoint CV, 2022.02 in the temporal setting). In the temporal setting, clinical labels and the auxiliary OTP score are from Release 22.02, while the annotation strings come from a single later OTP snapshot shared by the temporal training and test cohorts, as the 2022 ontology lacks roughly half of the evaluation cohort’s disease terms. Training used TensorFlow/Keras [20, 27], Adam (learning rate 7*×*10*^−^*^3^, batch size 512), binary cross-entropy loss, ReduceLROnPlateau (factor 0.2, patience 1), and early stopping (patience 2) on a target-disjoint validation subset of the training data. The auxiliary loss weight was 0.2 in target-disjoint CV and 0.1 in the temporal runs. Hyperparameters were not searched; all settings are fixed in the released code.

### 3.3. Evaluation and Baselines

#### 3.3.1. Target-Disjoint Cross-Validation

To evaluate generalization to novel targets unseen during training, we use 5*×*5-fold target-disjoint cross-validation (25 folds). In each fold, 20% of targets are withheld; all associations involving held-out targets are removed from training. This is attribute-cold-start rather than annotation-free cold-start: held-out targets remain represented by text, ontology, pathway, and tractability features.

#### 3.3.2. Temporal Validation

A model trained on OTP Release 22.02 is evaluated on pairs whose clinical label changed by Release 25.06. Temporal labels are defined by release difference: we construct the 2025 cohort and remove any exact (diseaseId, targetId, label) tuple already present in 2022. The retained pairs are those absent from the 2022 frame and those whose label changed; all such negatives are kept rather than sampled. The resulting test set contains 403,132 pairs (16,074 positives, 3.99%) over 11,624 diseases and 1,521 targets, compared with 9,741 diseases in the 2022 training frame — a strong covariate shift relative to target-disjoint CV in both disease coverage and prevalence. We report the area under the receiver operating characteristic curve (ROC-AUC) and the area under the precision–recall curve (PR-AUC) over all test pairs, together with disease-centric shortlist metrics defined in Section 4.2.

#### 3.3.3. Baselines

We benchmark against nine methods including learned models, retrospective priors, and cold-start controls: (*i*) *OTTree*: CatBoost [28] trained on the same OTRec feature set, isolating the contribution of the two-tower architecture from the feature engineering; (*ii*) *Han et al.* [6]: XGBoost on five OTP association scores (genetic association, RNA expression, affected pathway, animal model, somatic mutation) plus five computed target-similarity features, quoted from the original publication; (*iii*) *Open Targets score*: the OTP overall association score, which itself aggregates clinical and known evidence and is a retrospective baseline; (*iv*) *Disease Mean*: the per-disease positive rate computed on the training split, broadcast to every candidate target for that disease; (*v*) *Target Mean*: the per-target positive rate computed on the training split, broadcast to every candidate disease for that target; (*vi*) *Matrix Factorization*: TruncatedSVD on the binary positive interaction matrix; (*vii*) *Node2Vec* [29]: bipartite graph of positive disease–target pairs; node embeddings learned via random walks (dimensions 64, walk length 20, walks per node 10, context window 10, 3 epochs); test-disease embeddings are scored by cosine similarity to the mean training-target embedding, so this control is constant across targets within a disease; (*viii*) *TF-IDF cosine*: lexical similarity between concatenated disease and target text fields; (*ix*) *Frozen BioClinical ModernBERT + MLP* [30]: static disease and target embeddings from BioClinical ModernBERT scored with a small MLP head over concatenation, absolute difference, elementwise product, and cosine similarity; test targets are unseen during training.

## 4. Results

### 4.1. Benchmark Comparison

In the 5*×*5 target-disjoint CV (Table 1, Figure 3A), OTRec achieves ROC-AUC 0.950 / PR-AUC 0.844, marginally surpassing OTTree (0.947 / 0.772) and exceeding the OTP score (0.914 / 0.455). The matched architecture comparison is OTRec versus OTTree: the ROC-AUC difference is small (+0.003), but OTRec improves PR-AUC by 0.072 and ranks true targets higher (Section 4.2). The BioClinical ModernBERT encoder [30] sits above the simple priors but below the top models on PR-AUC (0.666 against 0.844 for OTRec). Among the simple controls, Disease Mean is the strongest historical prior (0.873 / 0.466), whereas a leakage-free Target Mean collapses to the global prior (0.500 / 0.102) because every test target is unseen in its evaluation fold. Matrix Factorization, Node2Vec, and TF-IDF remain much weaker than OTRec, indicating that the target-disjoint regime genuinely tests generalization to novel targets and that OTRec’s advantage comes from learned content transfer rather than surface text or interaction memorization.

**Table 1:** Target-disjoint cross-validation (mean *±* SD over 25 folds). A horizontal rule separates ML baselines from cold-start controls; Han et al. is quoted from the original publication (different OTP release, random 70/30 target split).

| Model | ROC-AUC | ±SD | PR-AUC | ±SD |
| --- | --- | --- | --- | --- |
| **OTRec (ours)** | **0.950** | 0.011 | **0.844** | 0.017 |
| OTTree (CatBoost) | 0.947 | 0.005 | 0.772 | 0.023 |
| Han et al. (2022) | 0.910 | 0.014 | 0.726 | 0.046 |
| Frozen BioClinical ModernBERT + MLP | 0.912 | 0.008 | 0.666 | 0.028 |
| Open Targets score | 0.914 | 0.005 | 0.455 | 0.030 |
| Disease Mean | 0.873 | 0.004 | 0.466 | 0.022 |
| Target Mean | 0.500 | 0 | 0.102 | 0.008 |
| Matrix Factorization | 0.811 | 0.008 | 0.265 | 0.014 |
| Node2Vec | 0.684 | 0.006 | 0.139 | 0.013 |
| TF-IDF cosine | 0.513 | 0.016 | 0.106 | 0.009 |

**Figure 3:**
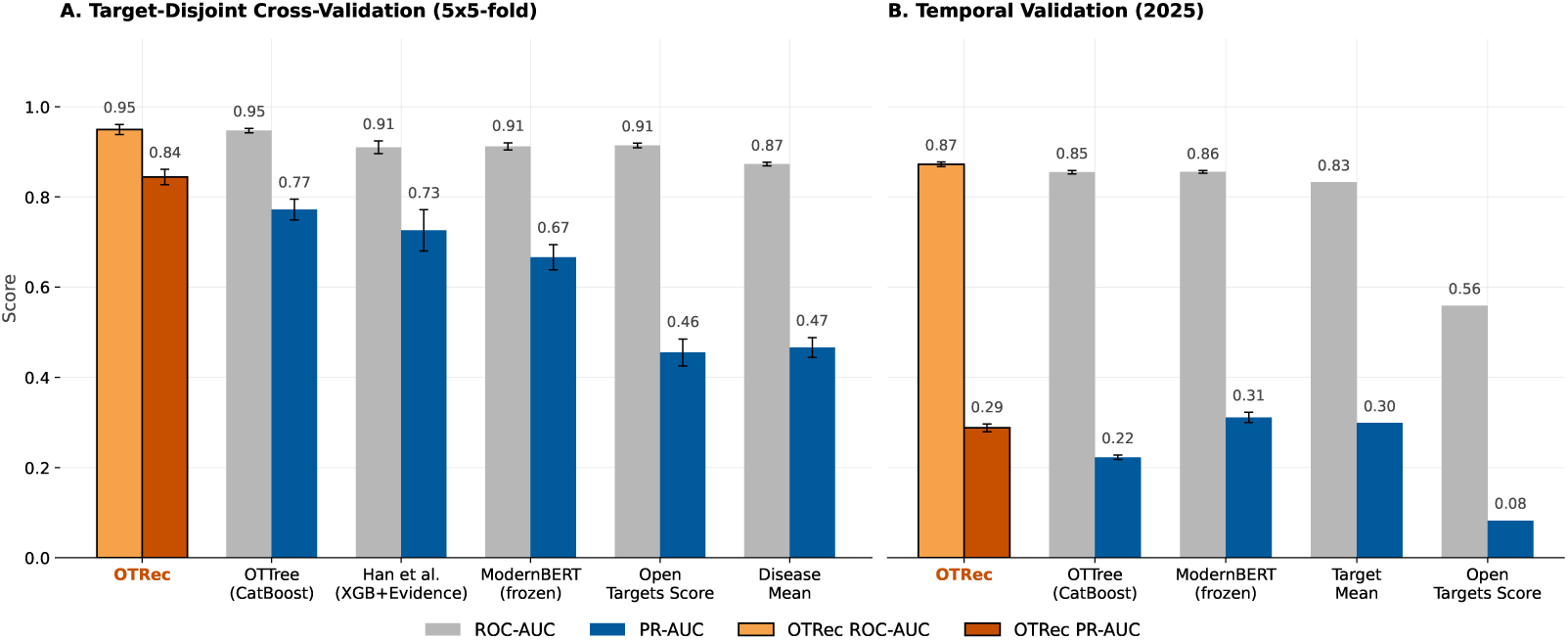
Benchmark performance. **(A)** Target-disjoint 5*×*5 cross-validation results. **(B)** Release-based temporal validation using 2022 clinical labels and 2025 outcomes. Error bars denote standard deviation (25 folds in A, five runs in B). OTRec (orange) attains the best ROC-AUC in both settings and the best PR-AUC in cross-validation.

In the temporal setting (Table 2, Figure 3B), OTRec improves over the OTP score on ROC-AUC by 0.313 (0.872 *±* 0.005 vs. 0.559), and reaches PR-AUC 0.288 *±* 0.009. BioClinical ModernBERT leads temporal PR-AUC (0.311 *±* 0.011), ahead of the Target Mean prior (0.299) and of OTRec, though its pretraining corpus post-dates the 2022 cutoff; the disease-only prior is far weaker (0.052). Unlike in target-disjoint CV, nearly all temporal test targets appear in the 2022 training frame, so their historical positive rates are informative. Matrix Factorization, Node2Vec and TF-IDF cosine are substantially weaker on both metrics. Temporal prevalence is 3.99% against 10.18% in target-disjoint CV, so PR-AUC is not comparable across the two settings. OTRec’s advantage over these historical priors is concentrated in pair-and disease-level ranking (Section 4.2).

**Table 2:** Release-based temporal validation: 2022 clinical labels, 2025 outcomes (mean *±* SD over five runs). “–”: deterministic baselines and the single Node2Vec run.

| Model | ROC-AUC | ±SD | PR-AUC | ±SD |
| --- | --- | --- | --- | --- |
| **OTRec (this study)** | **0.872** | 0.005 | 0.288 | 0.009 |
| OTTree (CatBoost) | 0.855 | 0.004 | 0.223 | 0.005 |
| Frozen BioClinical ModernBERT + MLP | 0.856 | 0.003 | **0.311** | 0.011 |
| Target Mean | 0.832 | – | 0.299 | – |
| Matrix Factorization | 0.553 | – | 0.050 | – |
| Node2Vec | 0.533 | – | 0.042 | – |
| TF-IDF cosine | 0.512 | – | 0.042 | – |
| Disease Mean | 0.594 | – | 0.052 | – |
| Open Targets score | 0.559 | – | 0.082 | – |

### 4.2. Disease-Centric Shortlist Utility

ROC-AUC and PR-AUC measure pairwise discrimination, but curators typically use shortlists. We re-analyzed target-disjoint out-of-fold predictions disease-by-disease, restricting to the 2,329 diseases with at least one held-out positive target (Table 3).

**Table 3:** Disease-centric shortlist performance in target-disjoint cross-validation across 2,329 diseases with at least one held-out positive target. Hit@*k* is the proportion of diseases with at least one true positive among the top *k* targets; P@5 and R@5 are mean precision and recall at rank 5; MRR is the mean reciprocal rank of the first true positive.

| Model | Hit@1 | Hit@5 | P@5 | R@5 | Hit@10 | Hit@25 | MRR |
| --- | --- | --- | --- | --- | --- | --- | --- |
| **OTRec** | **0.682** | **0.806** | **0.645** | **0.321** | 0.848 | 0.897 | **0.741** |
| OTTree (CatBoost) | 0.487 | 0.732 | 0.537 | 0.272 | 0.848 | **0.951** | 0.600 |

OTRec placed a true positive at rank 1 for 68.2% of diseases (vs. 48.7% for OTTree), and led on the mean reciprocal rank (MRR), the mean of 1*/*rank of the first true positive. OTRec leads at *k≤*5; OTTree recovers more positives at *k*=25.

We repeated this analysis on the temporal test set, over the 1,455 diseases with at least one later positive target (Table 4; ties are assigned the worst rank in their block). OTRec leads OTTree, Target Mean and the OTP score on every shortlist metric. This holds on poorly annotated diseases: across the 610 diseases with no clinically associated target in 2022, OTRec reaches Hit@1 0.575 and MRR 0.680, against 0.538 / 0.636 for Target Mean and 0.139 / 0.196 for the OTP score. Stratified by therapeutic area, by target annotation volume and by number of known 2022 targets (Supplementary S8), OTRec leads OTTree on MRR in every subgroup except cardiovascular disease (0.472 vs. 0.474) and the 106 diseases with 1–2 known targets (0.603 vs. 0.623). These subgroups also separate the two kinds of ranking. In several therapeutic areas, and in three of the four target-annotation-volume quartiles, Target Mean attains a higher *pair-level* ROC-AUC than OTRec for genetic, familial and congenital disease, 0.847 against 0.807, while ranking worse *within* each disease (MRR 0.497 against 0.594). A globally popular target list scores well when all pairs are pooled and poorly on making a shortlist of “top targets”.

**Table 4:** Disease-centric shortlist in the temporal test across 1,455 diseases with at least one target that acquired a clinical-progression label by 2025.

| Model | Hit@1 | Hit@5 | Hit@10 | MRR |
| --- | --- | --- | --- | --- |
| **OTRec** | **0.463** | **0.730** | **0.833** | **0.586** |
| OTTree (CatBoost) | 0.418 | 0.673 | 0.786 | 0.535 |
| Target Mean | 0.405 | 0.625 | 0.720 | 0.511 |
| Open Targets score | 0.263 | 0.416 | 0.484 | 0.342 |

### 4.3. Cold-Start and Feature Analyses

We stratified the temporal test set by each target’s number of clinical indications in the 2022 frame (Table 5). Of the 119 zero-indication targets, 117 are absent from the 2022 training frame altogether, so this subgroup is a genuine cold start. OTRec reaches ROC-AUC 0.887 on it, while Target Mean is at chance (0.505). Performance is weaker on the subgroup with only one clinical indication.

**Table 5:** Temporal test-target stratification by number of clinically associated indications in the 2022 frame. Positive prevalence in parentheses.

| 2022 target indications | Targets | Positives | ROC-AUC | PR-AUC |
| --- | --- | --- | --- | --- |
| 0 | 119 | 618 (1.2%) | 0.887 | 0.255 |
| Exactly 1 | 104 | 117 (0.5%) | 0.722 | 0.013 |
| $\geq 2$ | 1,298 | 15,339 (4.7%) | 0.869 | 0.293 |

Across four matched runs, adding ontology, GO/pathway, tractability, and constraint annotations had little effect on pooled performance (ROC-AUC 0.874 vs. 0.872; PR-AUC 0.285 vs. 0.292 for text-only and full-feature models, respectively), but improved performance on cold-start targets, increasing ROC-AUC from 0.881 to 0.888 and PR-AUC from 0.187 to 0.248. Cold-start PR-AUC improved in all four runs. Ablation suggests that GO/pathway and tractability annotations drive this gain (S3, Table S3). In a single-run ablation, removing the auxiliary OTP-score head did not reduce temporal-split performance (ROC-AUC 0.879, PR-AUC 0.293).

Because the temporal annotation strings come from a later shared snapshot, we also assessed sensitivity to annotation timing. The clinical tractability tokens (“Approved Drug”, “Advanced Clinical”, “Phase 1 Clinical”, “Clinical Precedence”) are the most direct inputs that might encode post-2022 outcomes. Removing them in a single matched rerun barely moved pooled ROC-AUC / PR-AUC from 0.877*/*0.286 to 0.870*/*0.282. On zero-indication targets ROC-AUC changed from 0.887 to 0.886 and PR-AUC from 0.255 to 0.201. Rebuilding all annotation text from Release 22.02 itself gives ROC-AUC 0.847*±*0.010 (four matched reruns) on the 58% of test pairs whose disease terms exist in the 2022 ontology, vs. 0.854 for the reference run.

### 4.4. Disease Repurposing Candidates

We identified 214,968 previously unlinked candidate associations with score *≥*0.65 for 4,347 diseases and 2,808 targets, including 2,322 target-orphan diseases, defined here as diseases with no clinically associated target in OTP. We generated candidates by screening the druggable genome, removing known links, and keeping up to 200 predictions per disease. The 0.65 threshold was set from the precision–recall curve on out-of-fold CV data (92% in-distribution precision, 62% recall). Candidate-bearing diseases carry a median of 14 candidates (interquartile range 2–78), and approximately 77% of the *∼*19,000 scored diseases have none above threshold; 1,614 of the 2,322 target-orphan diseases have more than one candidate (median 3).

Some targets are nominated in many diseases: for the median disease 2 of the top 10 candidates come from the 20 most frequently nominated targets. For example, GUCY1B2 and TUBB8 appear in 64.9% and 64.7% of candidate diseases. These two illustrate different mechanisms. OTP’s known-drug table maps the nitrovasodilator drug class to GUCY1B2 across 116 diseases in

Release 25.06, and 20 of its 21 label-changed temporal-test pairs became positive by 2025; TUBB8 carries no clinical record of its own but shares the annotation profile of *β*-tubulins such as TUBB1, a chemotherapy target with known-drug evidence across 445 diseases. Broadly tractable targets enter trials across many indications, which is why Target Mean is a strong baseline (Table 2) and why debiasing against it costs accuracy (S6; cf. Section 4.2); we therefore release the raw score together with a per-target nomination frequency. The temporal evaluation tests later clinical-label acquisition within the release-difference cohort, but it does not directly estimate precision for the final score-thresholded set of previously unlinked predictions. The ranked predictions should therefore not be treated as a validated discovery list. We annotated selected candidates for interestingness, novelty, and plausibility with the InterFeat framework [31]; the full list is in the repository (Appendix S1).

The candidates are browsable through an interactive Gradio web demo [32] (Figure 4) supporting both forward (disease *→* ranked targets) and reverse (target *→* ranked diseases) queries against the trained model, hosted at https://huggingface.co/spaces/GrimSqueaker/OTRec.

**Figure 4:**
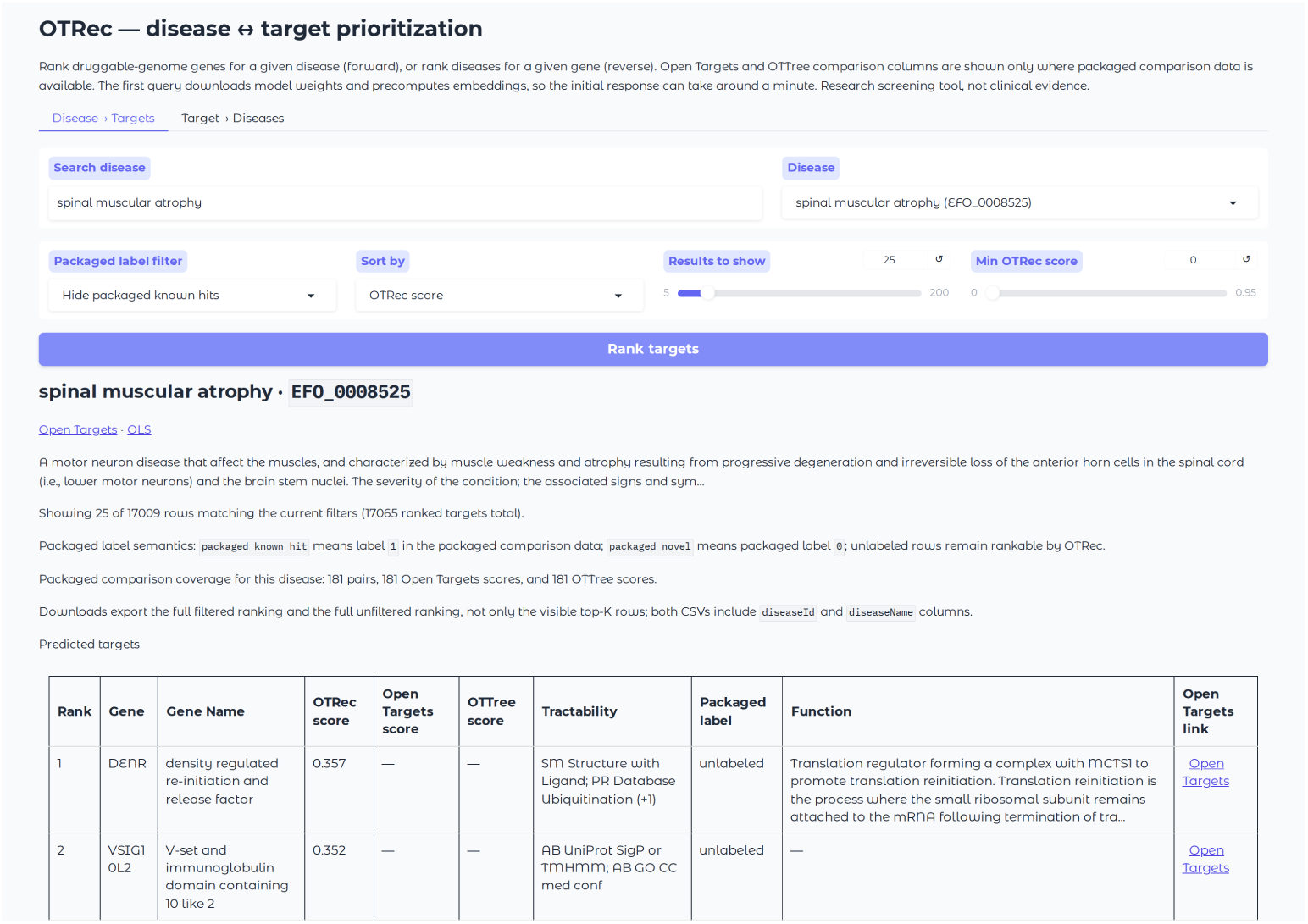
OTRec interactive demo. Users select a disease or target; the app returns the ranked candidate list with OTP cross-links, tractability annotations and druggable-genome ranking.

### 4.5. Illustrative Candidate Examples

The following three cases illustrate highly ranked associations. They were chosen for discussion rather than by a systematic screen, and none is experimental evidence or a treatment recommendation; they are intended to stimulate research and to frame hypotheses for experimental follow-up. *POLA1* / *POLA2* in central nervous system cancer (2022*→*2025). OTRec ranked the catalytic and accessory subunits of replicative DNA polymerase *α*— *POLA1* (score 0.961, rank 18 of 95) and *POLA2* (score 0.966, rank 8 of 95)— among the top candidates for central nervous system cancer (EFO_0000326), despite a near-zero OTP known-drug score in 2022 (*∼*0.06). Both pairs acquired clinical-progression labels by 2025. Glioma literature supports replication-stress dependencies [33, 34]; direct clinical validation of POLA1/POLA2 for this indication remains limited. POL*α* (POLA1) is over-expressed in glioblastoma relative to normal brain, and the POL*α* inhibitor ST1926 is cytotoxic to glioblastoma cell lines [35, 36]. In other tumor types, POLA1 loss of function is synthetically lethal with ATR and CHK1 inhibition [37, 38], and POLA1 depletion sensitizes BRCA1-deficient models to PARP inhibitors [39]; these combinations have not been tested for POLA1 in glioma.

*NPY2R* and *GIPR* in obesity (2022*→*2025). Neither gene appears anywhere in the 2022 training frame, and both carry a 2022 OTP overall association score of 0.000. OTRec nonetheless ranked *NPY2R* first and *GIPR* third among the 63 candidates for obesity (scores 0.681 and 0.613; OTTree 0.181 and 0.434), and both acquired clinical-progression labels by 2025. GIPR is one of the two receptors engaged by tirzepatide, whose phase 3 obesity trial reported in mid-2022, after the training snapshot [40]. GIPR was first characterized through its role in incretin-mediated insulin secretion [41, 42] and is now understood as a regulator of energy balance, adipose tissue biology and appetite [43, 44], though aspects of its pharmacology remain contested, with both receptor agonism and antagonism reported to reduce body weight [45, 46].

*FDPS* in bone chondrosarcoma. The highest-scoring candidate for bone chondrosarcoma (MONDO_0000515), which carries no clinically associated targets in the used release, is *FDPS* (score 0.944; rank 1 of 32). FDPS is the target of nitrogen-containing bisphosphonates [47], which reduce chondrosarcoma cell viability in vitro [48, 49] and slow tumor progression in rat models [50, 51]. Adjacent chondrosarcoma terms carry clinical annotations, so the no-known-target flag is per ontology term rather than per disease.

### 4.6. Properties of the Candidate Set

These are cohort-level properties of the released scores, not explanations of individual predictions. Predicted targets lacking approved drugs are less genetically constrained (*p <* 3.3 *×* 10*^−^*^19^, Cohen’s *d*=*−*0.37) and enriched for secreted proteins (+8.4%, *p <* 2.1 *×* 10*^−^*^7^), indicating divergent tractability profiles. They map to significantly fewer annotated pathways (*p <* 7.2 *×* 10*^−^*^50^), reflecting lower annotation coverage rather than functional irrelevance. Paralog counts and sequence identity do not differ significantly (*p*=0.63) at the cohort level, although within an individual disease the highest-ranked candidates are frequently drawn from a single gene family (quantified below). The model assigns higher scores to disease-specific targets (median 0.936) than to targets linked to many diseases (median 0.878; *p <* 6.3 *×* 10*^−^*^17^). Score magnitude and candidacy frequency are distinct: GUCY1B2, a target linked to many diseases, is nonetheless the most frequently nominated candidate overall (Section 4.4). OTRec scores track clinical progression, not just trial entry: across *∼*65,000 known positive associations, out-of-fold scores correlate with the maximum clinical trial phase (1–4) reached (Spearman *ρ*=0.20, weak but significant). All p-values reported in this section are Bonferroni-corrected for the six tests performed.

#### 4.6.1. Gene-Family Structure of the Predictions

Members of a gene family may receive similar top predictions, raising the question of whether this reflects shared biology or merely generic popularity of heavily studied diseases. Disease popularity alone sets a high baseline, with two random druggable genes sharing 14.7 of their top 30 predicted diseases. This is because a small set of intensively trialled indications dominates rankings. We compared, for each of the 471 HGNC gene groups [52] with at least ten members, the mean pairwise overlap of members’ top-30 disease lists against a null that draws random gene sets from the same lists, conditioning exactly on disease popularity [53]. Family coherence survives this control: 297 of 471 families (63%) share significantly more predicted diseases than the popularity null (BH *q<*0.05), whereas an arbitrary grouping used as a negative control shows no excess. This mirrors the label space itself: clinical development recurrently targets whole families and complexes [54] and the family signal is prospective, not only descriptive: for known clinical pairs, the same disease appears in a sibling’s top-30 list 1.23*×* more often than popularity predicts, with larger lifts for protein complexes such as the ribosome.

As a case study we examine the RAB GTPases, 64 members (composed of RABs and RAB binding proteins), none with clinical evidence in the release used, whose members share similar candidate lists. The family is significantly coherent (+3.7 excess shared diseases per pair, *q*=5.8*×*10*^−^*^4^), with many shared oncological candidates, including specific carcinomas that are only moderately popular genome-wide. This reinforces that family-coherent shortlists are expected and partly informative, but within-family ordering carries little extra signal. This analysis used the deployed webapp model, which is retrained on Open Targets 25.12 OTRec (held-out ROC-AUC 0.947, PR-AUC 0.824); per-family statistics are included (S9).

## 5. Discussion

OTRec’s advantage is concentrated in curator-friendly ranked shortlists for each disease, and on targets without prior clinical history, where retrospective evidence scores have little to aggregate. The comparison with Target Mean separates two kinds of signal: global target popularity, which is strong when all pairs are pooled, and disease-based ranking, where the learned representations lead (Section 4.2).

The current study is limited by target scope, label noise, feature timing and temporal coverage. We use the Finan et al. [19] druggable genome and OTP-defined diseases; non-coding targets, genes outside this set, and diseases absent from OTP are not ranked by default. OTRec ranks gene targets independently and does not model contextual factors such as direction of modulation, therapeutic modality, tissue context, dose, safety or off-target liabilities; downstream polypharmacology and selection remain the researcher’s responsibility. The temporal labels are release-separated (2022 vs. 2025), while the annotation text comes from one shared later snapshot; matched reruns quantify this effect as minor (Section 4.3).

The 2022*→*2025 evaluation uses one release pair. We report variance across five runs, but rolling release-pair evaluations (e.g., 2021*→*2024) could more fully test the stability of the prospective predictions. Earlier OTP releases are sparser and their formats change, which limits how far back we can go. OTRec uses text vectorization rather than a pretrained language model tower. This trades representational richness for vocabulary adaptation and inference scale (*∼*87M pairs at prediction time). A frozen BioClinical ModernBERT model improves substantially over TF-IDF but remained below OTRec and OTTree in target-disjoint CV. Stronger end-to-end encoder adaptation remains an open direction. However, the underlying language model was trained on information from the 2025 test period, so it is not a fair comparison. All non-clinical pairs are treated as implicit negatives *y*=0; the temporal relabeling does not solve positive–unlabeled learning. Sweeping the matched OTTree baseline did not change the model ordering (Appendix S5).

In terms of limitations, OTRec may surface hypotheses for diseases, but it can also reproduce the bias of public databases towards well-studied diseases, populations and broadly tractable targets, which is why we release nomination frequency alongside the scores. We present it as a screening and prioritization aid, not as a substitute for biological validation, clinical evidence, safety assessment, or expert review.

## 6. Conclusion

Ranking druggable targets across OTP diseases against later clinical evidence is a tractable, precisely defined surrogate for prospective therapeutic relevance. Learned representations from text and ontology improve over the retrospective evidence score in target-disjoint CV, in temporal disease-level ranking, and on targets with no prior clinical indication, while pooled temporal PR-AUC does not beat a strong historical target prior, possibly due to how medical research is focused. We release the previously unlinked candidate associations, an interactive interface, and the benchmark splits and analyses.

## Data availability

This study uses public data. Open Targets Platform data are released under Creative Commons Zero (CC0 1.0) public-domain; other evidence sources retain their own licenses. Code, trained models, ranked predictions, the benchmark splits, and the interactive query interface are freely available under the MIT license at https://github.com/LinialLab/OTRec. The interactive platform is hosted on Hugging Face Spaces at https://huggingface.co/ spaces/GrimSqueaker/OTRec.

## CRediT authorship contribution statement

**Dan Ofer:** Conceptualization, Methodology, code, Data curation, analysis, Writing.

**Michal Linial:** Supervision, Writing.

