## Supplemental data S3 for "OTRec: A Deep Learning Recommender for Druggable Disease–Target Prioritization"

### output-EDA-Analysis-Novel_Predictions

December 18, 2025

### 1 Disease–Target Prediction Analysis and Novelty Detection

This notebook provides an end-to-end workflow for exploring a dataset of disease–gene predictions generated by a machine-learning model trained on OpenTargets features. It includes functions for loading and cleaning the data, computing descriptive statistics, visualising distributions and counts, identifying novel predictions with minimal evidence, filtering out promiscuous genes or diseases, and performing anomaly detection via **IsolationForest**. The analysis is designed to surface interpretable insights for drug-repurposing and novelty detection research.

To run this notebook, place the compressed CSV file (`positive_preds_train_and_novelIndirect.csv.gz`) in the working directory. All required libraries (pandas, numpy, matplotlib, seaborn, scikit-learn) should be installed in your environment.

- Warning: current candidates data missing things like early onset alzheimers disease?

[1]:
```
import os
import numpy as np
import pandas as pd
import matplotlib.pyplot as plt
import seaborn as sns
from sklearn.ensemble import IsolationForest
from sklearn.preprocessing import StandardScaler

pd.set_option('display.precision',3)
pd.set_option('display.max_columns', 30)
```

[2]:
```
DROP_CANCER = True
```

#### 1.1 Helper functions

[3]:
```
def load_data(filepath: str,fillnan_val=-1) -> pd.DataFrame:
    '''Load the compressed CSV file and prepare the dataframe.

    Parameters
    ----------
    filepath : str
        Path to the gzipped CSV file.
```

```
    Returns
    -------
    pd.DataFrame
        DataFrame with NaNs filled and numeric columns converted where␣
↪appropriate.
    '''
    if filepath.endswith('.gz'):
        df = pd.read_csv(filepath, compression='gzip')
    else:
        df = pd.read_csv(filepath)
    print(f"replacing nan label with {fillnan_val}")
    df['label'] = df['label'].fillna(fillnan_val)
    score_cols = ['score_direct', 'score_known_drug', 'score_literature',
                  'score_affected_pathway', 'score_genetic_association',
                  'score_rna_expression', 'score_somatic_mutation']
    numeric_cols = ['pred', 'num_known_targets', 'in_aba'] + score_cols
    for col in numeric_cols:
        df[col] = pd.to_numeric(df[col], errors='coerce')
    df[numeric_cols] = df[numeric_cols].fillna(0)
    return df


def summarise_data(df: pd.DataFrame) -> None:
    '''Print summary statistics for the dataset.'''
    print(f"Dataset shape: {df.shape}")
    label_counts = df['label'].value_counts(dropna=False).sort_index()
    print("Label counts (after filling NaN->-1):")
    print(label_counts)
    pred_counts = df['pred_label'].value_counts(dropna=False).sort_index()
    print("Predicted label counts:")
    print(pred_counts)
    score_cols = ['score_direct', 'score_known_drug', 'score_literature',
                  'score_affected_pathway', 'score_genetic_association',
                  'score_rna_expression', 'score_somatic_mutation']
    # evidence_means = df.groupby('label_filled')[score_cols].mean().round(3)
    evidence_means = df.select_dtypes("number").groupby('label').mean().
↪round(3) # include more
    print("Mean evidence scores by label:")
    display(evidence_means)


def plot_pred_distribution(df: pd.DataFrame) -> None:
    '''Plot the distribution of predicted probabilities by original label group.
↪'''
    plt.figure(figsize=(7,4))
    groups = sorted(df['label'].unique())
    colors = ['blue','orange','green','red']
```

```
    for i, label in enumerate(groups):
        subset = df[df['label'] == label]
        plt.hist(subset['pred'], bins=10, alpha=0.5, density=True,
                 label=f'label={label}', color=colors[i % len(colors)])
    plt.title('Distribution of predicted probability by label group')
    plt.xlabel('Predicted probability')
    plt.ylabel('Density')
    plt.legend()
    plt.tight_layout()
    plt.show()


def compute_gene_disease_counts(df: pd.DataFrame, subset: str = 'unknown') ->␣
 ↪tuple:
    '''Compute counts of predicted associations per gene and per disease for a␣
 ↪subset.

    Parameters
    ----------
    df : pd.DataFrame
        Input DataFrame with predictions.
    subset : str
        Which subset to consider: 'unknown', 'positive', 'negative', or 'all'.

    Returns
    -------
    tuple
        Two Series: counts per gene and counts per disease.
    '''
    if subset == 'unknown':
        data = df[df['label'] == -1]
    elif subset == 'positive':
        data = df[df['label'] == 1]
    elif subset == 'negative':
        data = df[df['label'] == 0]
    else:
        data = df
    gene_counts = data['approvedSymbol'].value_counts()
    disease_counts = data['disease_name'].value_counts()
    return gene_counts, disease_counts


def plot_top_counts(counts: pd.Series, title: str, n: int = 10) -> None:
    '''Plot a bar chart of the top n counts.'''
    top_counts = counts.head(n)
    plt.figure(figsize=(8,4))
    top_counts.plot(kind='bar')
```

```
    plt.title(title)
    plt.xlabel('Name')
    plt.ylabel('Count')
    plt.xticks(rotation=45, ha='right')
    plt.tight_layout()
    plt.show()


def compute_novel_predictions(df: pd.DataFrame) -> pd.DataFrame:
    '''Identify novel predictions and compute evidence gaps.'''
    score_cols = ['score_direct', 'score_known_drug', 'score_literature',
                  'score_affected_pathway', 'score_genetic_association',
                  'score_rna_expression', 'score_somatic_mutation']
    novel = df[df['label'] == -1].copy()
    novel['evidence_mean'] = novel[score_cols].mean(axis=1)
    novel['pred_minus_evidence'] = novel['pred'] - novel['evidence_mean']
    novel_sorted = novel.sort_values('pred_minus_evidence', ascending=False)
    return novel_sorted


def filter_specific_targets(df: pd.DataFrame, max_gene_count: int = 100,␣
 ↪max_disease_count: int = 500) -> pd.DataFrame:
    '''Filter predictions to those where gene and disease occur fewer than␣
 ↪specified times.'''
    gene_counts = df['approvedSymbol'].value_counts()
    disease_counts = df['disease_name'].value_counts()
    allowed_genes = gene_counts[gene_counts <= max_gene_count].index
    allowed_diseases = disease_counts[disease_counts <= max_disease_count].index
    filtered = df[(df['approvedSymbol'].isin(allowed_genes)) &
                  (df['disease_name'].isin(allowed_diseases))].copy()
    return filtered


def run_isolation_forest(df: pd.DataFrame, contamination: float = 0.05,␣
 ↪random_state: int = 42) -> pd.DataFrame:
    '''Compute anomaly scores for predictions using IsolationForest.'''
    score_cols = ['score_direct', 'score_known_drug', 'score_literature',
                  'score_affected_pathway', 'score_genetic_association',
                  'score_rna_expression', 'score_somatic_mutation']
    num_cols = ['pred', 'num_known_targets', 'in_aba'] + score_cols
    X = df[num_cols].astype(float)
    scaler = StandardScaler()
    X_scaled = scaler.fit_transform(X)
    iso = IsolationForest(n_estimators=200,␣
 ↪contamination=contamination,n_jobs=-1,max_samples=8_000,
                          random_state=random_state)
    iso.fit(X_scaled)
```

```
    anomaly_scores = -iso.decision_function(X_scaled)
    outliers = iso.predict(X_scaled)
    df = df.copy()
    df['anomaly_score'] = anomaly_scores
    df['is_outlier'] = (outliers == -1)
    return df
```

[4]:
```
import numpy as np
import pandas as pd
from typing import Literal, Optional

def shortlist_specific_pairs(
    df: pd.DataFrame,
    gene_col: str = "approvedSymbol",
    disease_col: str = "disease_name",
    pred_col: str = "pred",
    label_col: Optional[str] = "label",
    pred_min: float = 0.75,
    hub_quantile: float = 0.99,
    score: Literal["tfidf", "npmi"] = "tfidf",
    top_k_per_gene: int = 3,
) -> pd.DataFrame:
    """
    Short, explainable pipeline to surface *specific* (not hub-driven) disease-
target pairs.

    Steps
    -----
    1) Focus: keep novel/unknown pairs and high-confidence predictions: label␣
 ↪is NA (if present) and pred >= pred_min.
    2) Hub filter: drop top `hub_quantile` by degree for genes and diseases in␣
 ↪this working slice.
    3) Specificity scoring:
       - "tfidf": tf(g→d) * idf(d), where tf is within-gene share and idf␣
 ↪penalizes globally common diseases.
       - "npmi": normalized pointwise mutual information, highlighting␣
 ↪co-occurrence beyond independence.
    4) Shortlist: return top-K diseases per gene by chosen score, with helpful␣
 ↪context columns.

    Returns
    -------
    DataFrame with columns:
      [gene_col, disease_col, 'cnt', 'gene_total', 'dis_deg', 'spec_score',␣
 ↪'npmi', 'pred_mean']
```

```
    Notes
    -----
    - Keep the method blurb in the paper to ~2-3 sentences: degree-based hub␣
↪removal + TF-IDF (or nPMI) specificity.
    - All operations are on the *working slice* (unknown & high-confidence), so␣
↪the shortlist stays actionable.
    """
    work = df.copy()

    # 1) Focus on novel + high-confidence
    if label_col in work.columns and work[label_col].notna().any():
        novel_mask = work[label_col].isna()
    else:
        novel_mask = np.ones(len(work), dtype=bool)  # no labels provided →␣
↪treat all as candidate
    conf_mask = work[pred_col] >= pred_min
    work = work[novel_mask & conf_mask].copy()
    if work.empty:
        return work.assign(spec_score=np.nan, npmi=np.nan)

    # 2) Degree-based hub filter (simple & explainable)
    deg_gene = work.groupby(gene_col)[disease_col].nunique()
    deg_dis  = work.groupby(disease_col)[gene_col].nunique()
    g_cut = deg_gene.quantile(hub_quantile)
    d_cut = deg_dis.quantile(hub_quantile)
    keep_genes = deg_gene[deg_gene < g_cut].index
    keep_dis   = deg_dis[deg_dis   < d_cut].index
    work = work[work[gene_col].isin(keep_genes) & work[disease_col].
↪isin(keep_dis)].copy()
    if work.empty:
        return work.assign(spec_score=np.nan, npmi=np.nan)

    # 3) Specificity scores
    pairs = (work.groupby([gene_col, disease_col]).size()
                 .rename("cnt").reset_index())

    gene_total = pairs.groupby(gene_col)["cnt"].sum().rename("gene_total")
    dis_deg    = pairs.groupby(disease_col)[gene_col].nunique().
↪rename("dis_deg")
    pairs = pairs.merge(gene_total, on=gene_col).merge(dis_deg, on=disease_col)
    N_pairs = float(pairs["cnt"].sum())

    # TF-IDF-like specificity (explainable, one-liner)
    tf  = pairs["cnt"] / pairs["gene_total"].clip(lower=1)
    idf = np.log((N_pairs + 1.0) / (pairs["dis_deg"] + 1.0))  # global rarity␣
↪of the disease
```

```
    spec_score = tf * idf
    pairs["spec_score"] = spec_score

    # nPMI (chance-corrected alternative; useful as a sensitivity check)
    # add-one smoothing keeps it stable for small counts
    p_xy = (pairs["cnt"] + 1.0) / (N_pairs + 1.0)
    # p_x: per-gene mass; p_y: per-disease mass
    # gene_total already computed, need disease totals
    dis_total = pairs.groupby(disease_col)["cnt"].sum().rename("dis_total")
    pairs = pairs.merge(dis_total, on=disease_col)
    p_x = (pairs["gene_total"] + 1.0) / (N_pairs + 1.0)
    p_y = (pairs["dis_total"] + 1.0) / (N_pairs + 1.0)
    PMI  = np.log(p_xy / (p_x * p_y))
    pairs["npmi"] = PMI / (-np.log(p_xy))

    # Optionally attach mean predicted probability for quick triage
    pred_mean = (work[[gene_col, disease_col, pred_col]]
                 .groupby([gene_col, disease_col])[pred_col].mean()
                 .rename("pred_mean"))
    pairs = pairs.merge(pred_mean, on=[gene_col, disease_col], how="left")

    # 4) Shortlist: rank by requested score and keep top-K diseases per gene
    key_score = "spec_score" if score == "tfidf" else "npmi"
    out = (pairs.sort_values(key_score, ascending=False)
                 .groupby(gene_col)
                 .head(top_k_per_gene)
                 .reset_index(drop=True))

    cols = [gene_col, disease_col, "cnt", "gene_total", "dis_deg",␣
 ↪"spec_score", "npmi", "pred_mean"]
    return out[cols].sort_values([gene_col, key_score], ascending=[True, False])
```

#### 1.2 Load the dataset

[5]:
```
# file_path = 'positive_preds_train_and_novelIndirect.csv.gz' ## ORIG, from␣
 ↪catboost CV
file_path ="DL_novel_candidates_predictions.csv" ## alt - from DL (diff number␣
 ↪cases, no CV, predicted top for all data inc. train + inc known positives
```

[6]:
```
df = pd.read_csv(file_path)
## compat workrarounds for new dl pred, to work with code for old preds
df["Gene_name"] = df["targetSymbol"]
df.rename(columns={"diseaseName":"disease_name","targetSymbol":"approvedSymbol",
                   # "label":"pred_label",
                   "disease_num_known_clinical_targets":
 ↪"num_known_targets"},inplace=True,errors="ignore")
```

```
# df["label"] =1
df["pred_label"] =1
df["pred"] =1
df
```

[6]:

```
             diseaseId     disease_name          targetId approvedSymbol  \
0       UBERON_0000104       life cycle  ENSG00000123201        GUCY1B2
1       UBERON_0000104       life cycle  ENSG00000261456          TUBB8
2       UBERON_0000104       life cycle  ENSG00000088832         FKBP1A
3       UBERON_0000104       life cycle  ENSG00000176014          TUBB6
4       UBERON_0000104       life cycle  ENSG00000173213         TUBB8B
…                    …                …                …              …
147228      DOID_10113  trypanosomiasis  ENSG00000151834         GABRA2
147229      DOID_10113  trypanosomiasis  ENSG00000163285         GABRG1
147230      DOID_10113  trypanosomiasis  ENSG00000182256         GABRG3
147231      DOID_10113  trypanosomiasis  ENSG00000183454         GRIN2A
147232      DOID_10113  trypanosomiasis  ENSG00000186297         GABRA5

        score  label            source  num_known_targets  orphan Gene_name  \
0       0.948     -1  model_prediction                  0    True   GUCY1B2
1       0.824     -1  model_prediction                  0    True     TUBB8
2       0.734     -1  model_prediction                  0    True    FKBP1A
3       0.710     -1  model_prediction                  0    True     TUBB6
4       0.700     -1  model_prediction                  0    True    TUBB8B
…         …    …                 …                  …       …         …
147228  1.000      1    known_positive                 11   False    GABRA2
147229  1.000      1    known_positive                 11   False    GABRG1
147230  1.000      1    known_positive                 11   False    GABRG3
147231  1.000      1    known_positive                 11   False    GRIN2A
147232  1.000      1    known_positive                 11   False    GABRA5

        pred_label  pred
0                1     1
1                1     1
2                1     1
3                1     1
4                1     1
…                …     …
147228           1     1
147229           1     1
147230           1     1
147231           1     1
147232           1     1

[147233 rows x 12 columns]
```

[7]:
```
# df = load_data(file_path,fillnan_val=0)
# # display(df.loc[df["disease_name"].str.contains("alz",case=False,na=False)].
 ↪drop_duplicates("disease_name"))
df = df.round(4)

print(df.shape[0])
if DROP_CANCER:
    print("Drop cancer diseases") # lots of them and they'r ediff
    df = df.loc[~df["disease_name"].str.
 ↪contains("cancer|carcinom|neoplas|tumor|lymphoma|leukemia|gliob",case=False,na=False)]
    print(df.shape[0])

try:
    ## add features: NOTE: TODO: Add these in advance from the FULL Dataset.␣
 ↪This is just a proxy hack!!!
    if "disease_max_drug_score" not in df.columns:
        df["disease_max_drug_score"] = df.
 ↪groupby(["diseaseId"])["score_known_drug"].transform("max")
    if "target_max_drug_score_overall" not in df.columns:
        df["target_max_drug_score"] = df.
 ↪groupby(["Gene_name"])["score_known_drug"].transform("max")
except Exception as e:
    print(e)
summarise_data(df)
```

```
147233
Drop cancer diseases
97332
'Column not found: score_known_drug'
Dataset shape: (97332, 12)
Label counts (after filling NaN->-1):
label
-1    58104
 1    39228
Name: count, dtype: int64
Predicted label counts:
pred_label
1    97332
Name: count, dtype: int64
Mean evidence scores by label:

       score  num_known_targets  pred_label  pred
label
-1     0.802             15.557         1.0   1.0
 1     1.000             98.613         1.0   1.0
```

[8]:
```
df.nunique()
```

[8]:
```
diseaseId           5854
disease_name        5853
targetId            1806
approvedSymbol      1804
score                390
label                  2
source                 2
num_known_targets    165
orphan                 2
Gene_name           1804
pred_label             1
pred                   1
dtype: int64
```

##### 1.2.1 cancer-disease filter

[9]:
```
### doesn't filter anything, maybe due to lack of annot innovel candidates␣
↪being predicted on.
##### we 'll stick to the disease name filter heursitic'
# if DROP_CANCER:
#     disease_df = pd.read_parquet('../data/opentargets/
↪disease',columns=['id','name','therapeuticAreas']).dropna(axis=1,how="all")
#     ta_map = disease_df.set_index('id')['name'].to_dict()
#     labels_to_remove = ['measurement', 'phenotype',
#                          # 'biological process',
#                          'cell proliferation disorder']
#     exploded_tas = disease_df[['id', 'name','therapeuticAreas']].
↪explode('therapeuticAreas').dropna()
#     exploded_tas['label'] = exploded_tas['therapeuticAreas'].map(ta_map)
#     exploded_tas = exploded_tas[exploded_tas['label'].isin(labels_to_remove)].
↪reset_index(drop=True)
#     ids_to_remove = exploded_tas['id'].unique()
#     names_to_remove = exploded_tas['name']#.unique()
#     print(df.shape[0])
    # df = df.loc[~df['diseaseId'].isin(ids_to_remove)]
#     print(df.shape[0])
```

[10]:
```
# if DROP_CANCER:
# # [6] CORRECTED: cancer-disease filter
#     # Load the disease data
#     disease_df = pd.read_parquet(
#         '../data/opentargets/disease',
#         columns=['id', 'name', 'therapeuticAreas']
#     ).dropna(subset=['id', 'name'])

#     # --- START OF FIX ---
```

```
#     # 1. Create a map from NAME to ID
#     #    (The reverse of your original ta_map)
#     name_to_id_map = disease_df.set_index('name')['id'].to_dict()

#     # 2. Your list of names to remove
#     labels_to_remove = [
#         'measurement', 'phenotype', 'biological process',
#         'cell proliferation disorder'
#     ]

#     # 3. Convert those names to their corresponding IDs (e.g., 'phenotype' ->␣
↪'EFO_0005541')
#     label_ids_to_remove = set(
#         name_to_id_map[name] for name in labels_to_remove if name in␣
↪name_to_id_map
#     )

#     # 4. Explode the therapeuticAreas
#     exploded_tas = disease_df[['id', 'therapeuticAreas']].
↪explode('therapeuticAreas').dropna()

#     # 5. Find rows where the 'therapeuticAreas' ID is one of the label IDs to␣
↪remove
#     matching_rows = exploded_tas[
#         exploded_tas['therapeuticAreas'].isin(label_ids_to_remove)
#     ]

#     # 6. The 'id' column of these rows is your correct list of diseases to␣
↪remove
#     #    This list will now contain EFO/MONDO IDs that match your␣
↪df['diseaseId']
#     ids_to_remove = matching_rows['id'].unique()

#     # --- END OF FIX ---

#     # Now, your original filter logic will work
#     print(f"Original df shape: {df.shape[0]}")
#     print(f"Found {len(ids_to_remove)} disease IDs to remove.")

#     df_filtered = df.loc[~df['diseaseId'].isin(ids_to_remove)]

#     print(f"New filtered df shape: {df_filtered.shape[0]}")

#     # You can check which rows *would* be removed by running this:
# df.loc[df['diseaseId'].isin(ids_to_remove)]
```

**target prioritisation eda**

- Compare known targets, predicted targets, and non-predicted targets
- Get diseases only level + predicted labels:

[11]:
```
df
```

[11]:
```
             diseaseId    disease_name         targetId approvedSymbol  \
0       UBERON_0000104      life cycle  ENSG00000123201        GUCY1B2
1       UBERON_0000104      life cycle  ENSG00000261456          TUBB8
2       UBERON_0000104      life cycle  ENSG00000088832         FKBP1A
3       UBERON_0000104      life cycle  ENSG00000176014          TUBB6
4       UBERON_0000104      life cycle  ENSG00000173213         TUBB8B
…                  …               …               …             …
147228      DOID_10113  trypanosomiasis  ENSG00000151834         GABRA2
147229      DOID_10113  trypanosomiasis  ENSG00000163285         GABRG1
147230      DOID_10113  trypanosomiasis  ENSG00000182256         GABRG3
147231      DOID_10113  trypanosomiasis  ENSG00000183454         GRIN2A
147232      DOID_10113  trypanosomiasis  ENSG00000186297         GABRA5

        score  label            source  num_known_targets  orphan Gene_name  \
0       0.948     -1  model_prediction                  0    True   GUCY1B2
1       0.824     -1  model_prediction                  0    True     TUBB8
2       0.734     -1  model_prediction                  0    True    FKBP1A
3       0.710     -1  model_prediction                  0    True     TUBB6
4       0.700     -1  model_prediction                  0    True    TUBB8B
…         …    …               …                …      …       …
147228  1.000      1    known_positive                 11   False    GABRA2
147229  1.000      1    known_positive                 11   False    GABRG1
147230  1.000      1    known_positive                 11   False    GABRG3
147231  1.000      1    known_positive                 11   False    GRIN2A
147232  1.000      1    known_positive                 11   False    GABRA5

        pred_label  pred
0                1     1
1                1     1
2                1     1
3                1     1
4                1     1
…              …    …
147228           1     1
147229           1     1
147230           1     1
147231           1     1
147232           1     1

[97332 rows x 12 columns]
```

[12]:
```
# df_dis = df.groupby(["disease_name","diseaseId"]).max().
↪select_dtypes("number").reset_index()
# df_dis

df_targ = df.groupby(["approvedSymbol","Gene_name","targetId"]).max().
↪select_dtypes("number").reset_index()
df_targ

target_priority = pd.read_parquet("../data/opentargets/target_prioritisation").
↪dropna(axis=0,how="all")
target_priority
```

```
---------------------------------------------------------------------------
FileNotFoundError                         Traceback (most recent call last)
Cell In[12], line 7
      4 df_targ = df.groupby(["approvedSymbol","Gene_name","targetId"]).max().
 ↪select_dtypes("number").reset_index()
      5 df_targ
----> 7 target_priority = pd.read_parquet("../data/opentargets/
 ↪target_prioritisation").dropna(axis=0,how="all")
      8 target_priority

File ~/anaconda3/lib/python3.13/site-packages/pandas/io/parquet.py:667, in␣
 ↪read_parquet(path, engine, columns, storage_options, use_nullable_dtypes,␣
 ↪dtype_backend, filesystem, filters, **kwargs)
    664     use_nullable_dtypes = False
    665 check_dtype_backend(dtype_backend)
--> 667 return impl.read(
    668     path,
    669     columns=columns,
    670     filters=filters,
    671     storage_options=storage_options,
    672     use_nullable_dtypes=use_nullable_dtypes,
    673     dtype_backend=dtype_backend,
    674     filesystem=filesystem,
    675     **kwargs,
    676 )

File ~/anaconda3/lib/python3.13/site-packages/pandas/io/parquet.py:267, in␣
 ↪PyArrowImpl.read(self, path, columns, filters, use_nullable_dtypes,␣
 ↪dtype_backend, storage_options, filesystem, **kwargs)
    264 if manager == "array":
    265     to_pandas_kwargs["split_blocks"] = True  # type: ignore[assignment]
--> 267 path_or_handle, handles, filesystem = _get_path_or_handle(
    268     path,
    269     filesystem,
    270     storage_options=storage_options,
```

```
    271     mode="rb",
    272 )
    273 try:
    274     pa_table = self.api.parquet.read_table(
    275         path_or_handle,
    276         columns=columns,
   (…)
    279         **kwargs,
    280     )

File ~/anaconda3/lib/python3.13/site-packages/pandas/io/parquet.py:140, in␣
 ↪_get_path_or_handle(path, fs, storage_options, mode, is_dir)
    130 handles = None
    131 if (
    132     not fs
    133     and not is_dir
   (…)
    138     # fsspec resources can also point to directories
    139     # this branch is used for example when reading from non-fsspec URLs
--> 140     handles = get_handle(
    141         path_or_handle, mode, is_text=False,␣
 ↪storage_options=storage_options
    142     )
    143     fs = None
    144     path_or_handle = handles.handle

File ~/anaconda3/lib/python3.13/site-packages/pandas/io/common.py:882, in␣
 ↪get_handle(path_or_buf, mode, encoding, compression, memory_map, is_text,␣
 ↪errors, storage_options)
    873         handle = open(
    874             handle,
    875             ioargs.mode,
   (…)
    878             newline="",
    879         )
    880     else:
    881         # Binary mode
--> 882         handle = open(handle, ioargs.mode)
    883     handles.append(handle)
    885 # Convert BytesIO or file objects passed with an encoding

FileNotFoundError: [Errno 2] No such file or directory: '../data/opentargets/
 ↪target_prioritisation'
```

[ ]:
```python
df_targ = df_targ.merge(target_priority,on="targetId",how="outer")
df_targ["label"] = df_targ["label"].fillna(-1)
```

[ ]:
```
# df_targ.select_dtypes("number").groupby("label").mean().plot(kind='line',␣
↪title='Mean Value by Category')
df_targ.select_dtypes("number").groupby("label").mean()
```

[ ]:
```
# df_targ.select_dtypes("number").groupby("label").mean().to_clipboard()
```

[ ]:
```
df.select_dtypes("number").groupby(['pred_label',"label"]).mean().round(3)
```

- 0 known `num_known_targets` (disease ? targets): leaves just label 0 (only a subset of label 0 a tthat!) and "missing label"/-1 samples.

This is as expected!

[ ]:
```
df.query("num_known_targets<1").label.value_counts(dropna=False)
```

[ ]:
```
# df.query("num_known_targets<?1")
```

**Nice example pred:** `disease:` pathological gambling | `target/gene:` HTR2C 5-hydroxytryptamine receptor 2C - that's a serotonin receptor (with known drugs) - `5-HT2C receptors mediate the release and increase of extracellular dopamine`

[ ]:
```
df.nunique()
```

[ ]:
```
df.loc[df["disease_name"].str.contains("anemia")]["label"].value_counts()
```

[ ]:
```
df
```

#### 1.3 Distribution of predicted probabilities

[ ]:
```
plot_pred_distribution(df)
```

#### 1.4 Gene and disease coverage in novel predictions

[13]:
```
novel_gene_counts, novel_disease_counts = compute_gene_disease_counts(df.
↪query("label<1"), subset='all')
print("Top genes with most novel associations:")
print(novel_gene_counts.head(10))
print("Top diseases with most novel associations:")
print(novel_disease_counts.head(10))

plot_top_counts(novel_gene_counts, 'Top genes with most novel associations',␣
↪n=15)
plot_top_counts(novel_disease_counts, 'Top diseases with most novel␣
↪associations', n=15)
```

```
Top genes with most novel associations:
approvedSymbol
```

```
GUCY1B2    4685
TUBB8      3016
FKBP1A     1759
TUBB8B     1678
TUBB6      1510
TUBB1      1282
GABRG3     1124
GABRA2     1009
GABRR3      946
GABRA5      910
Name: count, dtype: int64
Top diseases with most novel associations:
disease_name
amenorrhea                                        26
communicating hydrocephalus                       25
alcohol abuse                                     25
depressive disorder                               25
melanoma, cutaneous malignant, susceptibility to, 8    25
Richter syndrome                                  25
cheilitis                                         25
pancreas sarcoma                                  25
calcinosis                                        25
optic nerve disorder                              25
Name: count, dtype: int64
```

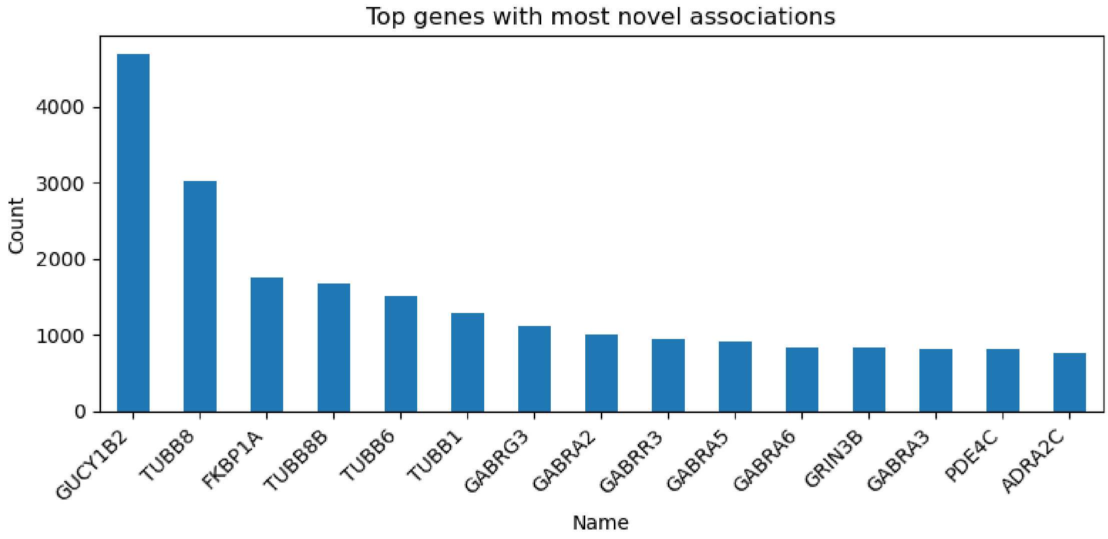

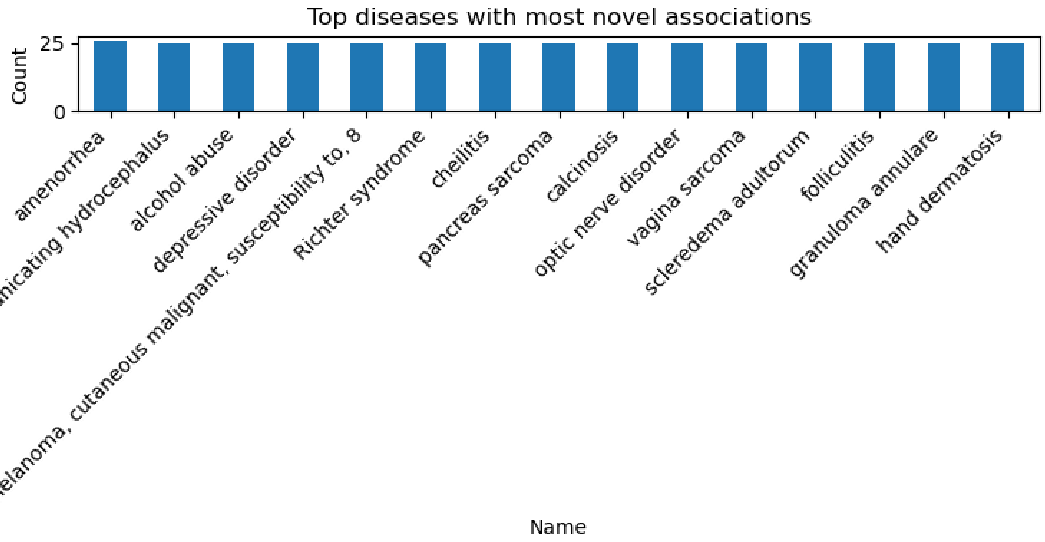


[14]:
```
display(novel_gene_counts.describe().round(1))
novel_gene_counts.hist()
```

```
count    1122.0
mean       51.8
std       223.7
min         1.0
25%         1.0
50%         5.0
75%        18.0
max      4685.0
Name: count, dtype: float64
```

[14]: <Axes: >

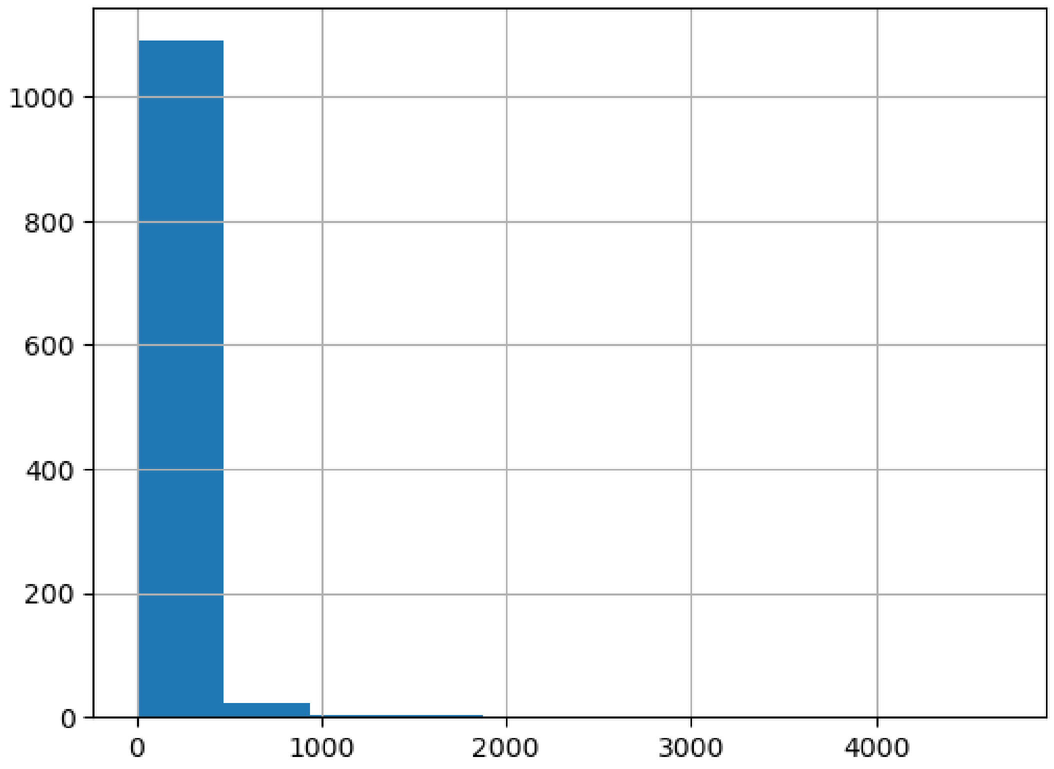


Get results for Orphan genes or diseases with 0~1 known associations (each) ###### Orphan diseases?

[19]:
```
df.query("num_known_targets<1")["disease_name"].nunique()
```

[19]: 4082

[20]:
```
orphan_novel_gene_counts, orphan_novel_disease_counts =␣
↪compute_gene_disease_counts(df.query("num_known_targets<1"), subset='all')
# orphan_novel_gene_counts, orphan_novel_disease_counts =␣
↪compute_gene_disease_counts(df, subset='unknown')

print("Orphan diseases with most novel associations:")
print(orphan_novel_disease_counts.head(10))

print("Genes with most predicted associations with orphan diseases:")
print(orphan_novel_gene_counts.head(10))

plot_top_counts(orphan_novel_gene_counts, 'Genes with most novel associations␣
↪in Orphan diseases', n=10)
plot_top_counts(orphan_novel_disease_counts, 'Orphan diseases with most novel␣
↪associations', n=10)
```

```
Orphan diseases with most novel associations:
disease_name
trichomoniasis                          25
acute laryngitis                        25
carotid artery dissection               25
chronic ethmoidal sinusitis             25
herpes simplex dermatitis               25
progressive bulbar palsy                25
heart transplant rejection              25
periostitis                             25
esophagus squamous cell papilloma       25
chronic eosinophilic pneumonia          25
Name: count, dtype: int64
Genes with most predicted associations with orphan diseases:
approvedSymbol
GUCY1B2     3737
TUBB8       2529
FKBP1A      1307
TUBB8B      1306
TUBB6       1184
TUBB1        990
GABRG3       590
GABRA2       514
TUBB2A       504
TUBB2B       501
Name: count, dtype: int64
```

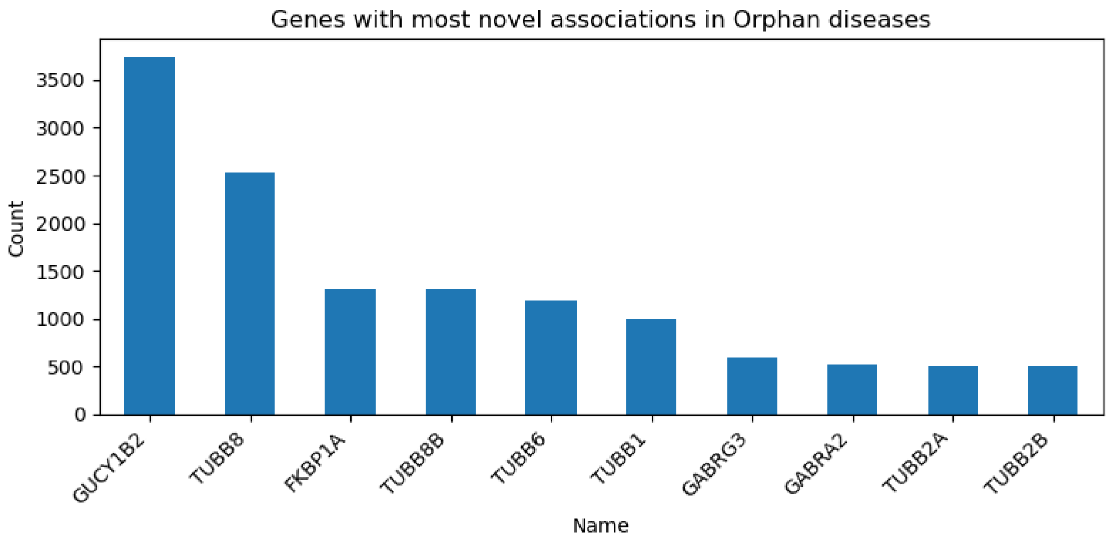

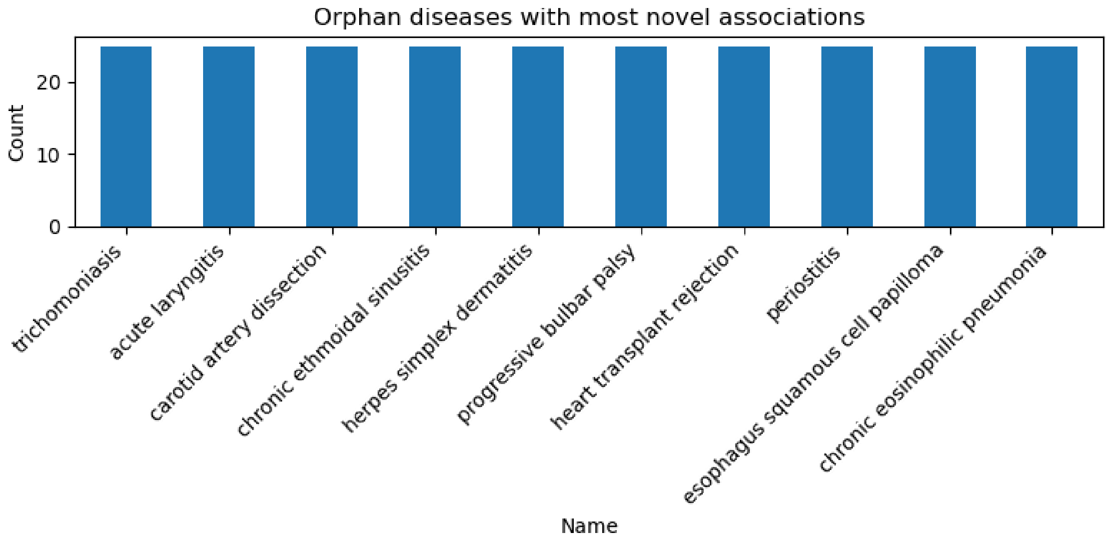


#### 1.5 Novel predictions and evidence gaps

[17]:
```
novel_sorted = compute_novel_predictions(df)
print("Top novel predictions with largest prediction-evidence difference:")
cols_to_show =␣
↪['disease_name','approvedSymbol','pred','evidence_mean','pred_minus_evidence']
display(novel_sorted[cols_to_show].head(6))
```

```
---------------------------------------------------------------------------
KeyError                                  Traceback (most recent call last)
Cell In[17], line 1
----> 1 novel_sorted = compute_novel_predictions(df)
      2 print("Top novel predictions with largest prediction-evidence difference:
  ↪")
      3 cols_to_show =␣
  ↪['disease_name','approvedSymbol','pred','evidence_mean','pred_minus_evidence'

Cell In[3], line 112, in compute_novel_predictions(df)
    108 score_cols = ['score_direct', 'score_known_drug', 'score_literature',
    109               'score_affected_pathway', 'score_genetic_association',
    110               'score_rna_expression', 'score_somatic_mutation']
    111 novel = df[df['label'] == -1].copy()
--> 112 novel['evidence_mean'] = novel[score_cols].mean(axis=1)
    113 novel['pred_minus_evidence'] = novel['pred'] - novel['evidence_mean']
    114 novel_sorted = novel.sort_values('pred_minus_evidence', ascending=False

File ~/anaconda3/lib/python3.13/site-packages/pandas/core/frame.py:4108, in␣
  ↪DataFrame.__getitem__(self, key)
```

```
   4106     if is_iterator(key):
   4107         key = list(key)
-> 4108     indexer = self.columns._get_indexer_strict(key, "columns")[1]
   4110 # take() does not accept boolean indexers
   4111 if getattr(indexer, "dtype", None) == bool:

File ~/anaconda3/lib/python3.13/site-packages/pandas/core/indexes/base.py:6200,
 ↪in Index._get_indexer_strict(self, key, axis_name)
   6197 else:
   6198     keyarr, indexer, new_indexer = self._reindex_non_unique(keyarr)
-> 6200 self._raise_if_missing(keyarr, indexer, axis_name)
   6202 keyarr = self.take(indexer)
   6203 if isinstance(key, Index):
   6204     # GH 42790 - Preserve name from an Index

File ~/anaconda3/lib/python3.13/site-packages/pandas/core/indexes/base.py:6249,
 ↪in Index._raise_if_missing(self, key, indexer, axis_name)
   6247 if nmissing:
   6248     if nmissing == len(indexer):
-> 6249         raise KeyError(f"None of [{key}] are in the [{axis_name}]")
   6251     not_found = list(ensure_index(key)[missing_mask.nonzero()[0]].
 ↪unique())
   6252     raise KeyError(f"{not_found} not in index")

KeyError: "None of [Index(['score_direct', 'score_known_drug',␣
 ↪'score_literature',\n       'score_affected_pathway',␣
 ↪'score_genetic_association',\n       'score_rna_expression',␣
 ↪'score_somatic_mutation'],\n      dtype='object')] are in the [columns]"
```

#### 1.6 Filter out highly promiscuous genes and diseases

```
[ ]: filtered_df = filter_specific_targets(df[df['label'] == -1],
                                          max_gene_count=200,
                                          max_disease_count=10)
     print(f"Filtered novel dataset shape: {filtered_df.shape}")

     filtered_novel_sorted = compute_novel_predictions(filtered_df)
     print("Top novel predictions after filtering promiscuous genes/diseases:")
     display(filtered_novel_sorted[cols_to_show].head(10))
```

```
[ ]: # Assume df has at least: approvedSymbol, disease_name, pred, (optional) label
     shortlist = shortlist_specific_pairs(
         df,
         pred_min=0.70,          # your confidence threshold
         hub_quantile=0.99,      # drop top 1% hubs by degree
         # score="tfidf",          # or "npmi"
         top_k_per_gene=3
```

```
)
display(shortlist.head(15))
```

##### 1.7 Anomaly detection using IsolationForest

[ ]:
```
anomaly_df = run_isolation_forest(df, contamination=0.1)
print(anomaly_df['anomaly_score'].describe())
print(f"Number of outliers flagged: {anomaly_df['is_outlier'].sum()}")

num_cols =␣
 ↪['pred','num_known_targets','in_aba','score_direct','score_known_drug',
          ␣
 ↪'score_literature','score_affected_pathway','score_genetic_association',
            'score_rna_expression','score_somatic_mutation','anomaly_score']
correlation_matrix = anomaly_df[num_cols].corr()
plt.figure(figsize=(8,6))
sns.heatmap(correlation_matrix, annot=True, fmt='.2f', cmap='coolwarm',␣
 ↪center=0)
plt.title('Correlation matrix of numeric features and anomaly score')
plt.tight_layout()
plt.show()

anomalies = anomaly_df.sort_values('anomaly_score', ascending=False)
print("Top anomalous pairs:")
anomaly_cols =␣
 ↪['disease_name','approvedSymbol','pred','label','score_direct','score_known_drug','score_li
display(anomalies[anomaly_cols].head().round(2))
```

[ ]:
```
display(anomalies.loc[anomalies["label"]<1][anomaly_cols].head(10).round(2))
```

##### 1.8 Summary and next steps

This notebook loaded a large set of disease–target predictions and performed several analyses:

- **Summary statistics:** We examined label distributions and mean evidence scores, revealing that a large proportion of predictions correspond to unknown disease–gene pairs with little supporting evidence.
- **Distribution plots:** The predicted probability distribution is bimodal for known positives but skewed towards high scores for unknowns.
- **Gene and disease coverage:** Many novel predictions involve ubiquitous genes (e.g., tubulin family) and common disease terms. Filtering out highly promiscuous genes and diseases helps surface more specific associations.
- **Evidence gaps:** We computed the difference between the predicted probability and the mean evidence score, highlighting candidate associations where the model is highly confident despite minimal evidence.
- **Anomaly detection:** An IsolationForest flagged about 5% of pairs as anomalous; these typically have high evidence scores but only moderate predicted probabilities. Such discordant

cases may indicate areas where the model and evidence disagree.

Can adjust the filtering thresholds, revisit the evidence gap calculations, or integrate additional features (e.g., pathway membership or tissue specificity) to refine the candidate further.

- Note: many diseases are "missing", in the candidates (due to prior processing and what was kept as candidate or had a positive pred), including early onset alzheimers

[ ]:
```
df.loc[df["disease_name"].str.
 ↪contains("early|onset|youth|infant|variab|alz",case=False,na=False)].
 ↪drop_duplicates("disease_name")
```

[ ]:
```
## do we have male breast cancer? (n vm cancer filter beforehand)
df.loc[df["disease_name"].str.contains("male|breast",case=False,na=False)].
 ↪drop_duplicates("disease_name")
```

[ ]:
```
df.loc[(df["num_known_targets"]<5) & (df["pred"]>0.7) & (df["label"]<1)]
```

[ ]:
```

```
