## Supplemental data S4 for "OTRec: A Deep Learning Recommender for Druggable Disease–Target Prioritization"

### 2-output-EDA-rarity

December 18, 2025

`df_preds` `DL_novel_candidates_predictions.csv` * Combined output and input: contains known positive samples (label==1 / source:`known_positive`), and novel predicted samples (label == -1 / source: `model_prediction`) * Each row/instance is whether a given target (gene) is a clinical candidate (will enter clinical trials) for a given diseases. We have only cases here that were predicted to be positive, or are known positives (i.e are undergoing clinical trials)

Our goal is to analyze this data. and find insights.

- Note that for disease or target specific analyses, need to aggregate/deduplciate/max from many to 1, at diseaseId or targetId level (and avoid using features relevant noyl for disease or target if not analyzing them)
- Disease level: e.g. are target distributions of features different when split by disease type: rare/not rare, simple/complex (omim), known/novel .
  - may subset by known_positive / model_prediction ; AND additional subsets (rare/not rare, simple/complex)
- Target level: # paralogs, orthologs (need to add); distribution of target values between novel and known positives (may need to get max target value first); distribution of target values (inc )
  - e.g. target cases with many vs few preds as subsets: df_novel[“targetSymbol”].value_counts().head(20) vs those with 1-2 preds

In all cases output code for generation statistical analyses and useful plots. Highlight any insights or novel findings.

[1]:
```python
import pandas as pd
import re
import os
# from sklearn.feature_extraction.text import TfidfVectorizer
# from sklearn.linear_model            import LogisticRegression
from sklearn.metrics                  import roc_auc_score
import numpy as np
from sklearn.metrics import classification_report
```

[2]:
```python
# DATA_DIR = "../data/opentargets/"
```

[3]:
```python
# target_df = pd.read_parquet("../data/proc/target_df.parquet")
# target_df[['targetId', 'approvedSymbol', 'biotype', 'approvedName', 'go',
#        'functionDescriptions', 'subcellularLocations', 'targetClass',
#        'constraint', 'tep', 'proteinIds', 'tractability',
```

```
#         'pathways', 'count_synonyms', 'count_subcellularLocations', 'count_go',
#         'count_pathways', 'count_proteinIds', 'count_targetClass',
#         'count_safetyLiabilities', 'count_tractability',
#         'count_alternativeGenes', 'count_tep',
#         'score_syn', 'score_mis', 'score_lof']].to_parquet(("./copy_proc/
↪subset/target_df.parquet"))
```

[4]:
```
# disease_df = pd.read_parquet("../data/proc/disease_df.parquet")

# ## add columns for rare diseases, simple (omim - single gene) diseases
# # Ensure columns are treated as strings to avoid errors with NaN values
# disease_df['diseaseId'] = disease_df['diseaseId'].astype(str)
# disease_df['dbXRefs'] = disease_df['dbXRefs'].astype(str)

# # 1. Define Rare Disease (Exact match for Orphanet ID or reference)
# disease_df['disease_rare'] = disease_df['diseaseId'].str.
↪contains('Orphanet|ORPHA',case=False) | disease_df['dbXRefs'].str.
↪contains('Orphan|ORPHA',case=False)
# print(disease_df['disease_rare'].value_counts())
# # 2. Define Mendelian/Simple Disease (Exact match for OMIM ID or reference)
# disease_df['disease_has_omim'] = disease_df['diseaseId'].str.
↪contains('OMIM',case=False) | disease_df['dbXRefs'].str.
↪contains('OMIM',case=False)
# print(disease_df['disease_has_omim'].value_counts())
# print(disease_df.shape)
# display(disease_df.head(2))

# disease_df[['diseaseId', 'name', 'description', 'dbXRefs',␣
↪'therapeuticAreas',
#         'disease_rare', 'disease_has_omim']].to_parquet(("./copy_proc/subset/
↪disease_df.parquet"))
```

[5]:
```
disease_df = pd.read_parquet("./copy_proc/subset/disease_df.parquet")
disease_df
```

[5]:
```
             diseaseId                                              name  \
0         DOID_0050890                                   synucleinopathy
1           DOID_10113                                   trypanosomiasis
2           DOID_10718                                        giardiasis
3           DOID_13406                              pulmonary sarcoidosis
4            DOID_1947                                    trichomoniasis
…                  …                                                 …
38954   Orphanet_99942  Autosomal dominant Charcot-Marie-Tooth disease…
38955   Orphanet_99943  Autosomal dominant Charcot-Marie-Tooth disease…
38956   Orphanet_99945  Autosomal dominant Charcot-Marie-Tooth disease…
38957   Orphanet_99946  Autosomal dominant Charcot-Marie-Tooth disease…
38958   Orphanet_99947  Autosomal dominant Charcot-Marie-Tooth disease…
```

```
                                             description  \
0      A neurodegenerative disease that is characteri…
1      Infection with protozoa of the genus trypanosoma.
2      An infection of the small intestine caused by …
3      Sarcoidosis affecting the lung parenchyma. It …
4            An infection that is caused by Trichomonas.
…                                                    …
38954  Autosomal dominant Charcot-Marie-Tooth disease…
38955  Autosomal dominant Charcot-Marie-Tooth disease…
38956  Autosomal dominant Charcot-Marie-Tooth disease…
38957  Autosomal dominant Charcot-Marie-Tooth disease…
38958                                             None

                                                 dbXRefs  \
0      ['MESH:D000080874' 'MONDO:0000510' 'UMLS:C5191…
1      ['UMLS:C0041227' 'MONDO:0000940' 'ICD10CM:B56'…
2      ['MeSH:D005873' 'ICD9CM:007.1' 'MESH:D005873' …
3      ['SNOMEDCT:187230004' 'UMLS:C0036205' 'MONDO:0…
4      ['ICD9:131.8' 'MESH:D014245' 'ICD10CM:A59' 'IC…
…                                                    …
38954                    ['ICD10:G60.0' 'OMIM:607677']
38955                    ['ICD10:G60.0' 'OMIM:607736']
38956                    ['ICD10:G60.0' 'OMIM:608673']
38957                    ['ICD10:G60.0' 'OMIM:118210']
38958                    ['ICD10:G60.0' 'OMIM:609260']

                                 therapeuticAreas  disease_rare  \
0      [EFO_0000618, OTAR_0000018, OTAR_0000020]         False
1                                   [EFO_0005741]         False
2                    [EFO_0010282, EFO_0005741]         False
3                  [OTAR_0000010, OTAR_0000006]         False
4                                   [EFO_0005741]         False
…                                             …           …
38954               [OTAR_0000018, EFO_0000618]          True
38955               [EFO_0000618, OTAR_0000018]          True
38956               [OTAR_0000018, EFO_0000618]          True
38957               [EFO_0000618, OTAR_0000018]          True
38958               [OTAR_0000018, EFO_0000618]          True

       disease_has_omim
0                 False
1                 False
2                 False
3                 False
4                 False
…                     …
```

```
38954               True
38955               True
38956               True
38957               True
38958               True

[38959 rows x 7 columns]
```

[6]:
```python
disease_df.columns
```

[6]:
```
Index(['diseaseId', 'name', 'description', 'dbXRefs', 'therapeuticAreas',
       'disease_rare', 'disease_has_omim'],
      dtype='object')
```

[7]:
```python
# df_learn = pd.read_parquet("../data/proc/df_learn.parquet")
# print(df_learn.shape)
# display(df_learn)

target_df = pd.read_parquet("./copy_proc/target_df.parquet")
print(target_df.shape)
display(target_df.head(2))
```

```
(17065, 35)
          targetId approvedSymbol         biotype  \
0  ENSG00000000457          SCYL3  protein_coding
1  ENSG00000001167           NFYA  protein_coding

                                    genomicLocation alternativeGenes  \
0  {'chromosome': '1', 'end': 169894267, 'start':…             None
1  {'chromosome': '6', 'end': 41102403, 'start': …             None

                                approvedName  \
0                        SCY1 like pseudokinase 3
1  nuclear transcription factor Y subunit alpha

                                                  go  hallmarks synonyms  \
0  [GO:0005737, GO:0042802, GO:0016477, GO:000551…       NaN
1  [GO:0000785, GO:0005515, GO:0016602, GO:000635…       NaN

                                functionDescriptions  … count_tractability  \
0  May play a role in regulating cell adhesion/mi…  …                  1
1  Component of the sequence-specific heterotrime…  …                  2

  count_alternativeGenes count_hallmarks  count_functionDescriptions  \
0                      0               0                           1
1                      0               0                           1
```

```
  count_tep  sym                               target_text_embed  \
0         0  SCYL  SCYL SCY1 like pseudokinase 3  May play a role…
1         0  NFYA  NFYA nuclear transcription factor Y subunit al…

  score_syn  score_mis  score_lof
0   0.70818    0.98492    0.28151
1  -0.17463    2.78030    0.14619

[2 rows x 35 columns]
```

[8]:
```
target_df.select_dtypes("number").columns
```

[8]:
```
Index(['hallmarks', 'tep', 'count_synonyms', 'count_subcellularLocations',
       'count_go', 'count_pathways', 'count_proteinIds', 'count_targetClass',
       'count_safetyLiabilities', 'count_tractability',
       'count_alternativeGenes', 'count_hallmarks',
       'count_functionDescriptions', 'count_tep', 'score_syn', 'score_mis',
       'score_lof'],
      dtype='object')
```

[9]:
```
df_preds = pd.read_csv("DL_novel_candidates_predictions.csv.gz") # DL preds,␣
 ↪novel and positive

## join extra disease level info?
df_preds = df_preds.
 ↪merge(disease_df[['diseaseId','disease_has_omim','disease_rare']],on='diseaseId',how="left"
df_preds
```

[9]:
```
              diseaseId                          diseaseName         targetId  \
0        UBERON_0000104                           life cycle  ENSG00000123201
1        Orphanet_99878      Primary parathyroids hyperplasia  ENSG00000123201
2        Orphanet_99878      Primary parathyroids hyperplasia  ENSG00000261456
3        Orphanet_99842   Leukocyte adhesion deficiency type I  ENSG00000185231
4        Orphanet_99842   Leukocyte adhesion deficiency type I  ENSG00000106348
…                   …                                    …                …
190639       DOID_10113                      trypanosomiasis  ENSG00000151834
190640       DOID_10113                      trypanosomiasis  ENSG00000163285
190641       DOID_10113                      trypanosomiasis  ENSG00000182256
190642       DOID_10113                      trypanosomiasis  ENSG00000183454
190643       DOID_10113                      trypanosomiasis  ENSG00000186297

       targetSymbol  score  label            source  \
0           GUCY1B2  0.924     -1  model_prediction
1           GUCY1B2  0.878     -1  model_prediction
2             TUBB8  0.853     -1  model_prediction
3              MC2R  0.845     -1  model_prediction
4            IMPDH1  0.812     -1  model_prediction
```

```
…               …     …     …             …
190639       GABRA2  1.000      1    known_positive
190640       GABRG1  1.000      1    known_positive
190641       GABRG3  1.000      1    known_positive
190642       GRIN2A  1.000      1    known_positive
190643       GABRA5  1.000      1    known_positive

        disease_num_known_clinical_targets  orphan  disease_has_omim  \
0                                        0    True             False
1                                        0    True              True
2                                        0    True              True
3                                        1   False              True
4                                        1   False              True
…                                      …      …                …
190639                                  11   False             False
190640                                  11   False             False
190641                                  11   False             False
190642                                  11   False             False
190643                                  11   False             False

        disease_rare
0              False
1               True
2               True
3               True
4               True
…                …
190639         False
190640         False
190641         False
190642         False
190643         False

[190644 rows x 11 columns]
```

[10]:
```
df_known_pos = df_preds.query("label>0")
df_known_pos
```

[10]:
```
             diseaseId                                 diseaseName  \
6       Orphanet_99842              Leukocyte adhesion deficiency type I
12      Orphanet_98974               Fuchs endothelial corneal dystrophy
13      Orphanet_98974               Fuchs endothelial corneal dystrophy
71      Orphanet_98920  Spinal muscular atrophy with respiratory distr…
72      Orphanet_98920  Spinal muscular atrophy with respiratory distr…
…                  …                                               …
190639      DOID_10113                                 trypanosomiasis
190640      DOID_10113                                 trypanosomiasis
```

```
190641      DOID_10113                                    trypanosomiasis
190642      DOID_10113                                    trypanosomiasis
190643      DOID_10113                                    trypanosomiasis

               targetId targetSymbol  score  label          source  \
6       ENSG00000113302        IL12B    1.0      1  known_positive
12      ENSG00000067900        ROCK1    1.0      1  known_positive
13      ENSG00000134318        ROCK2    1.0      1  known_positive
71      ENSG00000172062         SMN1    1.0      1  known_positive
72      ENSG00000205571         SMN2    1.0      1  known_positive
…                   …          …    …    …             …
190639  ENSG00000151834       GABRA2    1.0      1  known_positive
190640  ENSG00000163285       GABRG1    1.0      1  known_positive
190641  ENSG00000182256       GABRG3    1.0      1  known_positive
190642  ENSG00000183454       GRIN2A    1.0      1  known_positive
190643  ENSG00000186297       GABRA5    1.0      1  known_positive

        disease_num_known_clinical_targets  orphan  disease_has_omim  \
6                                        1   False              True
12                                       2   False              True
13                                       2   False              True
71                                       2   False              True
72                                       2   False              True
…                                      …      …                …
190639                                  11   False             False
190640                                  11   False             False
190641                                  11   False             False
190642                                  11   False             False
190643                                  11   False             False

        disease_rare
6               True
12              True
13              True
71              True
72              True
…                …
190639         False
190640         False
190641         False
190642         False
190643         False

[67532 rows x 11 columns]
```

#### 0.1 Analyze novel predictions

[11]:
```
df_novel = df_preds.query("label<1")
df_novel
```

[11]:
```
              diseaseId                          diseaseName        targetId  \
0       UBERON_0000104                           life cycle  ENSG00000123201
1        Orphanet_99878      Primary parathyroids hyperplasia  ENSG00000123201
2        Orphanet_99878      Primary parathyroids hyperplasia  ENSG00000261456
3        Orphanet_99842  Leukocyte adhesion deficiency type I  ENSG00000185231
4        Orphanet_99842  Leukocyte adhesion deficiency type I  ENSG00000106348
…                     …                                    …               …
190628       DOID_10113                      trypanosomiasis  ENSG00000125675
190629       DOID_10113                      trypanosomiasis  ENSG00000152578
190630       DOID_10113                      trypanosomiasis  ENSG00000120251
190631       DOID_10113                      trypanosomiasis  ENSG00000153956
190632       DOID_10113                      trypanosomiasis  ENSG00000166148

       targetSymbol  score  label            source  \
0           GUCY1B2  0.924     -1  model_prediction
1           GUCY1B2  0.878     -1  model_prediction
2             TUBB8  0.853     -1  model_prediction
3              MC2R  0.845     -1  model_prediction
4            IMPDH1  0.812     -1  model_prediction
…                 …      …      …                 …
190628        GRIA3  0.739     -1  model_prediction
190629        GRIA4  0.736     -1  model_prediction
190630        GRIA2  0.716     -1  model_prediction
190631     CACNA2D1  0.715     -1  model_prediction
190632       AVPR1A  0.710     -1  model_prediction

        disease_num_known_clinical_targets  orphan  disease_has_omim  \
0                                        0    True             False
1                                        0    True              True
2                                        0    True              True
3                                        1   False              True
4                                        1   False              True
…                                        …       …                 …
190628                                  11   False             False
190629                                  11   False             False
190630                                  11   False             False
190631                                  11   False             False
190632                                  11   False             False

        disease_rare
0              False
1               True
```

```
2                True
3                True
4                True
…               …
190628          False
190629          False
190630          False
190631          False
190632          False

[123112 rows x 11 columns]
```

[12]:
```
df_novel.nunique()
```

[12]:
```
diseaseId                           4907
diseaseName                         4906
targetId                            2115
targetSymbol                        2113
score                                300
label                                  1
source                                 1
disease_num_known_clinical_targets   211
orphan                                 2
disease_has_omim                       2
disease_rare                           2
dtype: int64
```

**Overall distribution of genes predicted to be with different diseases:**

- We see aclear long tail

[13]:
```
df_novel["targetSymbol"].value_counts().hist()
```

[13]:
```
<Axes: >
```

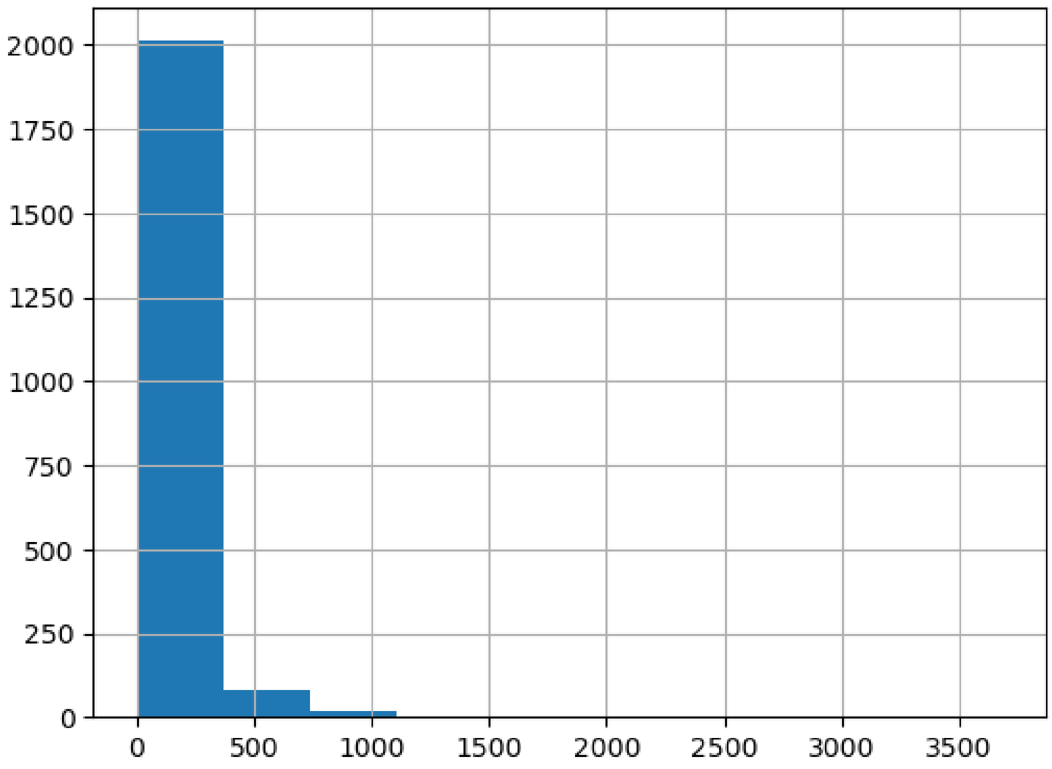


[14]:
```
df_novel["targetSymbol"].value_counts().describe().round(0)
```

[14]:
```
count    2113.0
mean       58.0
std       170.0
min         1.0
25%         1.0
50%         6.0
75%        27.0
max      3685.0
Name: count, dtype: float64
```

**genes predicted to be with many diseases:**

[15]:
```
df_novel["targetSymbol"].value_counts().head(20)
```

[15]:
```
targetSymbol
GUCY1B2    3685
TUBB8      2532
FKBP1A     1507
TUBB8B     1254
TUBB6      1182
```

```
TOP1MT     1112
POLD2      1073
TUBB1      1055
TUBB4A      986
TUBB4B      880
TUBB        857
TUBA3D      844
GABRA5      826
TUBA3E      808
ADRA2C      804
GABRA2      801
GABRG3      796
PDE4C       781
TUBAL3      775
TUBA8       765
Name: count, dtype: int64
```

###### genes predicted to be with few diseases (but still at least 1)

- Can we charactertize those associated with few cases?
- matching with known diseases?

[16]:
```
df_novel["targetSymbol"].value_counts().tail(15)
```

[16]:
```
targetSymbol
TRPM7      1
SLC12A9    1
SIGLEC7    1
NCR1       1
NCR2       1
HPGDS      1
EPHX1      1
TBK1       1
C5AR1      1
CD180      1
C5AR2      1
TRPM2      1
DAGLB      1
CES2       1
MCOLN2     1
Name: count, dtype: int64
```

- 540 targets are predicted to assoicate with 1 novel disease

[17]:
```
(df_novel["targetSymbol"].value_counts()<2).sum()
```

[17]:
```
540
```

[18]: disease_df

[18]:
```
              diseaseId                                             name  \
0          DOID_0050890                                   synucleinopathy
1            DOID_10113                                  trypanosomiasis
2            DOID_10718                                       giardiasis
3            DOID_13406                             pulmonary sarcoidosis
4             DOID_1947                                   trichomoniasis
…                   …                                                …
38954    Orphanet_99942  Autosomal dominant Charcot-Marie-Tooth disease…
38955    Orphanet_99943  Autosomal dominant Charcot-Marie-Tooth disease…
38956    Orphanet_99945  Autosomal dominant Charcot-Marie-Tooth disease…
38957    Orphanet_99946  Autosomal dominant Charcot-Marie-Tooth disease…
38958    Orphanet_99947  Autosomal dominant Charcot-Marie-Tooth disease…

                                            description  \
0       A neurodegenerative disease that is characteri…
1       Infection with protozoa of the genus trypanosoma.
2       An infection of the small intestine caused by …
3       Sarcoidosis affecting the lung parenchyma. It …
4             An infection that is caused by Trichomonas.
…                                                     …
38954   Autosomal dominant Charcot-Marie-Tooth disease…
38955   Autosomal dominant Charcot-Marie-Tooth disease…
38956   Autosomal dominant Charcot-Marie-Tooth disease…
38957   Autosomal dominant Charcot-Marie-Tooth disease…
38958                                              None

                                                dbXRefs  \
0       ['MESH:D000080874' 'MONDO:0000510' 'UMLS:C5191…
1       ['UMLS:C0041227' 'MONDO:0000940' 'ICD10CM:B56'…
2       ['MeSH:D005873' 'ICD9CM:007.1' 'MESH:D005873' …
3       ['SNOMEDCT:187230004' 'UMLS:C0036205' 'MONDO:0…
4       ['ICD9:131.8' 'MESH:D014245' 'ICD10CM:A59' 'IC…
…                                                     …
38954                    ['ICD10:G60.0' 'OMIM:607677']
38955                    ['ICD10:G60.0' 'OMIM:607736']
38956                    ['ICD10:G60.0' 'OMIM:608673']
38957                    ['ICD10:G60.0' 'OMIM:118210']
38958                    ['ICD10:G60.0' 'OMIM:609260']

                               therapeuticAreas  disease_rare  \
0       [EFO_0000618, OTAR_0000018, OTAR_0000020]         False
1                                   [EFO_0005741]         False
2                     [EFO_0010282, EFO_0005741]         False
3                    [OTAR_0000010, OTAR_0000006]         False
4                                   [EFO_0005741]         False
```

```
…                                       …            …
38954               [OTAR_0000018, EFO_0000618]          True
38955               [EFO_0000618, OTAR_0000018]          True
38956               [OTAR_0000018, EFO_0000618]          True
38957               [EFO_0000618, OTAR_0000018]          True
38958               [OTAR_0000018, EFO_0000618]          True

       disease_has_omim
0                 False
1                 False
2                 False
3                 False
4                 False
…                 …
38954              True
38955              True
38956              True
38957              True
38958              True

[38959 rows x 7 columns]
```

[19]:
```python
(df_novel["targetSymbol"].value_counts()<2).sum()
```

[19]:
```
540
```

**extra target level features** Load targetability - and max clinical phase reached - max clinical trial pahse is per target, NOT target X disease

[20]:
```python
# target_priority = pd.read_parquet(os.path.join(DATA_DIR,␣
↪"target_prioritisation")).dropna(axis=0,thresh=2)#.
↪dropna(subset=["maxClinicalTrialPhase"])
# target_priority.to_parquet("./copy_proc/target_prioritisation.
↪parquet",index=False)
```

[21]:
```python
target_priority = pd.read_parquet("./copy_proc/target_prioritisation.parquet")
target_priority
```

[21]:
```
              targetId  isInMembrane  isSecreted  hasSafetyEvent  hasPocket  \
0      ENSG00000000457           0.0         0.0             NaN        0.0
1      ENSG00000001167           0.0         0.0             NaN        0.0
2      ENSG00000001460           0.0         0.0             NaN        0.0
3      ENSG00000001629           1.0         0.0             NaN        0.0
4      ENSG00000003096           0.0         0.0             NaN        0.0
…                    …           …          …               …          …
26158  ENSG00000290316           NaN         NaN             NaN        NaN
26159  ENSG00000291237           0.0         0.0            -1.0        1.0
```

```
26160  ENSG00000295847          NaN         NaN              NaN        NaN
26161  ENSG00000295937          NaN         NaN              NaN        NaN
26162  ENSG00000305532          NaN         NaN              NaN        NaN

       hasLigand  hasSmallMoleculeBinder  geneticConstraint  \
0            0.0                     0.0          -0.598145
1            0.0                     0.0          -0.487914
2            0.0                     0.0           0.487914
3            0.0                     0.0          -0.880496
4            0.0                     0.0          -0.355595
…            …                     …                …
26158        NaN                     NaN                NaN
26159        0.0                     0.0                NaN
26160        NaN                     NaN                NaN
26161        NaN                     NaN                NaN
26162        NaN                     NaN                NaN

       paralogMaxIdentityPercentage  mouseOrthologMaxIdentityPercentage  \
0                          0.000000                            0.142440
1                               NaN                            0.985590
2                          0.000000                            0.000000
3                          0.000000                            0.664830
4                         -0.726135                            0.640065
…                             …                                …
26158                           NaN                                 NaN
26159                           NaN                            0.504505
26160                     -1.000000                                 NaN
26161                     -1.000000                                 NaN
26162                     -1.000000                                 NaN

       isCancerDriverGene  hasTEP  mouseKOScore  hasHighQualityChemicalProbes  \
0                     NaN     NaN      0.000000                           NaN
1                     NaN     NaN     -0.276242                           NaN
2                     NaN     NaN           NaN                           NaN
3                     NaN     NaN     -0.166128                           NaN
4                     NaN     NaN           NaN                           NaN
…                     …     …           …                           …
26158                 NaN     NaN           NaN                           NaN
26159                 NaN     NaN     -0.973375                           NaN
26160                 NaN     NaN           NaN                           NaN
26161                 NaN     NaN           NaN                           NaN
26162                 NaN     NaN           NaN                           NaN

       maxClinicalTrialPhase  tissueSpecificity  tissueDistribution
0                        NaN               -1.0                -1.0
1                        NaN               -1.0                -1.0
2                        NaN                0.5                -1.0
```

```
3                         NaN              -1.0                 -1.0
4                         NaN               0.5                  0.0
…                       …                 …                  …
26158                     NaN               0.5                  1.0
26159                     NaN               0.5                 -1.0
26160                     NaN               NaN                  NaN
26161                     NaN               NaN                  NaN
26162                     NaN               NaN                  NaN

[26163 rows x 17 columns]
```

[22]:

```
# target_priority.columns
[col for col in target_priority.select_dtypes("number").columns if␣
↪target_priority[col].nunique() > 3]
```

[22]:

```
['geneticConstraint',
 'paralogMaxIdentityPercentage',
 'mouseOrthologMaxIdentityPercentage',
 'mouseKOScore',
 'maxClinicalTrialPhase',
 'tissueSpecificity',
 'tissueDistribution']
```

[23]:

```
temp = df_known_pos.merge(target_priority,on=["targetId"],how="inner")
print(temp.shape[0])
print(temp.maxClinicalTrialPhase.corr(temp.score))
```

```
67532
nan

/home/ddofer/anaconda3/lib/python3.10/site-
packages/numpy/lib/function_base.py:2897: RuntimeWarning: invalid value
encountered in divide
  c /= stddev[:, None]
/home/ddofer/anaconda3/lib/python3.10/site-
packages/numpy/lib/function_base.py:2898: RuntimeWarning: invalid value
encountered in divide
  c /= stddev[None, :]
```

[ ]:

```

```

### 1 EDA

- Compare both novel/non novel ; and analyses on target or disease level

[24]:

```
import pandas as pd
import numpy as np
```

```python
import seaborn as sns
import matplotlib.pyplot as plt
from scipy import stats
import ast

# Set visual style
sns.set(style="whitegrid", context="talk")
plt.rcParams['figure.figsize'] = (12, 8)

# --- CONFIGURATION: Feature Descriptions ---
FEATURE_DESCRIPTIONS = {
    "isInMembrane": "Indicates if the target protein is located in the cell or␣
 ↪plasma membrane",
    "isSecreted": "Indicates if the target protein is secreted or predicted to␣
 ↪be secreted",
    "hasSafetyEvent": "Indicates if there are known safety events associated␣
 ↪with the target",
    "hasPocket": "Indicates if the target has predicted binding pockets␣
 ↪suitable for small molecule binding",
    "hasLigand": "Indicates if the target binds at least one high-quality␣
 ↪ligand",
    "hasSmallMoleculeBinder": "Indicates if the target has at least one small␣
 ↪molecule binder",
    "geneticConstraint": "Represents the genetic constraint of the target,␣
 ↪indicating how intolerant it is to genetic variation",
    "paralogMaxIdentityPercentage": "Maximum percentage identity among the␣
 ↪target's paralogues",
    "mouseOrthologMaxIdentityPercentage": "Maximum percentage identity between␣
 ↪the target and its mouse ortholog",
    "isCancerDriverGene": "Indicates if the target is identified as a cancer␣
 ↪driver gene",
    "hasTEP": "Indicates if a Target Enabling Package (TEP) is available for␣
 ↪the target",
    "mouseKOScore": "Represents the phenotypic score from mouse knockout models␣
 ↪for the target",
    "hasHighQualityChemicalProbes": "Indicates if there are high-quality␣
 ↪chemical probes available for the target",
    "maxClinicalTrialPhase": "Highest clinical trial phase that the target has␣
 ↪reached for any indication",
    "tissueSpecificity": "Describes the specificity of the target's expression␣
 ↪across different tissues",
    "tissueDistribution": "Describes the distribution pattern of the target's␣
 ↪expression in various tissues",
    # Scores (gnomAD)
    "score_lof": "Loss-of-function intolerance score (pLI/LOEUF likely)",
    "score_mis": "Missense intolerance score (Z-score likely)",
```

```python
    "score_syn": "Synonymous intolerance score (Z-score likely)"
}

def extract_clean_label(val):
    """
    Safely extracts a readable label from complex targetClass objects.
    """
    if pd.isna(val) or val == "":
        return "Unknown"

    # If it's already a dictionary
    if isinstance(val, dict):
        return val.get('label', str(val))

    # If it's a string, it might be a stringified dict or just a label
    if isinstance(val, str):
        val = val.strip()
        if val.startswith('{') and 'label' in val:
            try:
                # distinct safe eval
                parsed = ast.literal_eval(val)
                if isinstance(parsed, dict):
                    return parsed.get('label', val)
            except (ValueError, SyntaxError):
                pass

    return str(val)

def calculate_cohens_d(group1, group2):
    """
    Calculates Cohen's d effect size.
    """
    diff = group1.mean() - group2.mean()
    n1, n2 = len(group1), len(group2)
    var1 = group1.var()
    var2 = group2.var()

    # Pooled standard deviation
    pooled_var = ((n1 - 1) * var1 + (n2 - 1) * var2) / (n1 + n2 - 2)
    pooled_sd = np.sqrt(pooled_var)

    return diff / pooled_sd

def load_and_merge_data():
    """
    Loads data from the specific subset paths and pre-processes.
    """
```

```
    print("Loading data...")

    # --- 1. Load Data (Using updated paths) ---
    try:
        preds = pd.read_csv('./copy_proc/subset/DL_novel_candidates_predictions.
↪csv.gz')
    except FileNotFoundError:
        print("Gzipped prediction file not found, trying unzipped...")
        preds = pd.read_csv('./copy_proc/subset/DL_novel_candidates_predictions.
↪csv')

    disease_df = pd.read_parquet('./copy_proc/subset/disease_df.parquet')
    target_df = pd.read_parquet('./copy_proc/subset/target_df.parquet')
    target_prioritisation = pd.read_parquet('./copy_proc/subset/
↪target_prioritisation.parquet')

    # --- 2. Merging ---
    print("Merging metadata...")

    # Merge Disease Metadata
    disease_df = disease_df.rename(columns={'id': 'diseaseId'})
    if 'disease_rare' in disease_df.columns:
        disease_df['disease_rare'] = disease_df['disease_rare'].astype(bool)

    df = preds.merge(disease_df, on='diseaseId', how='left', suffixes=('',␣
↪'_meta'))

    # Merge Target Metadata
    target_df = target_df.rename(columns={'id': 'targetId'})
    df = df.merge(target_df, on='targetId', how='left', suffixes=('',␣
↪'_target_meta'))

    # Merge Target Prioritisation
    target_prioritisation = target_prioritisation.rename(columns={'id':␣
↪'targetId'})
    df = df.merge(target_prioritisation, on='targetId', how='left')

    # --- 3. Label Normalization ---
    # Create a clean 'type' column for Known vs Novel
    if 'source' in df.columns:
        df['type'] = df['source'].apply(lambda x: 'Known Positive' if 'known'␣
↪in str(x).lower() else 'Novel Prediction')
    elif 'label' in df.columns:
        df['type'] = df['label'].apply(lambda x: 'Known Positive' if x == 1␣
↪else 'Novel Prediction')
    else:
```

```
        df['type'] = 'Unknown'

    # --- 4. Derive Missing Columns ---
    # Ensure 'disease_has_omim' exists
    if 'disease_has_omim' not in df.columns:
        if 'dbXRefs' in df.columns:
             # Handle cases where dbXRefs might be lists or strings
             df['disease_has_omim'] = df['dbXRefs'].astype(str).str.
↪contains('OMIM', case=False, na=False)
        else:
             df['disease_has_omim'] = False
             print("Warning: 'disease_has_omim' could not be derived.␣
↪Defaulting to False.")

    print(f"Data merged successfully. Shape: {df.shape}")
    print(f"Split counts:\n{df['type'].value_counts()}")
    return df

# --- ANALYSIS FUNCTIONS ---

def analyze_orphan(df):
    """
    Compares scores between Orphan and Non-Orphan diseases.
    """
    print("\n--- Orphan vs Non-Orphan Analysis ---")

    if 'orphan' in df.columns:
        if df['orphan'].nunique() < 2:
            print("Insufficient data: Less than 2 groups found for 'orphan'.")
            return

        plt.figure(figsize=(8, 6))
        sns.boxplot(x='orphan', y='score', data=df, palette='Set2')
        plt.title('Prediction Scores: Orphan vs Non-Orphan Diseases')
        plt.xlabel('Is Orphan Disease?')
        plt.ylabel('Model Confidence Score')
        plt.show()

        # Mann-Whitney U Test
        group_true = df[df['orphan'] == True]['score']
        group_false = df[df['orphan'] == False]['score']

        if len(group_true) > 0 and len(group_false) > 0:
            stat, p_val = stats.mannwhitneyu(group_true, group_false)
            print(f"Mann-Whitney U Test: P-value = {p_val:.4e}")
            print("Result: " + ("Significant difference." if p_val < 0.05 else␣
↪"No significant difference."))
```

```
    else:
        print("Column 'orphan' not found.")

def analyze_rarity(df):
    """
    Compares scores between Rare vs Non-rare diseases.
    """
    print("\n--- Rarity Analysis (Rare vs Non-rare) ---")

    col_name = 'disease_rare'
    if col_name in df.columns:
        df_clean = df.dropna(subset=[col_name])

        if df_clean[col_name].nunique() < 2:
            print(f"Insufficient data for '{col_name}'.")
            return

        plt.figure(figsize=(8, 6))
        sns.boxplot(x=col_name, y='score', data=df_clean, palette='Set2')
        plt.title('Prediction Scores: Rare vs Non-rare')
        plt.xlabel('Is Rare Disease?')
        plt.ylabel('Model Confidence Score')
        plt.show()

        # Stats
        group_true = df_clean[df_clean[col_name] == True]['score']
        group_false = df_clean[df_clean[col_name] == False]['score']

        if len(group_true) > 0 and len(group_false) > 0:
            stat, p_val = stats.mannwhitneyu(group_true, group_false)
            print(f"Mann-Whitney U Test: P-value = {p_val:.4e}")
            print("Result: " + ("Significant difference." if p_val < 0.05 else␣
 ↪"No significant difference."))
    else:
        print(f"Column '{col_name}' not found.")

def analyze_omim(df):
    """
    Compares scores between simple (OMIM) and complex diseases.
    """
    print("\n--- OMIM/Complexity Analysis ---")

    col_name = 'disease_has_omim'
    if col_name in df.columns:
        df_clean = df.copy()
        df_clean[col_name] = df_clean[col_name].astype(bool)
```

```
        if df_clean[col_name].nunique() < 2:
            print("Insufficient data for OMIM analysis.")
            return

        plt.figure(figsize=(8, 6))
        sns.boxplot(x=col_name, y='score', data=df_clean, palette='Set2')
        plt.title('Prediction Scores: OMIM (Simple) vs Non-OMIM (Complex)')
        plt.xlabel('Has OMIM Annotation?')
        plt.ylabel('Model Confidence Score')
        plt.show()

        # Stats
        group_true = df_clean[df_clean[col_name] == True]['score']
        group_false = df_clean[df_clean[col_name] == False]['score']

        if len(group_true) > 0 and len(group_false) > 0:
            stat, p_val = stats.mannwhitneyu(group_true, group_false)
            print(f"Mann-Whitney U Test: P-value = {p_val:.4e}")
            print("Result: " + ("Significant difference." if p_val < 0.05 else␣
↪"No significant difference."))
    else:
        print(f"Column '{col_name}' not found.")

def analyze_score_distributions(df):
    print("\n--- Score Distribution ---")
    plt.figure(figsize=(10, 6))
    sns.histplot(df['score'], bins=50, kde=True, color='teal')
    plt.title('Distribution of Prediction Scores')
    plt.show()
    print(df['score'].describe())

# --- TARGET LEVEL ANALYSES ---

def analyze_target_prioritization_features(df):
    """
    Compares 'Prioritization Features' between Known (Established) and Novel␣
↪(Purely Predicted) targets.
    Aggregates to 1 row per target using strict logic.

    UPDATED: Includes Extended Feature List (Counts and Scores)
    """
    print("\n--- Target Prioritization Feature Analysis (Known vs Novel) ---")

    # 1. Deduplicate
    df_targets = df.sort_values("label", ascending=False).
↪drop_duplicates(subset="targetId").copy()
```

```python
print(f"Total Unique Targets: {len(df_targets)}")
print(f"Breakdown by Type:\n{df_targets['type'].value_counts()}")

# Extended list of numeric features
numeric_features = [
    # Prioritization
    'geneticConstraint',
    'paralogMaxIdentityPercentage',
    'mouseOrthologMaxIdentityPercentage',
    'mouseKOScore',
    'maxClinicalTrialPhase',
    'tissueSpecificity',
    'tissueDistribution',
    # New Count/Score Features
    'count_synonyms', 'count_subcellularLocations',
    'count_go', 'count_pathways', 'count_proteinIds', 'count_targetClass',
    'count_safetyLiabilities', 'count_tractability',
    'count_alternativeGenes', 'count_hallmarks',
    'count_functionDescriptions', 'count_tep',
    'score_syn', 'score_mis', 'score_lof'
]

boolean_features = [
    'isCancerDriverGene', 'hasTEP', 'hasHighQualityChemicalProbes',
    'isInMembrane', 'isSecreted', 'hasSafetyEvent', 'hasPocket',
    'hasLigand', 'hasSmallMoleculeBinder'
]

# 2. Numeric Feature Analysis
for feat in numeric_features:
    if feat in df_targets.columns:
        # Drop NaNs for this specific feature
        data = df_targets.dropna(subset=[feat])

        if data['type'].nunique() < 2:
            continue

        # PRINT DESCRIPTION
        desc = FEATURE_DESCRIPTIONS.get(feat, "")
        if desc:
            print(f"\n[?] {feat}: {desc}")

        known_vals = data[data['type'] == 'Known Positive'][feat]
        novel_vals = data[data['type'] == 'Novel Prediction'][feat]

        if len(known_vals) > 0 and len(novel_vals) > 0:
            stat, p_val = stats.mannwhitneyu(known_vals, novel_vals)
```

```
                # --- EFFECT SIZE CALCULATION ---
                cohens_d = calculate_cohens_d(known_vals, novel_vals)
                median_diff = known_vals.median() - novel_vals.median()

                print(f"Numeric Feature '{feat}':")
                print(f"  P-value    = {p_val:.4e}")
                print(f"  Cohen's d  = {cohens_d:.4f} (Effect Size)")
                print(f"  Median Diff= {median_diff:.4f}")

                # Check Significance AND Effect Size
                if p_val < 0.05 and abs(cohens_d) >= 0.2:
                    print(f"-> SIGNIFICANT & MEANINGFUL (d >= 0.2). Generating␣
↪Plot...")
                    plt.figure(figsize=(10, 6))
                    sns.violinplot(x='type', y=feat, data=data,␣
↪palette='muted', inner='quartile')
                    plt.title(f'{feat}: Known vs Novel (p={p_val:.2e},␣
↪d={cohens_d:.2f})')
                    plt.show()
                elif p_val < 0.05:
                     print(f"-> Statistically Significant but NEGLIGIBLE Effect␣
↪(d < 0.2). Plot suppressed.")
                else:
                    print(f"-> Not Significant.")
        else:
            pass

    # 3. Boolean/Categorical Feature Analysis
    for feat in boolean_features:
        if feat in df_targets.columns:
            # Ensure boolean
            df_targets[feat] = df_targets[feat].fillna(False).astype(bool)

            contingency = pd.crosstab(df_targets['type'], df_targets[feat])

            if contingency.size > 0:
                try:
                    # PRINT DESCRIPTION
                    desc = FEATURE_DESCRIPTIONS.get(feat, "")
                    if desc:
                        print(f"\n[?] {feat}: {desc}")

                    chi2, p_val, dof, ex = stats.chi2_contingency(contingency)

                    # Calculate Proportions difference (Effect Size for Binary)
```

```
                    props = df_targets.groupby('type')[feat].mean() # Mean of␣
↪boolean is proportion
                    if 'Known Positive' in props and 'Novel Prediction' in␣
↪props:
                        prop_diff = props['Known Positive'] - props['Novel␣
↪Prediction']
                    else:
                        prop_diff = 0

                    print(f"Categorical Feature '{feat}':")
                    print(f"  P-value    = {p_val:.4e}")
                    print(f"  Prop Diff = {prop_diff:.2%}")

                    # Threshold: P < 0.05 AND at least 5% difference in␣
↪prevalence
                    if p_val < 0.05 and abs(prop_diff) > 0.01: # 1% threshold␣
↪for boolean
                        print(f"-> SIGNIFICANT. Generating Plot...")
                        prop_data = df_targets.groupby('type')[feat].
↪value_counts(normalize=True).rename('proportion').reset_index()
                        prop_true = prop_data[prop_data[feat] == True]

                        if not prop_true.empty:
                            plt.figure(figsize=(8, 6))
                            sns.barplot(x='type', y='proportion',␣
↪data=prop_true, palette='pastel')
                            plt.title(f'Proportion of Targets with {feat}␣
↪(True)\n(p={p_val:.2e})')
                            plt.ylabel(f'Proportion ({feat}=True)')
                            plt.show()
                    else:
                         print(f"-> Not Significant or Negligible Difference.")

                except Exception as e:
                    print(f"Could not run Chi-Square on {feat}: {e}")


def analyze_target_properties_deduplicated(df):
    """
    Aggregates data to 1 row per target to analyze inherent properties.
    """
    print("\n--- Target Level Analysis (Unique Targets - General Stats) ---")

    # Deduplicate by targetId (keep highest score prediction)
    df_targets = df.sort_values('score', ascending=False).
↪drop_duplicates(subset=['targetId'])
```

```
    # A. Target Class
    if 'targetClass' in df_targets.columns:
        # Explode first (targets may have multiple classes)
        df_exploded = df_targets.explode('targetClass')
        # CLEAN the labels
        df_exploded['targetClass_clean'] = df_exploded['targetClass'].
↪apply(extract_clean_label)

        top_classes = df_exploded['targetClass_clean'].value_counts().head(10)

        plt.figure(figsize=(12, 6))
        sns.barplot(x=top_classes.values, y=top_classes.index, palette='magma')
        plt.title('Top Target Classes (Unique Targets)')
        plt.show()

    # B. Clinical Phase
    if 'maxClinicalTrialPhase' in df_targets.columns:
        plt.figure(figsize=(10, 6))
        sns.countplot(x='maxClinicalTrialPhase', data=df_targets,␣
↪palette='viridis')
        plt.title('Clinical Phase Distribution (Unique Targets)')
        plt.show()

def analyze_therapeutic_areas(df):
    print("\n--- Therapeutic Area Analysis ---")
    if 'therapeuticAreas' in df.columns:
        df_exploded = df.explode('therapeuticAreas')
        top_tas = df_exploded['therapeuticAreas'].value_counts().head(10)

        plt.figure(figsize=(12, 6))
        sns.barplot(x=top_tas.values, y=top_tas.index, palette='viridis')
        plt.title('Top 10 Therapeutic Areas')
        plt.show()

def analyze_repurposing_potential(df):
    print("\n--- Repurposing Potential ---")
    if 'maxClinicalTrialPhase' in df.columns:
        repurposed = df[(df['score'] > 0.8) & (df['maxClinicalTrialPhase'] >=␣
↪4)]
        print(f"High confidence (score > 0.8) candidates using Approved Drugs:␣
↪{len(repurposed)}")
        if not repurposed.empty:
            cols = ['diseaseName', 'targetSymbol', 'score']
            print(repurposed[cols].head(10).to_string(index=False))

def analyze_target_classes(df):
```

```
    """
    Analyzes score distribution by Target Class (All predictions).
    CRITICAL: Handles dictionary/complex objects in targetClass.
    """
    print("\n--- Target Class Volume (All Predictions) ---")
    if 'targetClass' in df.columns:
        # 1. Explode to handle lists
        df_exploded = df.explode('targetClass')

        # 2. Extract clean strings from dictionaries
        df_exploded['targetClass_clean'] = df_exploded['targetClass'].
 ↪apply(extract_clean_label)

        # 3. Filter Top 10
        top_classes = df_exploded['targetClass_clean'].value_counts().head(10).
 ↪index
        df_filtered = df_exploded[df_exploded['targetClass_clean'].
 ↪isin(top_classes)]

        # 4. Plot using the CLEAN column
        plt.figure(figsize=(12, 6))
        sns.boxplot(y='targetClass_clean', x='score', data=df_filtered,␣
 ↪order=top_classes, palette='coolwarm')
        plt.title('Score Distribution by Target Class')
        plt.ylabel('Target Class')
        plt.xlabel('Prediction Score')
        plt.show()

def main():
    # 1. Load FULL Data (Needed for comparison)
    df = load_and_merge_data()

    # 2. Filter for NOVEL predictions (label < 1) for specific analyses
    print("\n--- Filtering for Novel Predictions (label < 1) ---")
    df_novel = df.query("label < 1").copy()
    print(f"Novel Predictions Count: {len(df_novel)}")

    if len(df_novel) == 0:
        print("Warning: No novel predictions found. Check filtering logic.")

    # 3. Run Comparison (Known vs Novel) on FULL dataframe
    analyze_target_prioritization_features(df)

    # 4. Disease/Prediction Level Analysis (Novel Only)
    analyze_rarity(df_novel)
    analyze_omim(df_novel)
    analyze_orphan(df_novel)
```

```python
    analyze_score_distributions(df_novel)

    # 5. Target/Biology Level Analysis (Novel Only)
    analyze_therapeutic_areas(df_novel)
    analyze_repurposing_potential(df_novel)
    analyze_target_classes(df_novel)

    # 6. Strict Deduplicated Analysis (Novel Only)
    analyze_target_properties_deduplicated(df_novel)

if __name__ == "__main__":
    main()
```

```
Loading data…
Merging metadata…
Data merged successfully. Shape: (190644, 57)
Split counts:
type
Novel Prediction    123112
Known Positive       67532
Name: count, dtype: int64

--- Filtering for Novel Predictions (label < 1) ---
Novel Predictions Count: 123112

--- Target Prioritization Feature Analysis (Known vs Novel) ---
Total Unique Targets: 2613
Breakdown by Type:
type
Known Positive      1518
Novel Prediction    1095
Name: count, dtype: int64

[?] geneticConstraint: Represents the genetic constraint of the target,
indicating how intolerant it is to genetic variation
Numeric Feature 'geneticConstraint':
  P-value    = 3.3015e-19
  Cohen's d  = -0.3700 (Effect Size)
  Median Diff= -0.3006
-> SIGNIFICANT & MEANINGFUL (d >= 0.2). Generating Plot…
```

```
/tmp/ipykernel_903/1865608078.py:313: FutureWarning:

Passing `palette` without assigning `hue` is deprecated and will be removed in
v0.14.0. Assign the `x` variable to `hue` and set `legend=False` for the same
effect.

  sns.violinplot(x='type', y=feat, data=data, palette='muted', inner='quartile')
```

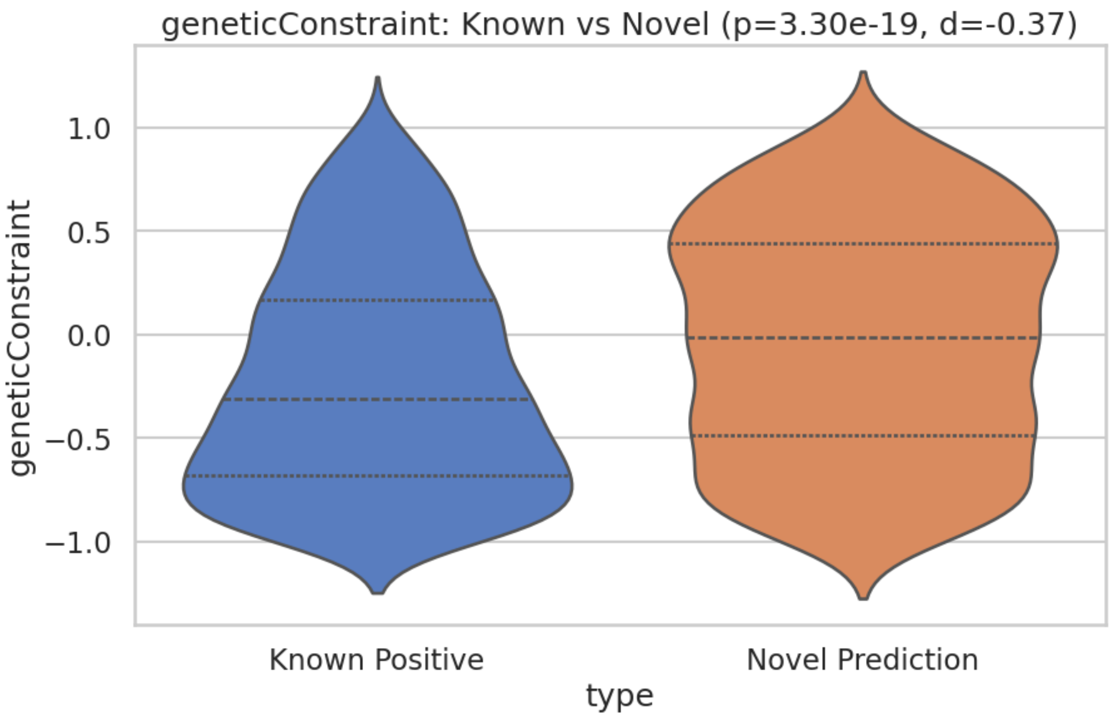


```
/tmp/ipykernel_903/1865608078.py:313: FutureWarning:

Passing `palette` without assigning `hue` is deprecated and will be removed in
v0.14.0. Assign the `x` variable to `hue` and set `legend=False` for the same
effect.

  sns.violinplot(x='type', y=feat, data=data, palette='muted', inner='quartile')


[?] paralogMaxIdentityPercentage: Maximum percentage identity among the target's
paralogues
Numeric Feature 'paralogMaxIdentityPercentage':
  P-value    = 6.3075e-01
  Cohen's d  = 0.0541 (Effect Size)
  Median Diff= 0.0000
-> Not Significant.

[?] mouseOrthologMaxIdentityPercentage: Maximum percentage identity between the
target and its mouse ortholog
Numeric Feature 'mouseOrthologMaxIdentityPercentage':
  P-value    = 2.3965e-05
  Cohen's d  = 0.1758 (Effect Size)
  Median Diff= 0.0871
-> Statistically Significant but NEGLIGIBLE Effect (d < 0.2). Plot suppressed.
```

```
[?] mouseKOScore: Represents the phenotypic score from mouse knockout models for
the target
Numeric Feature 'mouseKOScore':
  P-value    = 2.4033e-15
  Cohen's d  = -0.3302 (Effect Size)
  Median Diff= -0.2753
-> SIGNIFICANT & MEANINGFUL (d >= 0.2). Generating Plot…
```

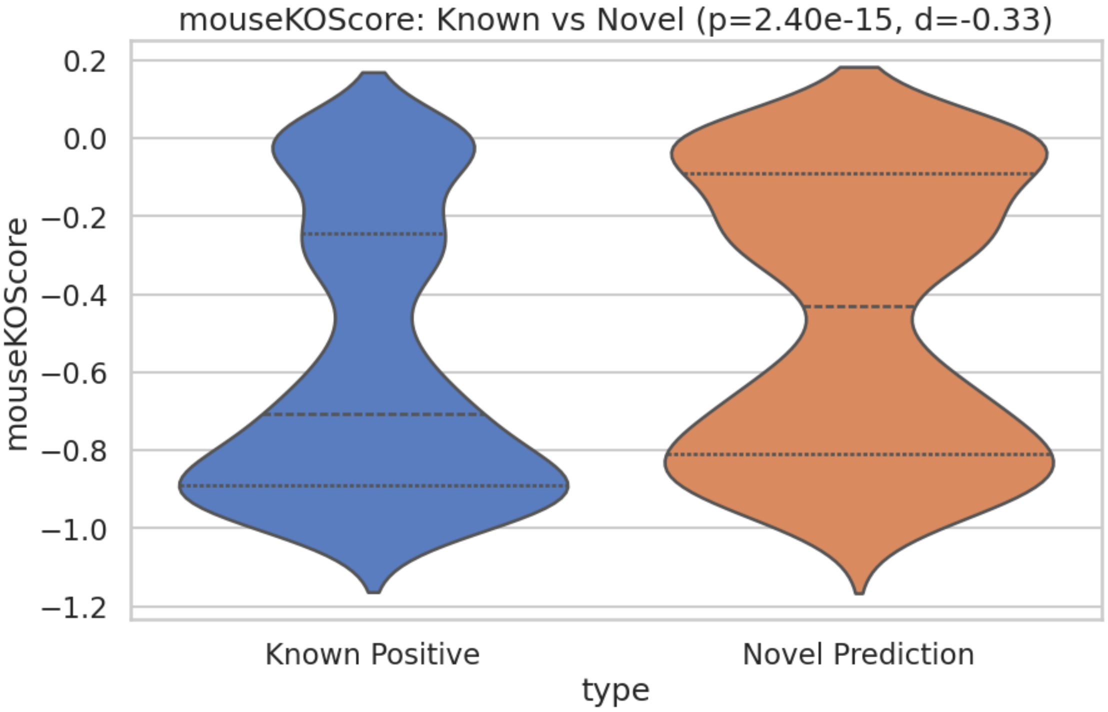


```
[?] maxClinicalTrialPhase: Highest clinical trial phase that the target has
reached for any indication
Numeric Feature 'maxClinicalTrialPhase':
  P-value    = 6.9048e-02
  Cohen's d  = 0.7113 (Effect Size)
  Median Diff= 0.2500
-> Not Significant.

[?] tissueSpecificity: Describes the specificity of the target's expression
across different tissues
Numeric Feature 'tissueSpecificity':
  P-value    = 2.3787e-05
  Cohen's d  = -0.1769 (Effect Size)
  Median Diff= 0.0000
-> Statistically Significant but NEGLIGIBLE Effect (d < 0.2). Plot suppressed.
```

```
[?] tissueDistribution: Describes the distribution pattern of the target's
expression in various tissues
Numeric Feature 'tissueDistribution':
  P-value    = 7.5068e-07
  Cohen's d  = -0.1947 (Effect Size)
  Median Diff= 0.0000
-> Statistically Significant but NEGLIGIBLE Effect (d < 0.2). Plot suppressed.
Numeric Feature 'count_synonyms':
  P-value    = 1.0000e+00
  Cohen's d  = nan (Effect Size)
  Median Diff= 0.0000
-> Not Significant.
Numeric Feature 'count_subcellularLocations':
  P-value    = 1.6846e-08
  Cohen's d  = 0.2242 (Effect Size)
  Median Diff= 1.0000
-> SIGNIFICANT & MEANINGFUL (d >= 0.2). Generating Plot…
```

```
/tmp/ipykernel_903/1865608078.py:74: RuntimeWarning: invalid value encountered
in scalar divide
  return diff / pooled_sd
/tmp/ipykernel_903/1865608078.py:313: FutureWarning:

Passing `palette` without assigning `hue` is deprecated and will be removed in
v0.14.0. Assign the `x` variable to `hue` and set `legend=False` for the same
effect.

  sns.violinplot(x='type', y=feat, data=data, palette='muted', inner='quartile')
```

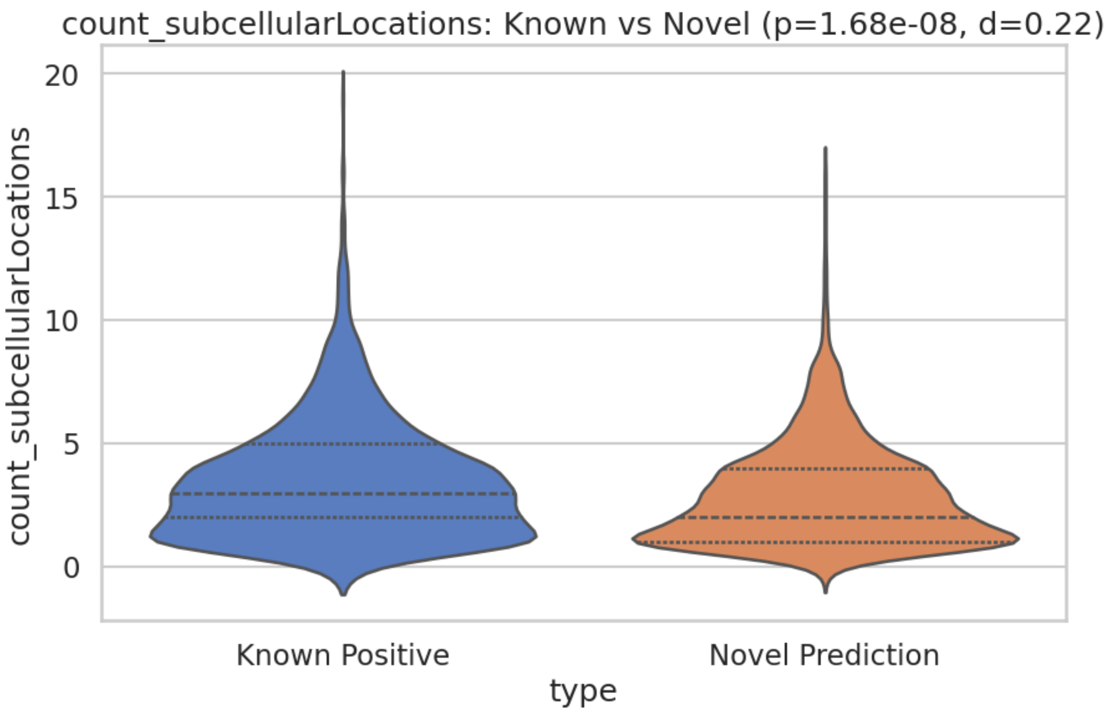


```
Numeric Feature 'count_go':
  P-value    = 4.5630e-54
  Cohen's d  = 0.5492 (Effect Size)
  Median Diff= 9.0000
-> SIGNIFICANT & MEANINGFUL (d >= 0.2). Generating Plot…
```

```
/tmp/ipykernel_903/1865608078.py:313: FutureWarning:

Passing `palette` without assigning `hue` is deprecated and will be removed in
v0.14.0. Assign the `x` variable to `hue` and set `legend=False` for the same
effect.

  sns.violinplot(x='type', y=feat, data=data, palette='muted', inner='quartile')
```

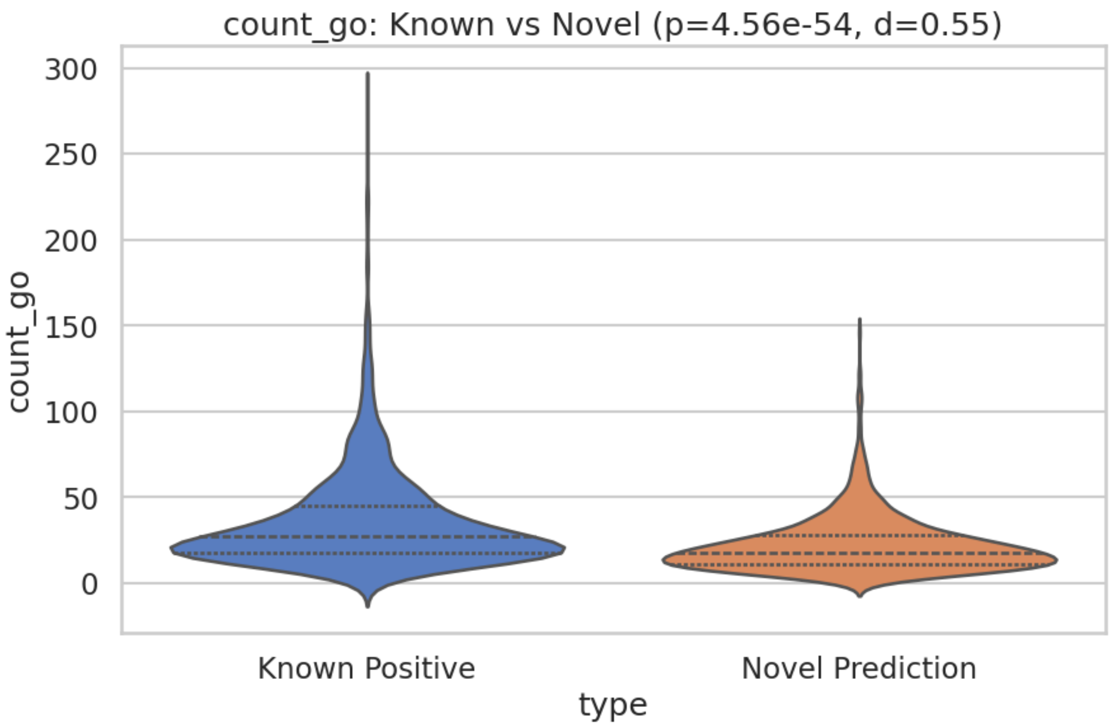


```
Numeric Feature 'count_pathways':
  P-value    = 7.1656e-50
  Cohen's d  = 0.3905 (Effect Size)
  Median Diff= 1.0000
-> SIGNIFICANT & MEANINGFUL (d >= 0.2). Generating Plot…
```

```
/tmp/ipykernel_903/1865608078.py:313: FutureWarning:

Passing `palette` without assigning `hue` is deprecated and will be removed in
v0.14.0. Assign the `x` variable to `hue` and set `legend=False` for the same
effect.

  sns.violinplot(x='type', y=feat, data=data, palette='muted', inner='quartile')
```

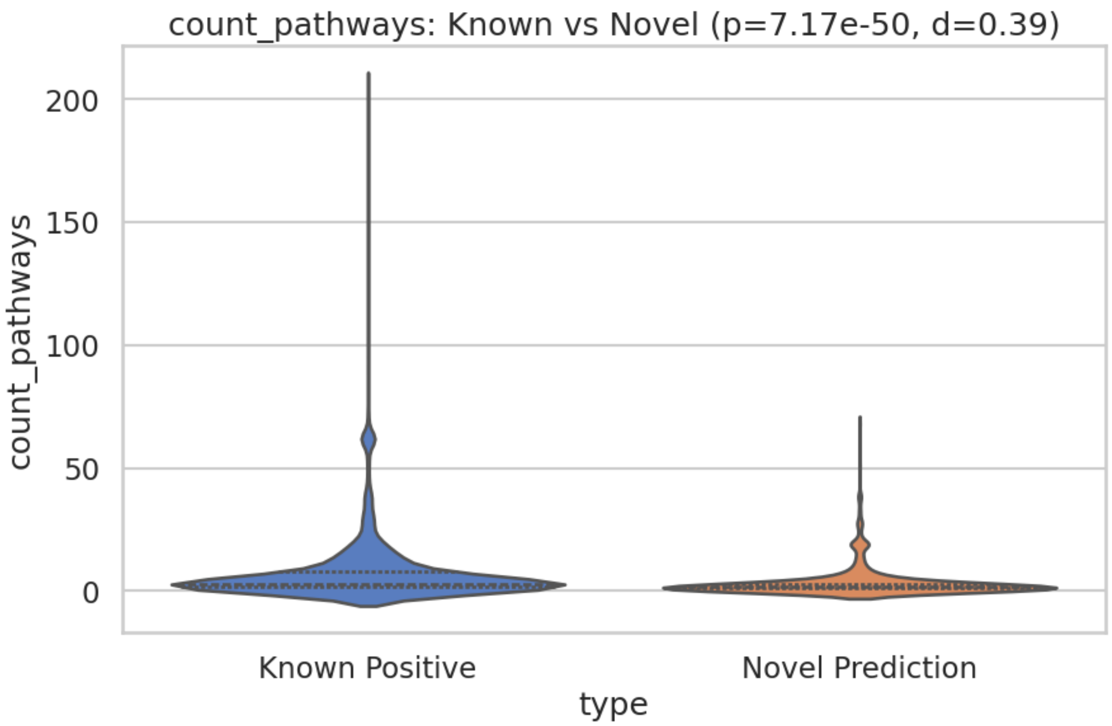


```
Numeric Feature 'count_proteinIds':
  P-value    = 8.2174e-11
  Cohen's d  = 0.1861 (Effect Size)
  Median Diff= 3.0000
-> Statistically Significant but NEGLIGIBLE Effect (d < 0.2). Plot suppressed.
Numeric Feature 'count_targetClass':
  P-value    = 3.0765e-52
  Cohen's d  = 0.3926 (Effect Size)
  Median Diff= 1.0000
-> SIGNIFICANT & MEANINGFUL (d >= 0.2). Generating Plot…
```

```
/tmp/ipykernel_903/1865608078.py:313: FutureWarning:

Passing `palette` without assigning `hue` is deprecated and will be removed in
v0.14.0. Assign the `x` variable to `hue` and set `legend=False` for the same
effect.

  sns.violinplot(x='type', y=feat, data=data, palette='muted', inner='quartile')
```

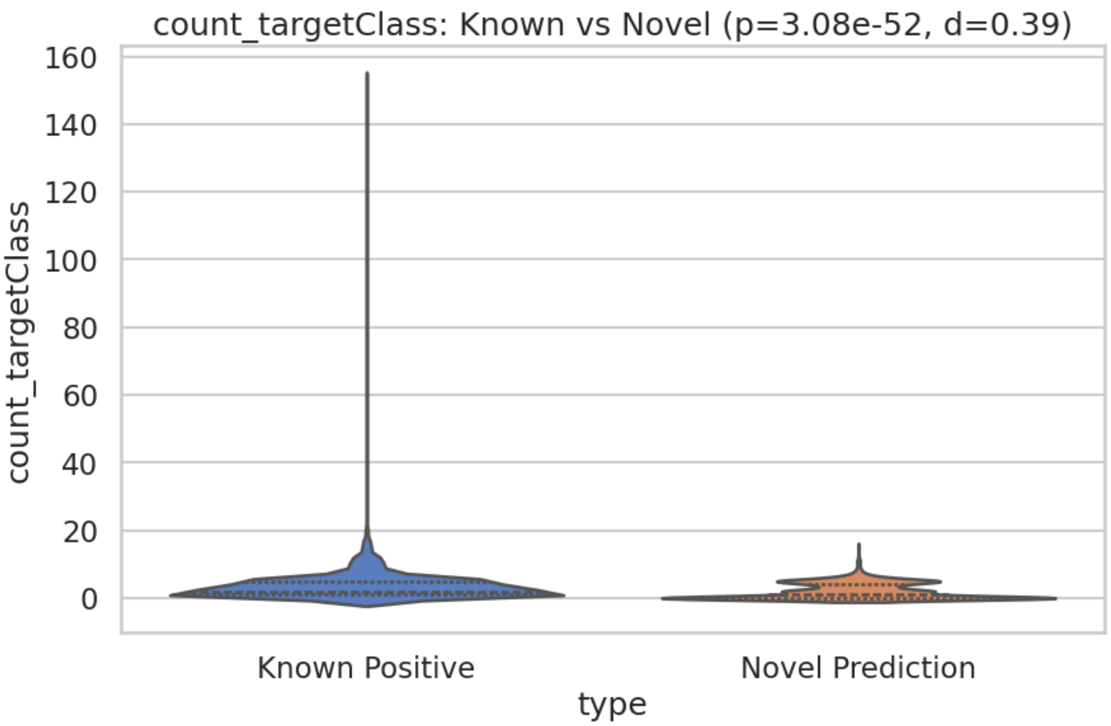


```
Numeric Feature 'count_safetyLiabilities':
  P-value    = 8.0753e-27
  Cohen's d  = 0.2839 (Effect Size)
  Median Diff= 0.0000
-> SIGNIFICANT & MEANINGFUL (d >= 0.2). Generating Plot…
```

```
/tmp/ipykernel_903/1865608078.py:313: FutureWarning:

Passing `palette` without assigning `hue` is deprecated and will be removed in
v0.14.0. Assign the `x` variable to `hue` and set `legend=False` for the same
effect.

  sns.violinplot(x='type', y=feat, data=data, palette='muted', inner='quartile')
```

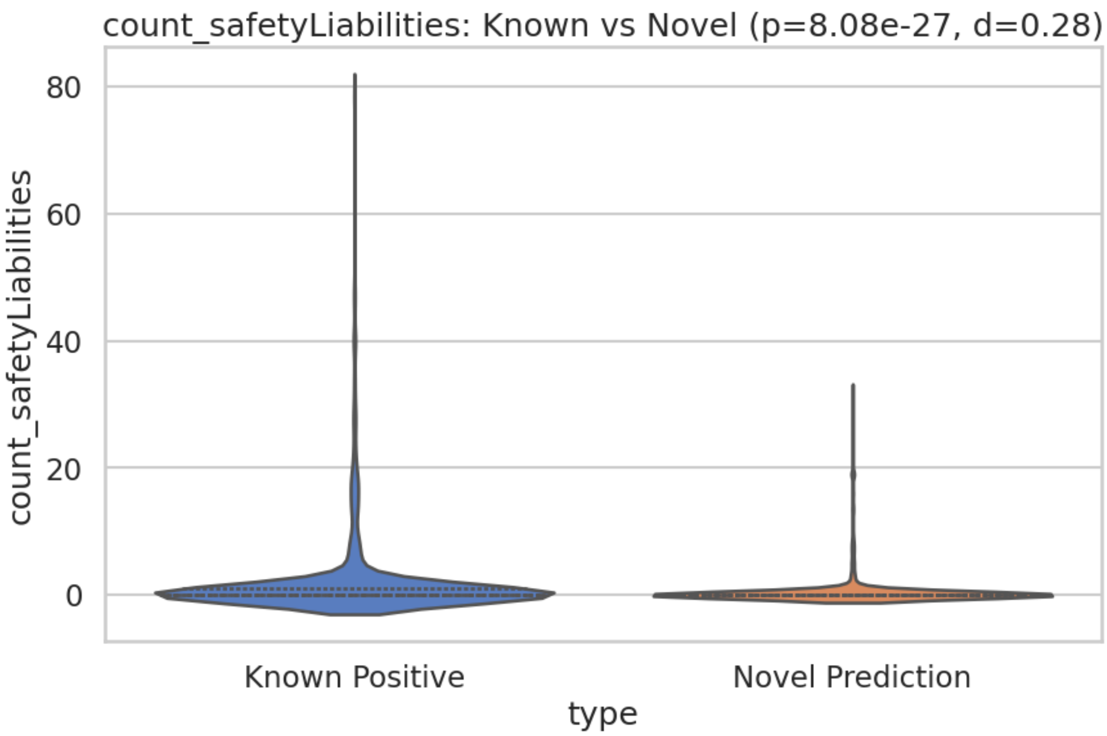


```
Numeric Feature 'count_tractability':
  P-value    = 7.9296e-140
  Cohen's d  = 1.0855 (Effect Size)
  Median Diff= 3.0000
-> SIGNIFICANT & MEANINGFUL (d >= 0.2). Generating Plot…
```

```
/tmp/ipykernel_903/1865608078.py:313: FutureWarning:

Passing `palette` without assigning `hue` is deprecated and will be removed in
v0.14.0. Assign the `x` variable to `hue` and set `legend=False` for the same
effect.

  sns.violinplot(x='type', y=feat, data=data, palette='muted', inner='quartile')
```

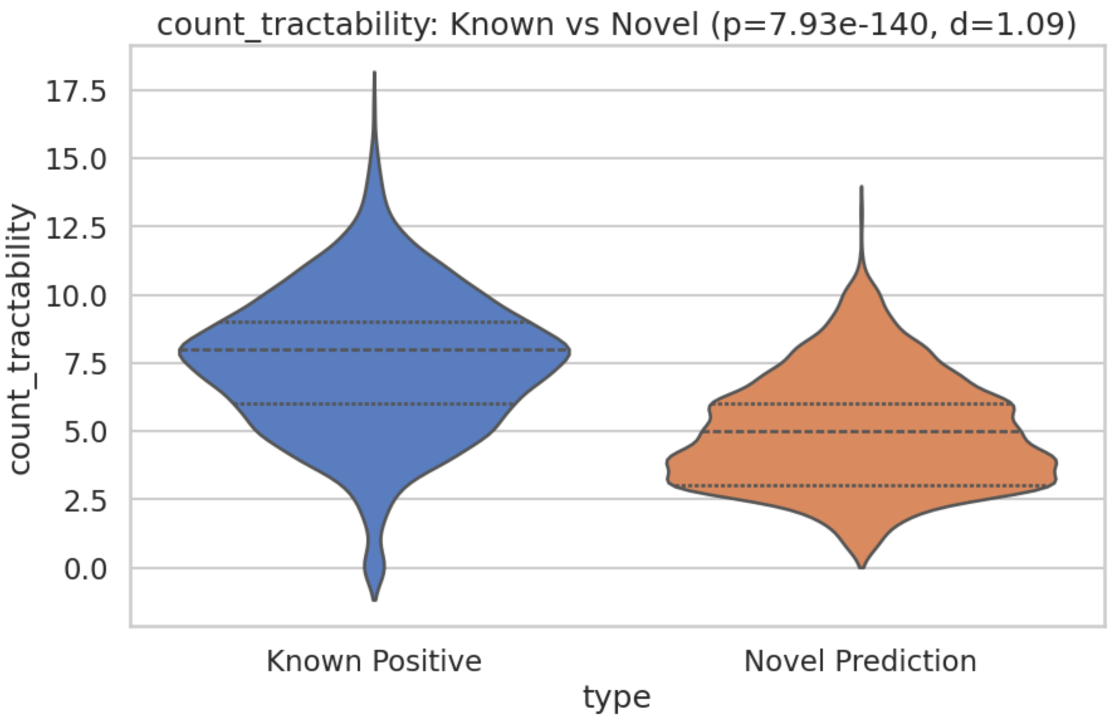


```
Numeric Feature 'count_alternativeGenes':
  P-value    = 2.8961e-01
  Cohen's d  = 0.0213 (Effect Size)
  Median Diff= 0.0000
-> Not Significant.
Numeric Feature 'count_tep':
  P-value    = 6.0793e-02
  Cohen's d  = 0.0744 (Effect Size)
  Median Diff= 0.0000
-> Not Significant.

[?] score_syn: Synonymous intolerance score (Z-score likely)
Numeric Feature 'score_syn':
  P-value    = 3.0070e-03
  Cohen's d  = -0.0804 (Effect Size)
  Median Diff= -0.1424
-> Statistically Significant but NEGLIGIBLE Effect (d < 0.2). Plot suppressed.

[?] score_mis: Missense intolerance score (Z-score likely)
Numeric Feature 'score_mis':
  P-value    = 3.0892e-14
  Cohen's d  = 0.2885 (Effect Size)
  Median Diff= 0.4442
-> SIGNIFICANT & MEANINGFUL (d >= 0.2). Generating Plot…
```

```
/tmp/ipykernel_903/1865608078.py:313: FutureWarning:

Passing `palette` without assigning `hue` is deprecated and will be removed in
v0.14.0. Assign the `x` variable to `hue` and set `legend=False` for the same
effect.

  sns.violinplot(x='type', y=feat, data=data, palette='muted', inner='quartile')
```

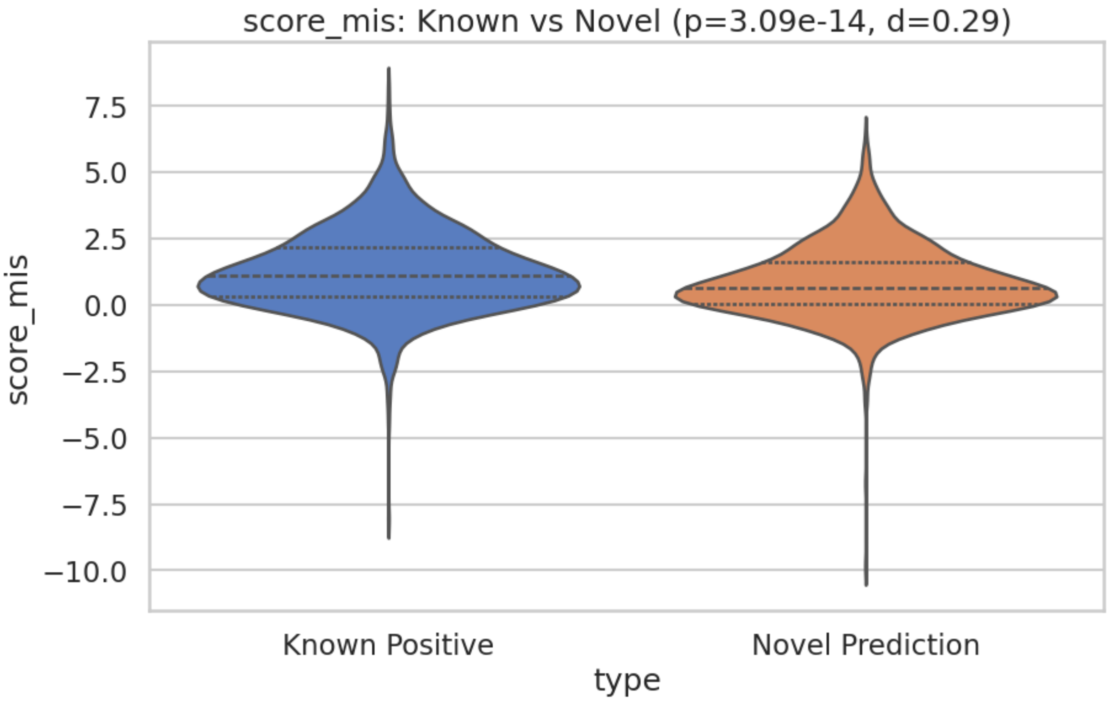


```
[?] score_lof: Loss-of-function intolerance score (pLI/LOEUF likely)
Numeric Feature 'score_lof':
  P-value    = 9.1869e-16
  Cohen's d  = 0.3043 (Effect Size)
  Median Diff= 0.0305
-> SIGNIFICANT & MEANINGFUL (d >= 0.2). Generating Plot…

/tmp/ipykernel_903/1865608078.py:313: FutureWarning:

Passing `palette` without assigning `hue` is deprecated and will be removed in
v0.14.0. Assign the `x` variable to `hue` and set `legend=False` for the same
effect.

  sns.violinplot(x='type', y=feat, data=data, palette='muted', inner='quartile')
```

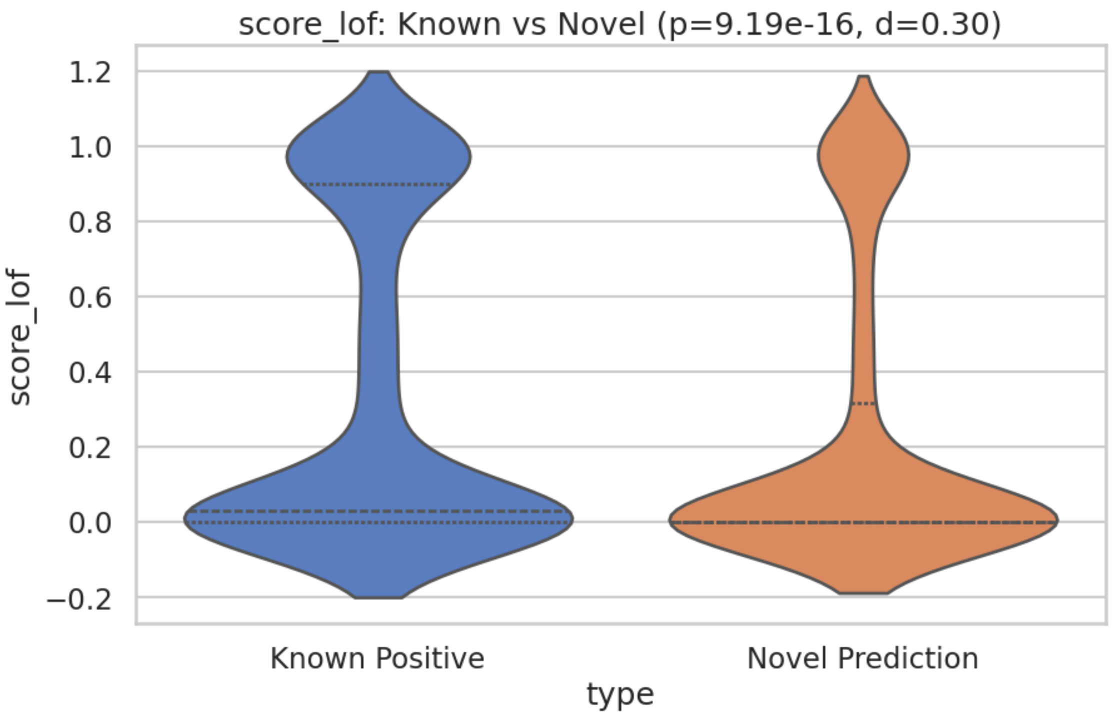


```
[?] isCancerDriverGene: Indicates if the target is identified as a cancer driver
gene
Categorical Feature 'isCancerDriverGene':
  P-value   = 1.0153e-08
  Prop Diff = 4.60%
-> SIGNIFICANT. Generating Plot…
```

```
/tmp/ipykernel_903/1865608078.py:359: FutureWarning:

Passing `palette` without assigning `hue` is deprecated and will be removed in
v0.14.0. Assign the `x` variable to `hue` and set `legend=False` for the same
effect.

  sns.barplot(x='type', y='proportion', data=prop_true, palette='pastel')
```

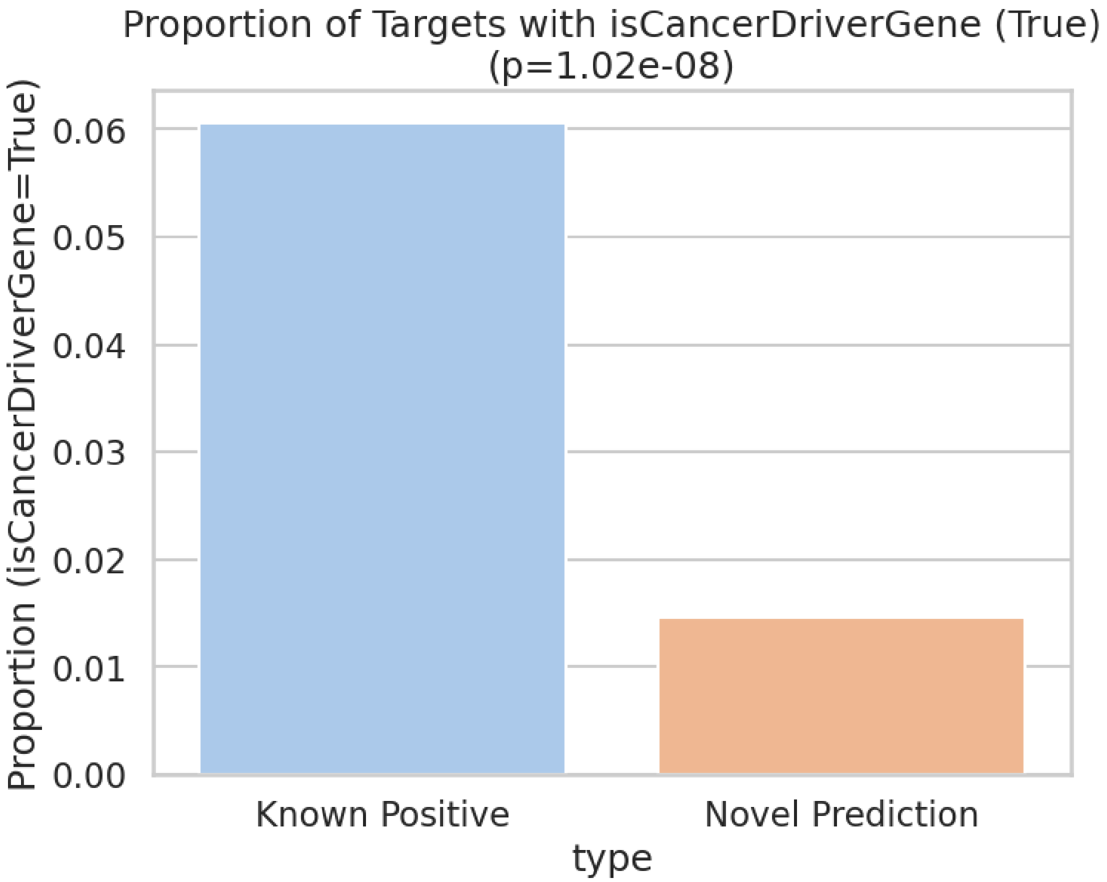


```
[?] hasTEP: Indicates if a Target Enabling Package (TEP) is available for the
target
Categorical Feature 'hasTEP':
  P-value   = 1.2423e-01
  Prop Diff = 0.44%
-> Not Significant or Negligible Difference.

[?] hasHighQualityChemicalProbes: Indicates if there are high-quality chemical
probes available for the target
Categorical Feature 'hasHighQualityChemicalProbes':
  P-value   = 1.8811e-20
  Prop Diff = 13.02%
-> SIGNIFICANT. Generating Plot…
```

```
/tmp/ipykernel_903/1865608078.py:359: FutureWarning:

Passing `palette` without assigning `hue` is deprecated and will be removed in
v0.14.0. Assign the `x` variable to `hue` and set `legend=False` for the same
effect.
```

```
  sns.barplot(x='type', y='proportion', data=prop_true, palette='pastel')
```

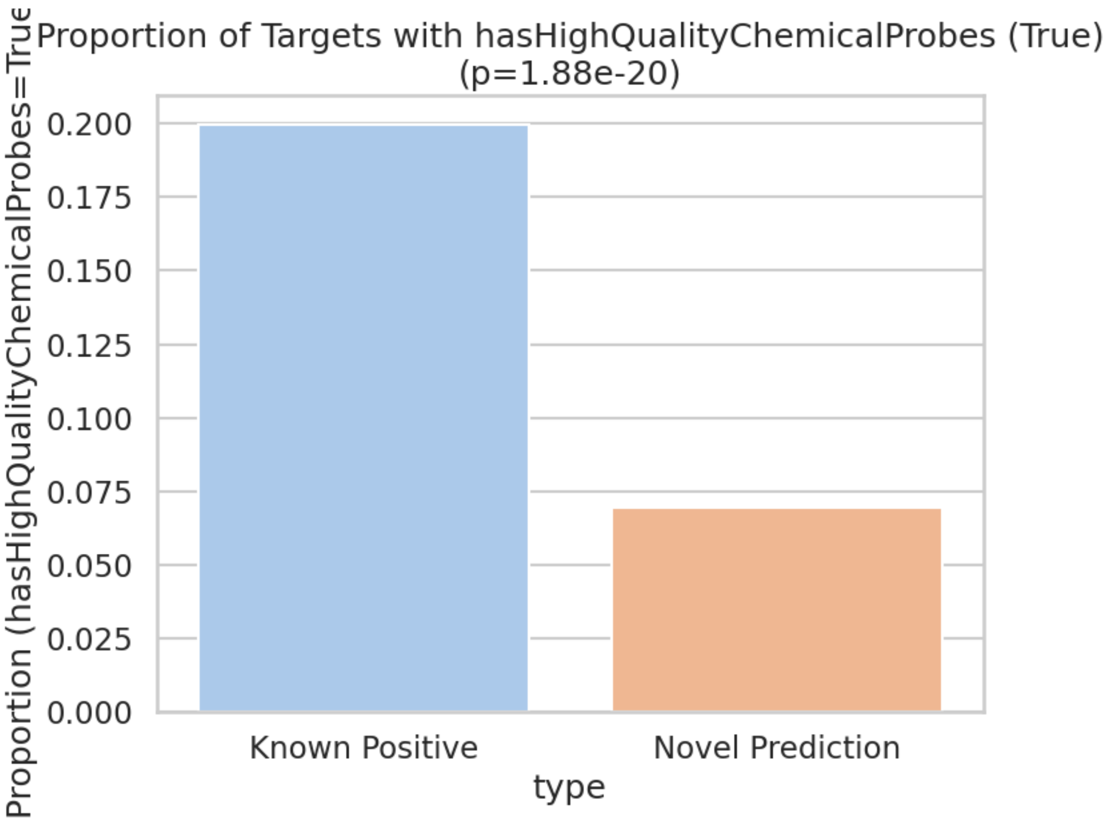


```
[?] isInMembrane: Indicates if the target protein is located in the cell or
plasma membrane
Categorical Feature 'isInMembrane':
  P-value   = 2.9521e-01
  Prop Diff = 2.07%
-> Not Significant or Negligible Difference.

[?] isSecreted: Indicates if the target protein is secreted or predicted to be
secreted
Categorical Feature 'isSecreted':
  P-value   = 2.0724e-07
  Prop Diff = -8.39%
-> SIGNIFICANT. Generating Plot…
```

```
/tmp/ipykernel_903/1865608078.py:359: FutureWarning:

Passing `palette` without assigning `hue` is deprecated and will be removed in
v0.14.0. Assign the `x` variable to `hue` and set `legend=False` for the same
effect.
```

```
sns.barplot(x='type', y='proportion', data=prop_true, palette='pastel')
```

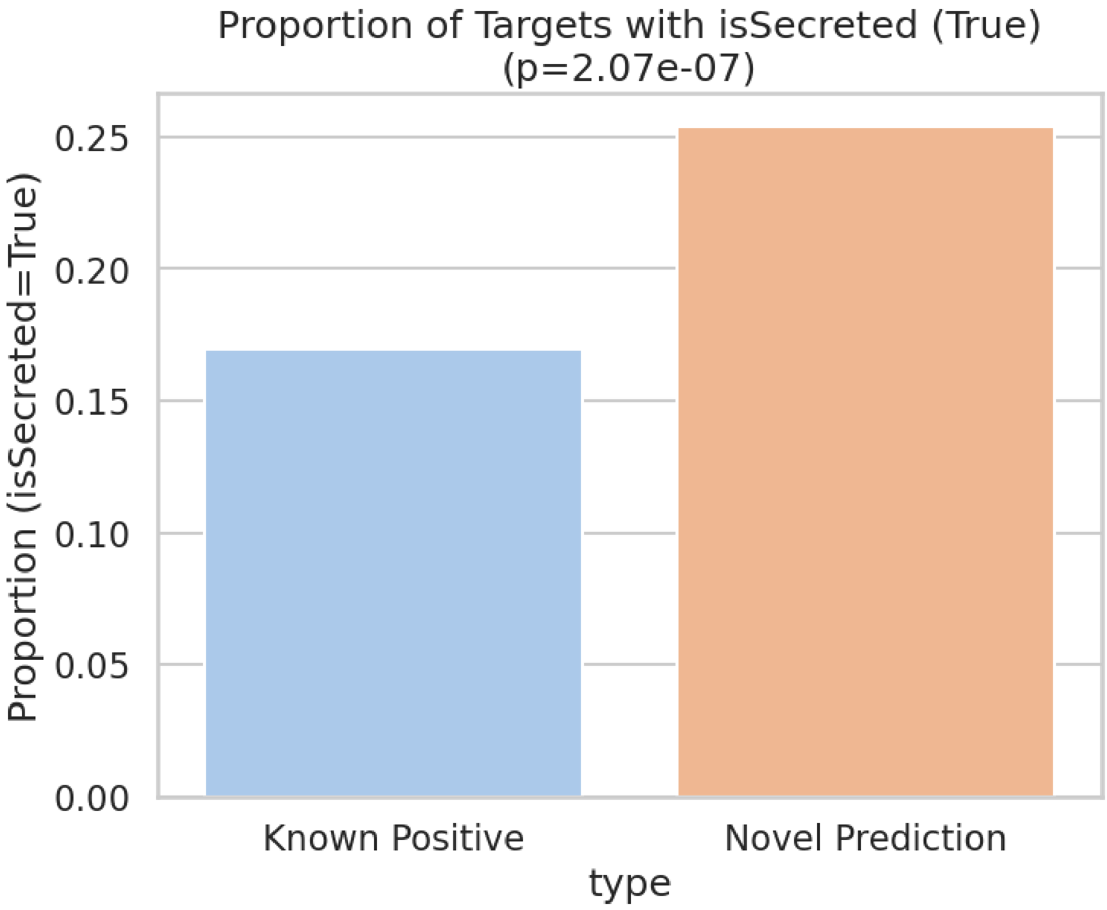


```
[?] hasSafetyEvent: Indicates if there are known safety events associated with
the target
Categorical Feature 'hasSafetyEvent':
  P-value   = 4.4563e-26
  Prop Diff = 16.73%
-> SIGNIFICANT. Generating Plot…
```

```
/tmp/ipykernel_903/1865608078.py:359: FutureWarning:

Passing `palette` without assigning `hue` is deprecated and will be removed in
v0.14.0. Assign the `x` variable to `hue` and set `legend=False` for the same
effect.

  sns.barplot(x='type', y='proportion', data=prop_true, palette='pastel')
```

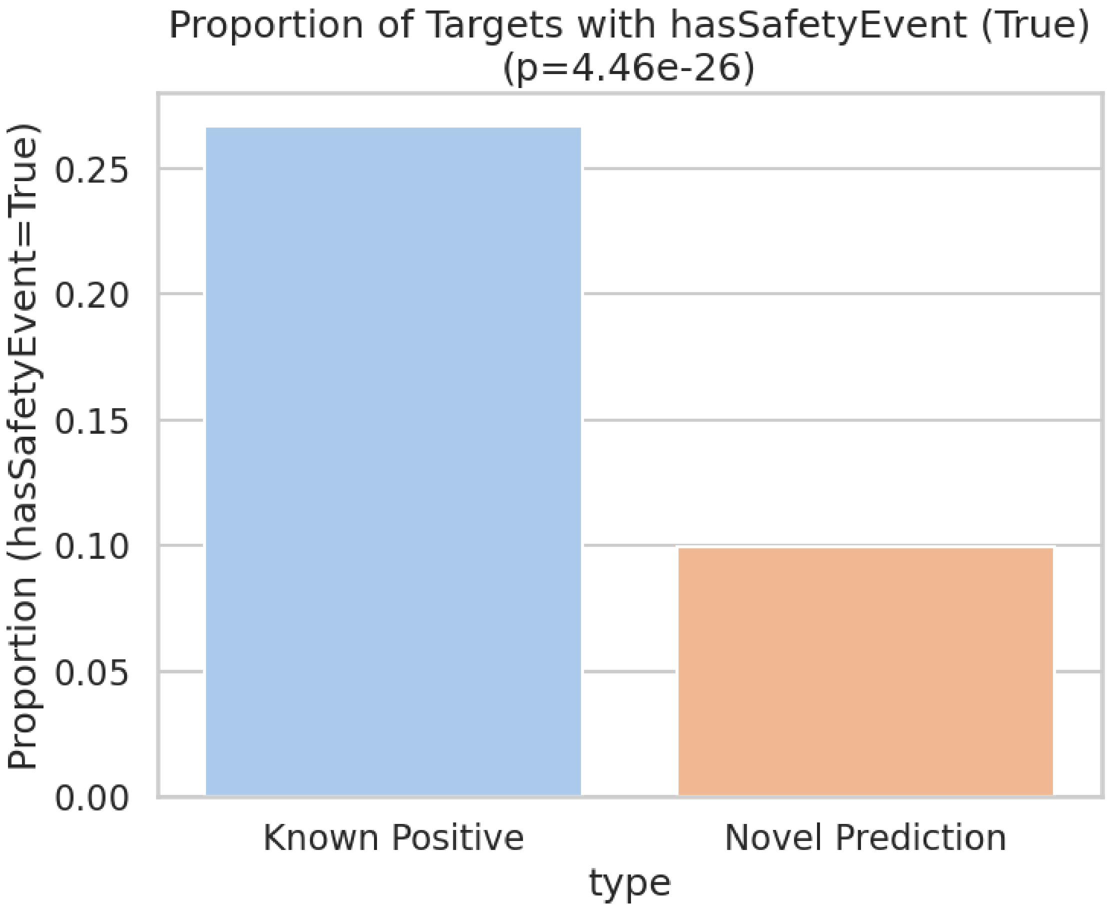


```
[?] hasPocket: Indicates if the target has predicted binding pockets suitable
for small molecule binding
Categorical Feature 'hasPocket':
  P-value   = 5.6725e-11
  Prop Diff = 9.17%
-> SIGNIFICANT. Generating Plot…
```

```
/tmp/ipykernel_903/1865608078.py:359: FutureWarning:

Passing `palette` without assigning `hue` is deprecated and will be removed in
v0.14.0. Assign the `x` variable to `hue` and set `legend=False` for the same
effect.

  sns.barplot(x='type', y='proportion', data=prop_true, palette='pastel')
```

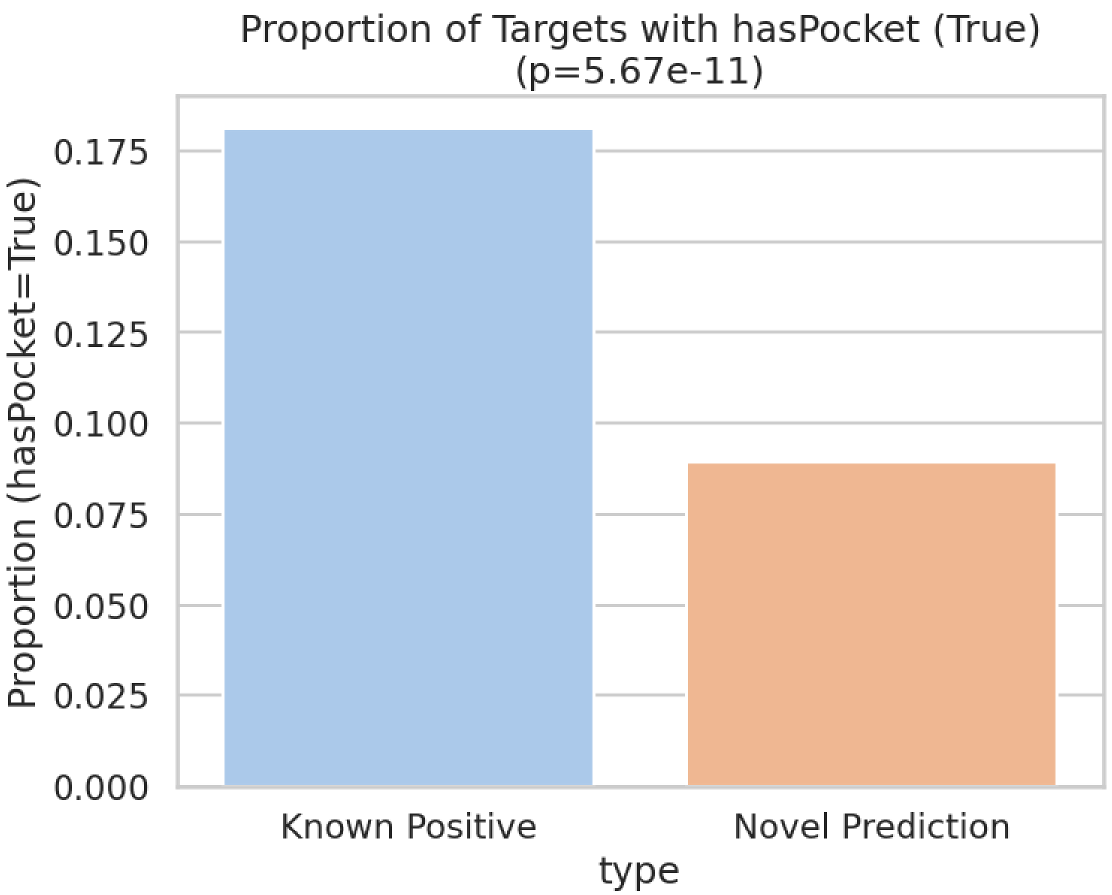


```
[?] hasLigand: Indicates if the target binds at least one high-quality ligand
Categorical Feature 'hasLigand':
  P-value   = 2.9892e-52
  Prop Diff = 30.15%
-> SIGNIFICANT. Generating Plot…
```

```
/tmp/ipykernel_903/1865608078.py:359: FutureWarning:

Passing `palette` without assigning `hue` is deprecated and will be removed in
v0.14.0. Assign the `x` variable to `hue` and set `legend=False` for the same
effect.

  sns.barplot(x='type', y='proportion', data=prop_true, palette='pastel')
```

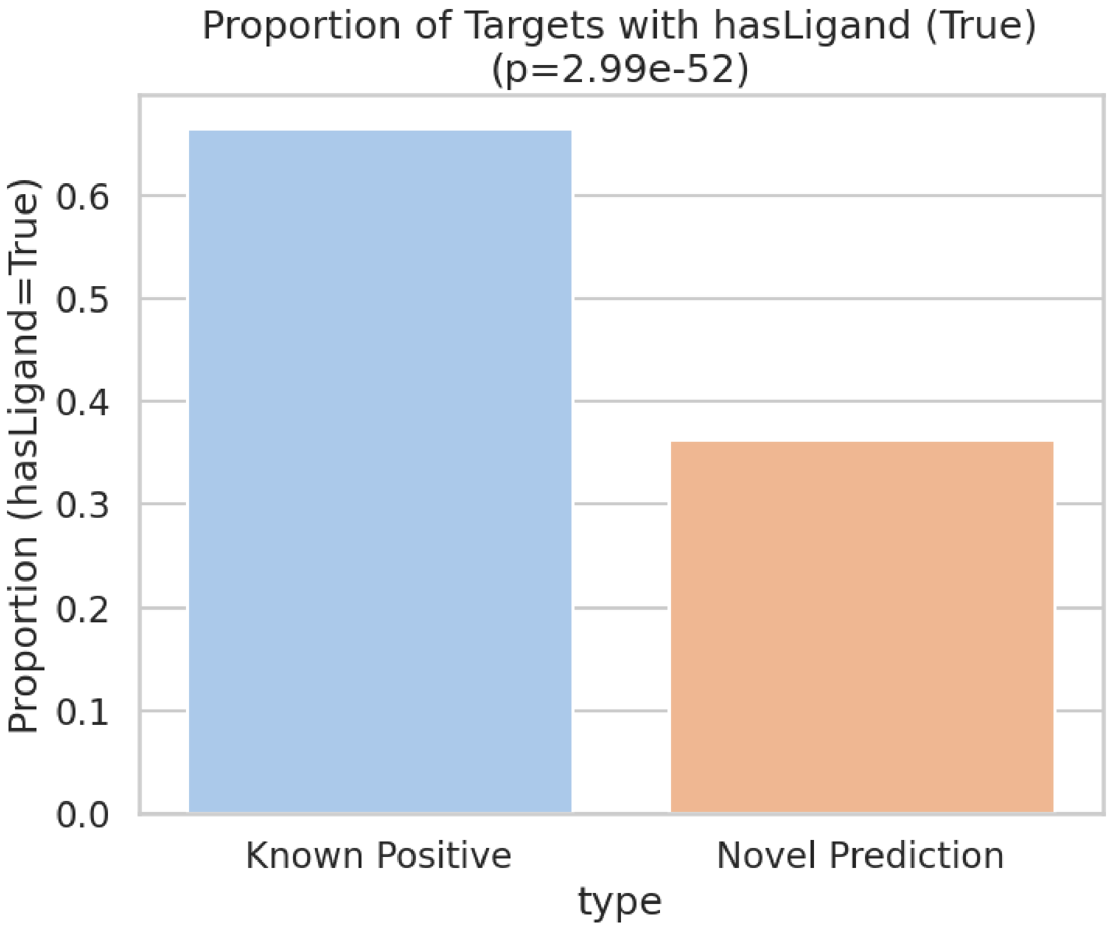


```
[?] hasSmallMoleculeBinder: Indicates if the target has at least one small
molecule binder
Categorical Feature 'hasSmallMoleculeBinder':
  P-value   = 1.8588e-37
  Prop Diff = 24.88%
-> SIGNIFICANT. Generating Plot…
```

```
/tmp/ipykernel_903/1865608078.py:359: FutureWarning:

Passing `palette` without assigning `hue` is deprecated and will be removed in
v0.14.0. Assign the `x` variable to `hue` and set `legend=False` for the same
effect.

  sns.barplot(x='type', y='proportion', data=prop_true, palette='pastel')
```

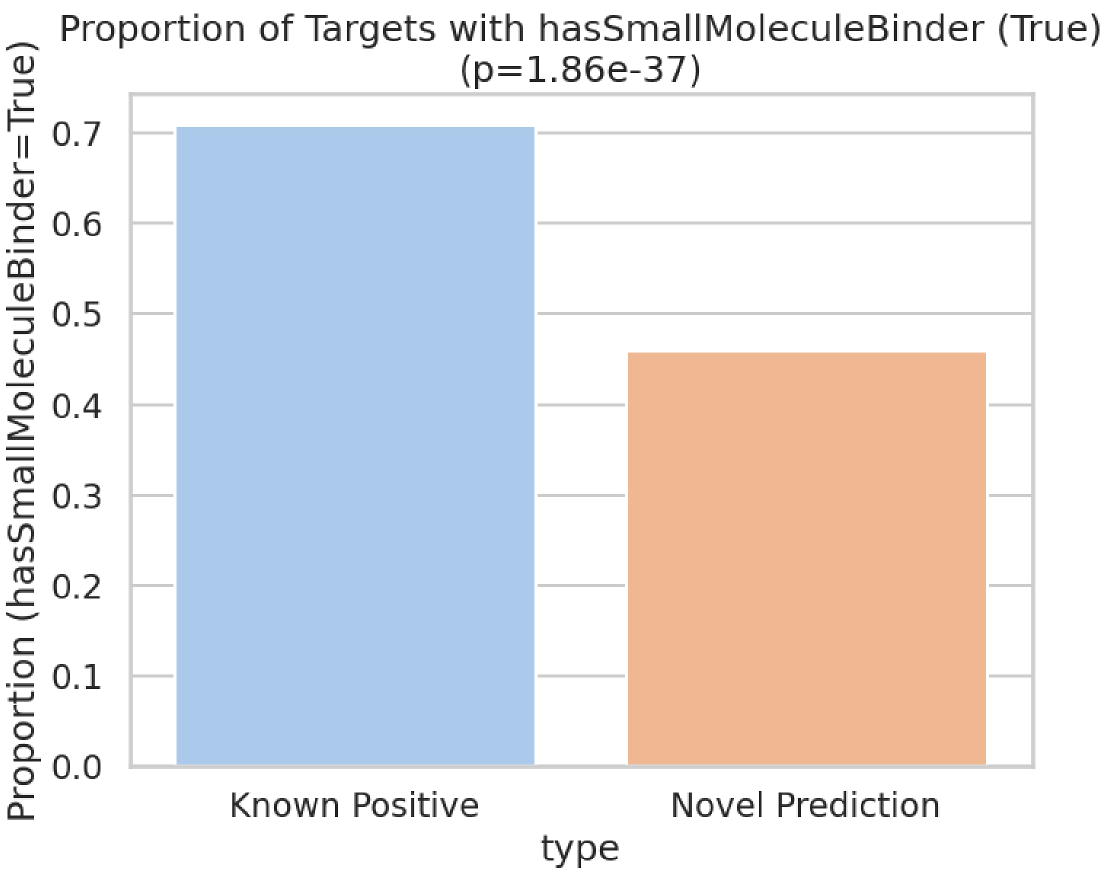


```
--- Rarity Analysis (Rare vs Non-rare) ---
/tmp/ipykernel_903/1865608078.py:180: FutureWarning:

Passing `palette` without assigning `hue` is deprecated and will be removed in
v0.14.0. Assign the `x` variable to `hue` and set `legend=False` for the same
effect.

  sns.boxplot(x=col_name, y='score', data=df_clean, palette='Set2')
```

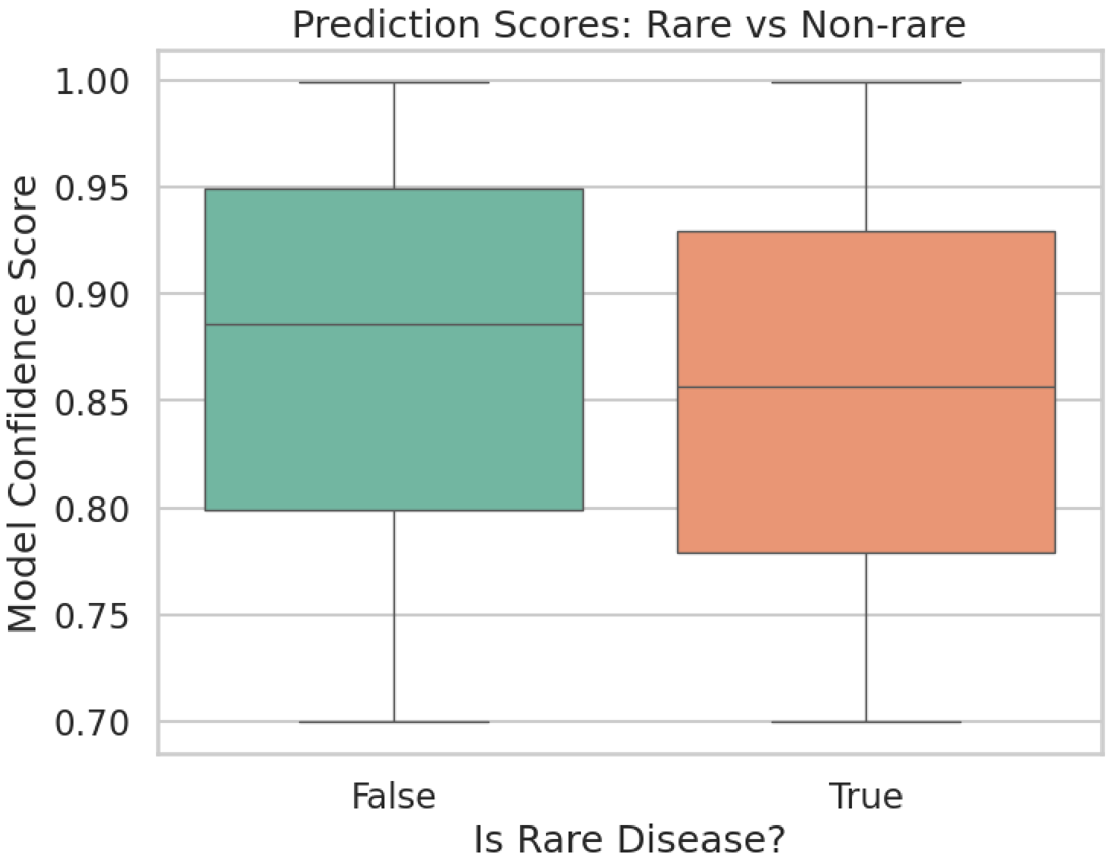


```
Mann-Whitney U Test: P-value = 1.9369e-274
Result: Significant difference.

--- OMIM/Complexity Analysis ---
/tmp/ipykernel_903/1865608078.py:213: FutureWarning:

Passing `palette` without assigning `hue` is deprecated and will be removed in
v0.14.0. Assign the `x` variable to `hue` and set `legend=False` for the same
effect.

  sns.boxplot(x=col_name, y='score', data=df_clean, palette='Set2')
```

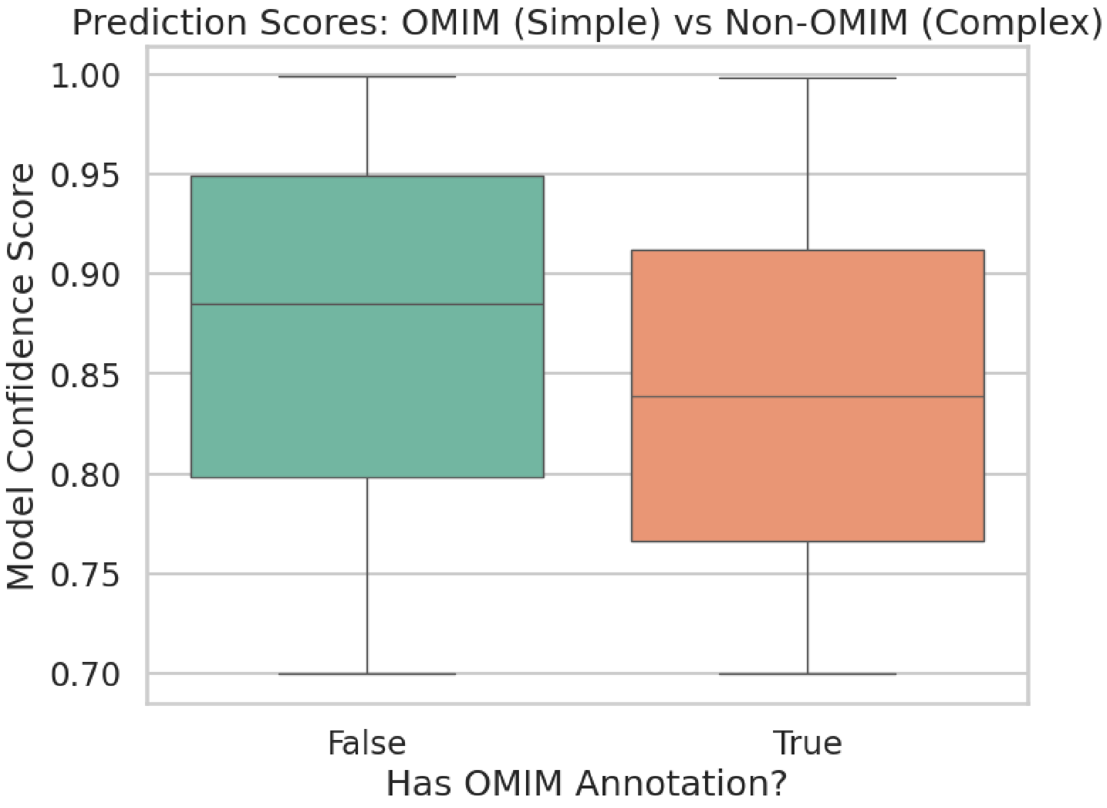


```
Mann-Whitney U Test: P-value = 0.0000e+00
Result: Significant difference.

--- Orphan vs Non-Orphan Analysis ---

/tmp/ipykernel_903/1865608078.py:148: FutureWarning:

Passing `palette` without assigning `hue` is deprecated and will be removed in
v0.14.0. Assign the `x` variable to `hue` and set `legend=False` for the same
effect.

  sns.boxplot(x='orphan', y='score', data=df, palette='Set2')
```

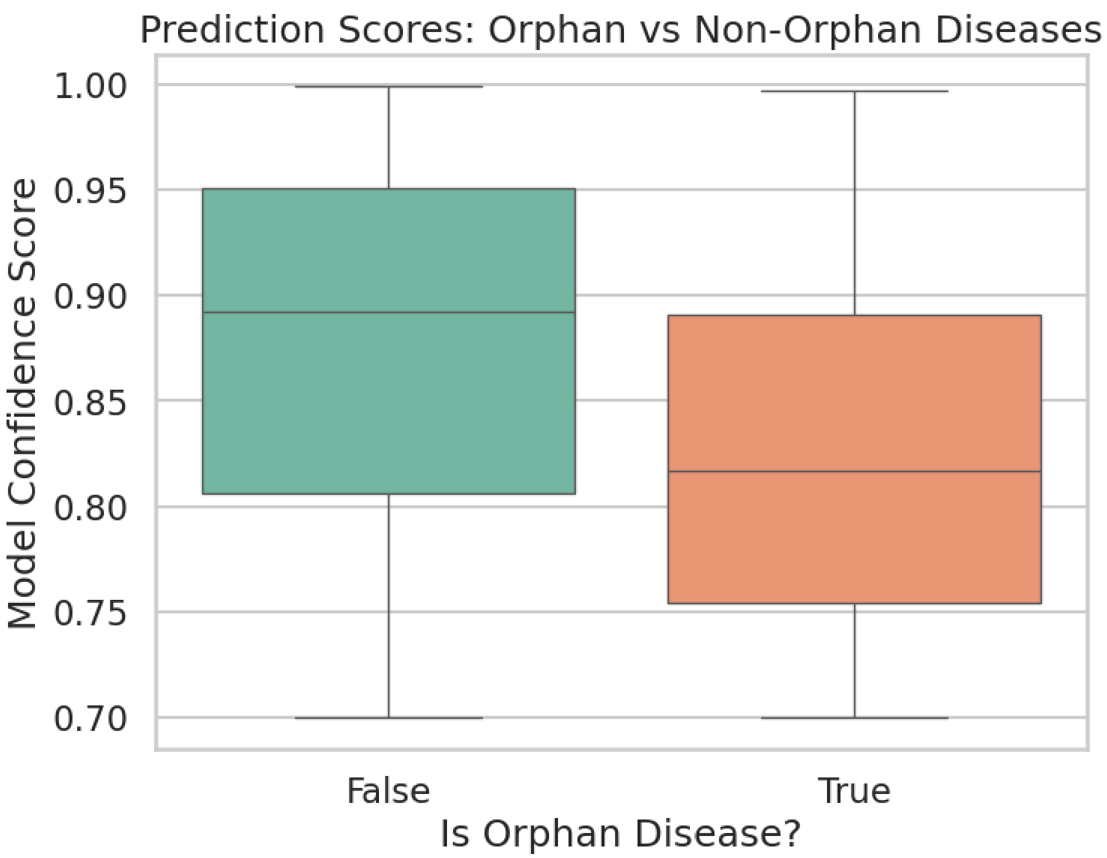


```
Mann-Whitney U Test: P-value = 0.0000e+00
Result: Significant difference.

--- Score Distribution ---
```

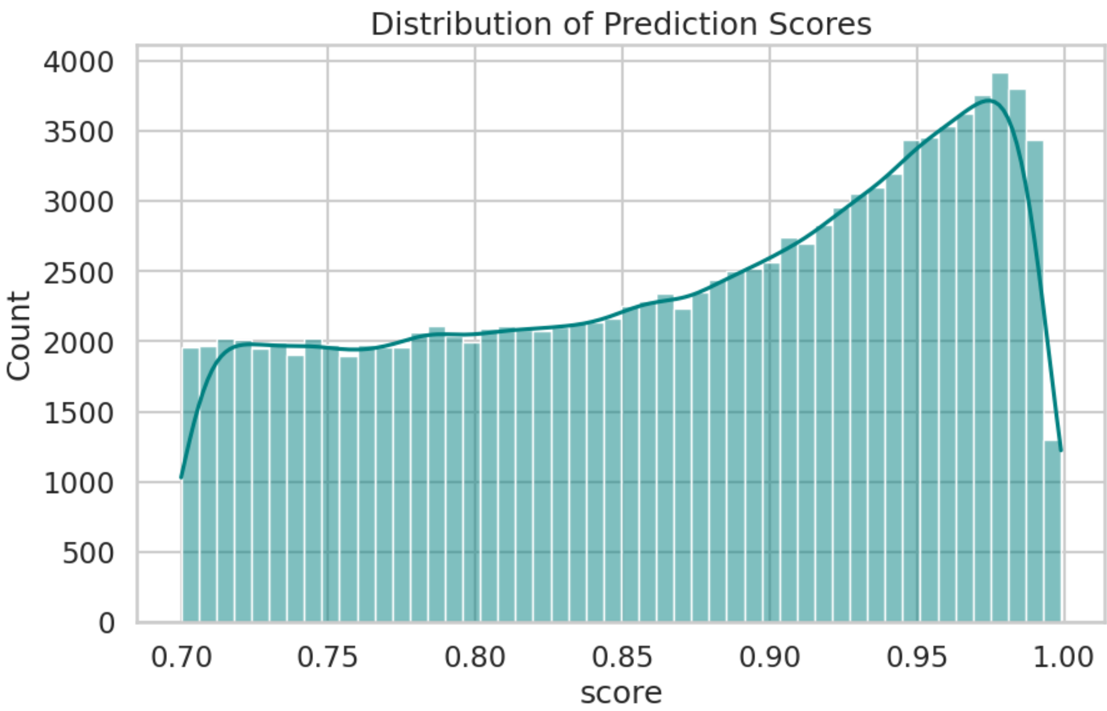


```
count    123112.000000
mean          0.866420
std           0.087011
min           0.700000
25%           0.792000
50%           0.878000
75%           0.945000
max           0.999000
Name: score, dtype: float64
```

```
--- Therapeutic Area Analysis ---
```

```
/tmp/ipykernel_903/1865608078.py:407: FutureWarning:

Passing `palette` without assigning `hue` is deprecated and will be removed in
v0.14.0. Assign the `y` variable to `hue` and set `legend=False` for the same
effect.

  sns.barplot(x=top_tas.values, y=top_tas.index, palette='viridis')
```

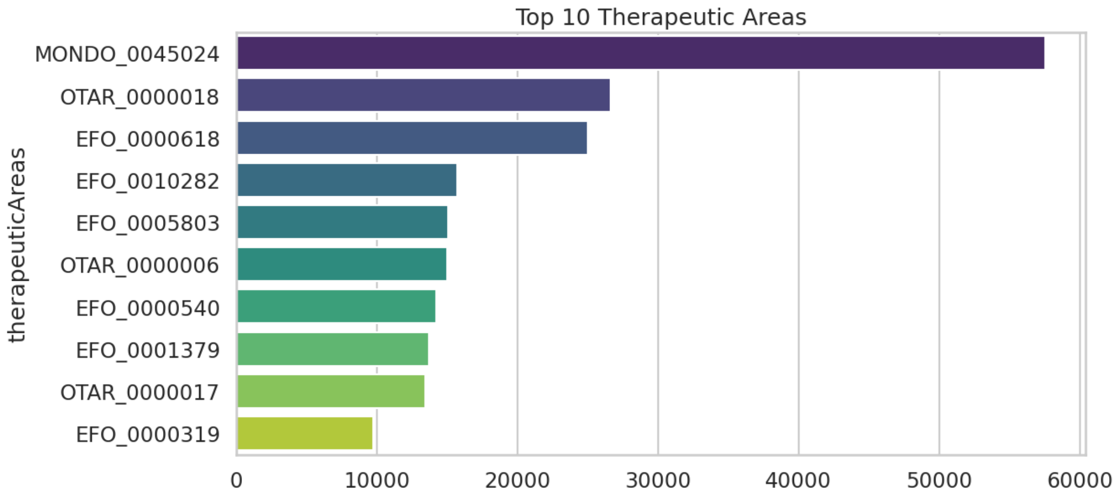


```
--- Repurposing Potential ---
High confidence (score > 0.8) candidates using Approved Drugs: 0

--- Target Class Volume (All Predictions) ---
```

```
/tmp/ipykernel_903/1865608078.py:439: FutureWarning:

Passing `palette` without assigning `hue` is deprecated and will be removed in
v0.14.0. Assign the `y` variable to `hue` and set `legend=False` for the same
effect.

  sns.boxplot(y='targetClass_clean', x='score', data=df_filtered,
order=top_classes, palette='coolwarm')
```

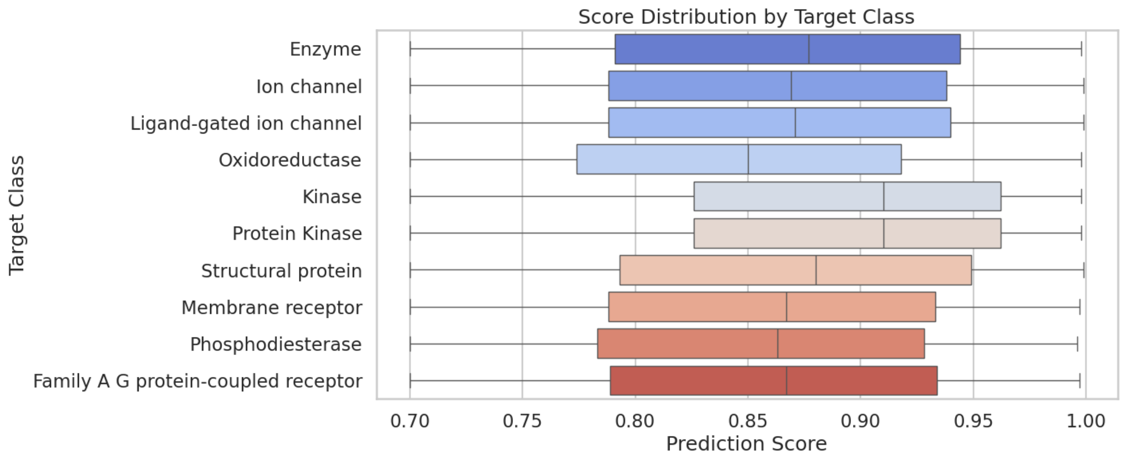

```
--- Target Level Analysis (Unique Targets - General Stats) ---
/tmp/ipykernel_903/1865608078.py:389: FutureWarning:

Passing `palette` without assigning `hue` is deprecated and will be removed in
v0.14.0. Assign the `y` variable to `hue` and set `legend=False` for the same
effect.

  sns.barplot(x=top_classes.values, y=top_classes.index, palette='magma')
```

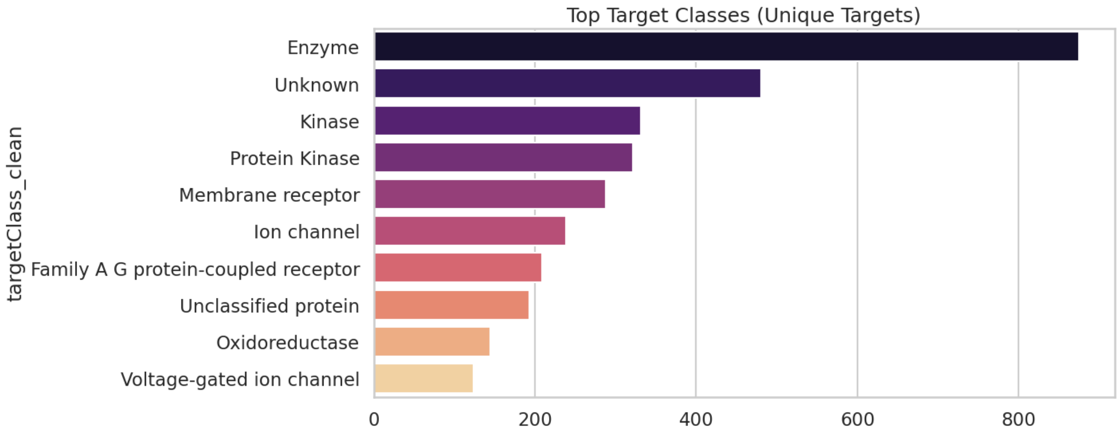


```
/tmp/ipykernel_903/1865608078.py:396: FutureWarning:

Passing `palette` without assigning `hue` is deprecated and will be removed in
v0.14.0. Assign the `x` variable to `hue` and set `legend=False` for the same
effect.

  sns.countplot(x='maxClinicalTrialPhase', data=df_targets, palette='viridis')
```

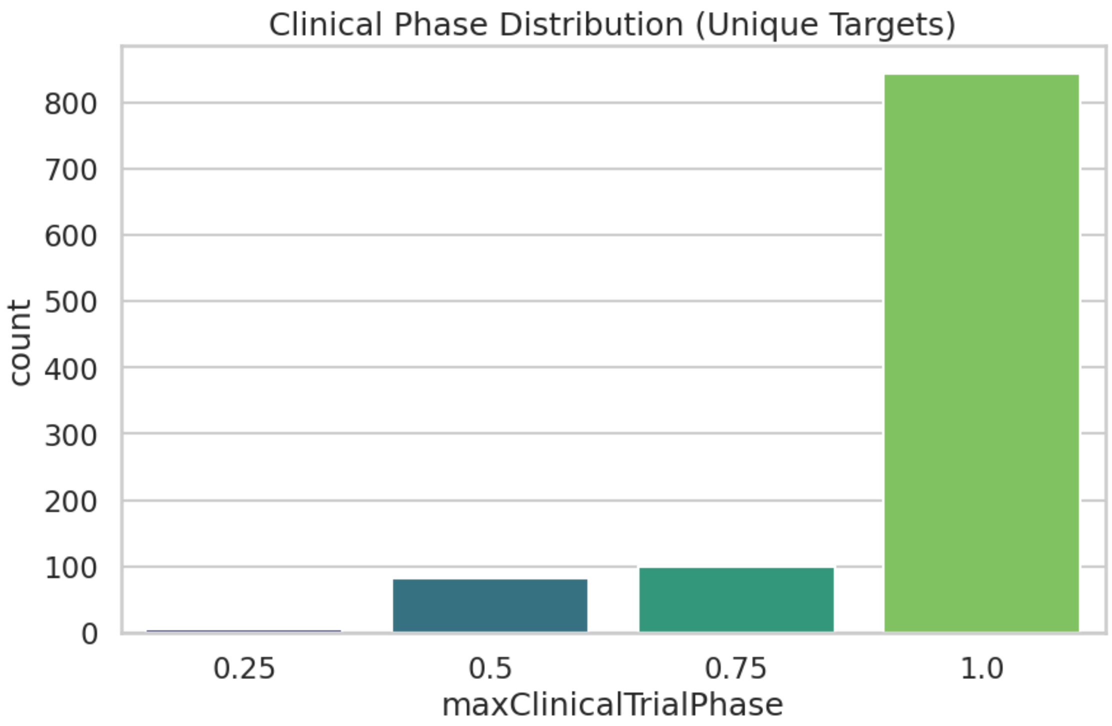


#### 2 More analyses

- e.g. singleton subsets and more

**1. Disease-level coverage & "singleton" diseases** Questions this block answers:

```
How many diseases in OpenTargets have:

no association at all (neither known nor predicted)?

only known positives?

only novel predictions?

both?

Among diseases with exactly one novel prediction, are they more likely to be rare / OMIM?

Do diseases with at least one novel prediction have more or fewer known clinical targets than
```

[25]:
```python
import numpy as np
from scipy import stats

def summarise_disease_coverage_and_singletons(df_preds, disease_df):
```

```
    """
    Disease-level coverage and singleton analysis.

    Assumes:
      - df_preds has columns: diseaseId, label, source, score,
        disease_num_known_clinical_targets, disease_rare, disease_has_omim.
      - disease_df has at least: diseaseId, name, disease_rare,␣
↪disease_has_omim.
    """
    df = df_preds.copy()
    df['is_novel'] = df['label'] < 1
    df['is_known'] = df['label'] > 0

    # Disease metadata universe
    disease_meta = disease_df[['diseaseId', 'name', 'disease_rare',␣
↪'disease_has_omim']].drop_duplicates()

    # Counts per disease
    novel_counts = df[df['is_novel']].groupby('diseaseId').size().
↪rename('n_novel')
    known_counts = df[df['is_known']].groupby('diseaseId').size().
↪rename('n_known')

    # Attach counts to all diseases in OT
    d = disease_meta.merge(novel_counts, on='diseaseId', how='left') \
                    .merge(known_counts, on='diseaseId', how='left')

    d[['n_novel', 'n_known']] = d[['n_novel', 'n_known']].fillna(0).astype(int)
    d['has_any'] = (d['n_novel'] + d['n_known']) > 0
    d['has_novel'] = d['n_novel'] > 0
    d['has_known'] = d['n_known'] > 0

    # Basic coverage stats
    total_diseases = d.shape[0]
    with_any = d['has_any'].sum()
    with_novel = d['has_novel'].sum()
    with_known = d['has_known'].sum()
    only_novel = ((d['has_novel']) & (~d['has_known'])).sum()
    only_known = ((d['has_known']) & (~d['has_novel'])).sum()
    no_preds = (~d['has_any']).sum()

    print("=== Disease-level coverage ===")
    print(f"Total diseases in disease_df: {total_diseases}")
    print(f"Diseases with any association (known or novel): {with_any}")
    print(f"  - with novel predictions: {with_novel}  (only novel:␣
↪{only_novel})")
```

```
    print(f"  - with known positives:   {with_known}  (only known:␣
↪{only_known})")
    print(f"Diseases with NO known or predicted targets: {no_preds}")

    # Rare vs non-rare coverage
    for flag_col, label in [('disease_rare', 'Rare'), ('disease_has_omim',␣
↪'OMIM / simple')]:
        if flag_col not in d.columns:
            continue
        sub = d.dropna(subset=[flag_col])
        for flag_val, desc in [(True, f"{label} diseases"),
                               (False, f"Non-{label} diseases")]:
            tmp = sub[sub[flag_col] == flag_val]
            if tmp.empty:
                continue
            n_total = tmp.shape[0]
            n_with_novel = (tmp['has_novel']).sum()
            n_with_any = (tmp['has_any']).sum()
            print(f"\n{desc}:")
            print(f"  total = {n_total}")
            print(f"  with  1 novel prediction = {n_with_novel} ({n_with_novel/
↪n_total:.2%})")
            print(f"  with any association (known or novel) = {n_with_any}␣
↪({n_with_any/n_total:.2%})")

        # Chi-square: does this flag affect probability of having any novel␣
↪prediction?
        contingency = pd.crosstab(sub[flag_col], sub['has_novel'])
        if contingency.shape == (2, 2):
            chi2, p, _, _ = stats.chi2_contingency(contingency)
            print(f"\nChi-square for {label} vs having  1 novel prediction:␣
↪p={p:.3e}")

    # Attach per-disease "number of known clinical targets" (max over rows in␣
↪df_preds)
    if 'disease_num_known_clinical_targets' in df.columns:
        dk = df.groupby('diseaseId')['disease_num_known_clinical_targets'] \
               .max().rename('n_known_trials')
        d = d.merge(dk, on='diseaseId', how='left')

        # Compare known-trial burden for diseases with vs without novel␣
↪predictions
        trials_with_novel = d.loc[d['has_novel'], 'n_known_trials'].dropna()
        trials_without_novel = d.loc[(~d['has_novel']) & d['has_any'],␣
↪'n_known_trials'].dropna()
```

```python
        if len(trials_with_novel) > 10 and len(trials_without_novel) > 10:
            u, p = stats.mannwhitneyu(trials_with_novel, trials_without_novel, alternative='two-sided')
            print("\nMann-Whitney test on #known clinical targets per disease")
            print("  (diseases WITH vs WITHOUT any novel predictions)")
            print(f"  median WITH = {trials_with_novel.median():.1f}, "
                  f"median WITHOUT = {trials_without_novel.median():.1f}, p={p:.3e}")

    # Singleton vs multi-novel diseases
    single_novel = d[d['n_novel'] == 1]
    multi_novel = d[d['n_novel'] >= 5]  # adjust threshold if you prefer

    print("\n=== Singleton vs multi-novel diseases ===")
    print(f"Diseases with exactly 1 novel prediction: {single_novel.shape[0]}")
    print(f"Diseases with  5 novel predictions:       {multi_novel.shape[0]}")

    # Are singleton diseases more likely to be rare?
    if 'disease_rare' in d.columns:
        cont_single = pd.crosstab(d['disease_rare'], d['n_novel'] == 1)
        if cont_single.shape == (2, 2):
            chi2, p, _, _ = stats.chi2_contingency(cont_single)
            print(f"Chi-square: rarity vs being singleton-novel disease: p={p:.3e}")

    return d  # useful for later slicing / plotting

# Example call (after df_preds & disease_df are loaded as in your notebook)
disease_summary = summarise_disease_coverage_and_singletons(df_preds, disease_df)
```

```
=== Disease-level coverage ===
Total diseases in disease_df: 38959
Diseases with any association (known or novel): 5260
  - with novel predictions: 4907  (only novel: 2931)
  - with known positives:   2329  (only known: 353)
Diseases with NO known or predicted targets: 33699

Rare diseases:
  total = 9156
  with  1 novel prediction = 1896 (20.71%)
  with any association (known or novel) = 2056 (22.46%)

Non-Rare diseases:
  total = 29803
  with  1 novel prediction = 3011 (10.10%)
  with any association (known or novel) = 3204 (10.75%)
```

```
Chi-square for Rare vs having  1 novel prediction: p=2.057e-157

OMIM / simple diseases:
  total = 7499
  with  1 novel prediction = 1382 (18.43%)
  with any association (known or novel) = 1498 (19.98%)

Non-OMIM / simple diseases:
  total = 31460
  with  1 novel prediction = 3525 (11.20%)
  with any association (known or novel) = 3762 (11.96%)

Chi-square for OMIM / simple vs having  1 novel prediction: p=2.976e-64

Mann-Whitney test on #known clinical targets per disease
  (diseases WITH vs WITHOUT any novel predictions)
  median WITH = 0.0, median WITHOUT = 2.0, p=2.471e-34

=== Singleton vs multi-novel diseases ===
Diseases with exactly 1 novel prediction: 1396
Diseases with  5 novel predictions:       2502
Chi-square: rarity vs being singleton-novel disease: p=2.348e-129
```

[ ]:

```
```

**2. Target-level coverage & "singleton" targets** Questions:

```
How many OT targets never appear in your dataset at all?

How many targets have only known positives, only novel predictions, or both?

How many targets are "promiscuous" (lots of diseases) vs "disease-specific" (1-
2 diseases)?

Are predicted targets more "tractable" (e.g. pockets, ligands, clinical phase) than completely
```

[26]:

```
def summarise_target_coverage_and_singletons(df_preds, target_df,␣
↪target_priority):
    """
    Target-level coverage and singleton analysis.

    Assumes:
      - df_preds has columns: targetId, label (or source), diseaseId,␣
↪disease_rare.
      - target_df has at least: targetId, approvedSymbol.
```

```
     - target_priority has target-level features including␣
↪maxClinicalTrialPhase,
       geneticConstraint, paralogMaxIdentityPercentage, etc.
    """
    df = df_preds.copy()
    df['is_novel'] = df['label'] < 1
    df['is_known'] = df['label'] > 0

    # Target universe
    target_meta = target_df[['targetId', 'approvedSymbol']].drop_duplicates()

    # Counts per target
    novel_counts = df[df['is_novel']].groupby('targetId').size().
↪rename('n_novel')
    known_counts = df[df['is_known']].groupby('targetId').size().
↪rename('n_known')

    t = target_meta.merge(novel_counts, on='targetId', how='left') \
                   .merge(known_counts, on='targetId', how='left')
    t[['n_novel', 'n_known']] = t[['n_novel', 'n_known']].fillna(0).astype(int)
    t['has_any'] = (t['n_novel'] + t['n_known']) > 0
    t['has_novel'] = t['n_novel'] > 0
    t['has_known'] = t['n_known'] > 0

    total_targets = t.shape[0]
    with_any = t['has_any'].sum()
    with_novel = t['has_novel'].sum()
    with_known = t['has_known'].sum()
    only_novel = ((t['has_novel']) & (~t['has_known'])).sum()
    only_known = ((t['has_known']) & (~t['has_novel'])).sum()
    no_preds = (~t['has_any']).sum()

    print("=== Target-level coverage ===")
    print(f"Total targets in target_df: {total_targets}")
    print(f"Targets with any association (known or novel): {with_any}")
    print(f"  - with novel predictions: {with_novel}  (only novel:␣
↪{only_novel})")
    print(f"  - with known positives:   {with_known}  (only known:␣
↪{only_known})")
    print(f"Targets with NO known or predicted association: {no_preds}")

    # Singleton vs multi-novel targets
    single_novel = t[t['n_novel'] == 1]
    multi_novel = t[t['n_novel'] >= 5]  # again, threshold tunable

    print("\n=== Singleton vs multi-novel targets ===")
    print(f"Targets with exactly 1 novel disease: {single_novel.shape[0]}")
```

```python
    print(f"Targets with  5 novel diseases:      {multi_novel.shape[0]}")

    # Attach target-prioritisation features for predicted vs never-predicted␣
↪targets
    if 'targetId' in target_priority.columns:
        tp = target_priority.copy()
        # Restrict to one row per targetId by simple 'first' aggregate
        tp = tp.groupby('targetId').first().reset_index()

        predicted_ids = set(df['targetId'].unique())
        tp['is_predicted_any'] = tp['targetId'].isin(predicted_ids)

        num_cols = [
            'geneticConstraint',
            'paralogMaxIdentityPercentage',
            'mouseOrthologMaxIdentityPercentage',
            'mouseKOScore',
            'maxClinicalTrialPhase',
            'tissueSpecificity',
            'tissueDistribution',
        ]
        num_cols = [c for c in num_cols if c in tp.columns]

        print("\n=== Target-prioritisation features: predicted vs never␣
↪predicted ===")
        for col in num_cols:
            pred_vals = tp.loc[tp['is_predicted_any'], col].dropna()
            unpred_vals = tp.loc[~tp['is_predicted_any'], col].dropna()
            if len(pred_vals) < 20 or len(unpred_vals) < 20:
                continue
            u, p = stats.mannwhitneyu(pred_vals, unpred_vals,␣
↪alternative='two-sided')
            print(f"{col}: median(pred)={pred_vals.median():.3f}, "
                  f"median(unpred)={unpred_vals.median():.3f}, p={p:.3e}")

    return t  # again, handy for later

# Example call
target_summary = summarise_target_coverage_and_singletons(df_preds, target_df,␣
↪target_priority)
```

```
=== Target-level coverage ===
Total targets in target_df: 17065
Targets with any association (known or novel): 2613
  - with novel predictions: 2115  (only novel: 1095)
  - with known positives:   1518  (only known: 498)
Targets with NO known or predicted association: 14452
```

```
=== Singleton vs multi-novel targets ===
Targets with exactly 1 novel disease: 541
Targets with  5 novel diseases:       1143

=== Target-prioritisation features: predicted vs never predicted ===
geneticConstraint: median(pred)=-0.189, median(unpred)=0.011, p=2.155e-33
paralogMaxIdentityPercentage: median(pred)=0.000, median(unpred)=0.000,
p=1.108e-74
mouseOrthologMaxIdentityPercentage: median(pred)=0.407, median(unpred)=0.260,
p=9.428e-20
mouseKOScore: median(pred)=-0.647, median(unpred)=-0.321, p=1.825e-52
maxClinicalTrialPhase: median(pred)=1.000, median(unpred)=0.500, p=4.244e-04
tissueSpecificity: median(pred)=0.500, median(unpred)=0.500, p=9.258e-30
tissueDistribution: median(pred)=0.000, median(unpred)=0.000, p=5.823e-18
```

[ ]:

```

```

3. Scores for singleton vs multi-disease / multi-target predictions

Questions:

```
Are "singletons" (only one disease or one target) getting systematically higher or lower score

i.e. does the model reserve highest scores for unique, focused predictions, or for very promis
```

[27]:

```
def score_properties_for_singletons(df_preds):
    """
    Compare score distributions for singleton vs multi-association diseases and␣
↪targets
    (novel predictions only).
    """
    df_novel = df_preds[df_preds['label'] < 1].copy()

    # Disease-level: how many novel disease-target pairs per disease?
    disease_counts = df_novel['diseaseId'].value_counts()
    df_novel['disease_n_novel'] = df_novel['diseaseId'].map(disease_counts)

    scores_single_disease = df_novel.loc[df_novel['disease_n_novel'] == 1,␣
↪'score']
    scores_multi_disease = df_novel.loc[df_novel['disease_n_novel'] > 1,␣
↪'score']

    print("=== Scores for diseases with 1 vs >1 novel prediction ===")
    print(f"n_single_disease = {len(scores_single_disease)}, "
          f"median = {scores_single_disease.median():.3f}")
    print(f"n_multi_disease  = {len(scores_multi_disease)}, "
          f"median = {scores_multi_disease.median():.3f}")
```

```
    if len(scores_single_disease) > 10 and len(scores_multi_disease) > 10:
        u, p = stats.mannwhitneyu(scores_single_disease, scores_multi_disease,␣
↪alternative='two-sided')
        print(f"Mann-Whitney p-value: {p:.3e}")

    # Target-level: number of diseases per target (using same df_novel)
    target_counts = df_novel['targetId'].value_counts()
    df_novel['target_n_novel'] = df_novel['targetId'].map(target_counts)

    scores_single_target = df_novel.loc[df_novel['target_n_novel'] == 1,␣
↪'score']
    scores_multi_target = df_novel.loc[df_novel['target_n_novel'] > 1, 'score']

    print("\n=== Scores for targets with 1 vs >1 novel disease ===")
    print(f"n_single_target = {len(scores_single_target)}, "
          f"median = {scores_single_target.median():.3f}")
    print(f"n_multi_target  = {len(scores_multi_target)}, "
          f"median = {scores_multi_target.median():.3f}")
    if len(scores_single_target) > 10 and len(scores_multi_target) > 10:
        u, p = stats.mannwhitneyu(scores_single_target, scores_multi_target,␣
↪alternative='two-sided')
        print(f"Mann-Whitney p-value: {p:.3e}")

    return df_novel  # carries disease_n_novel and target_n_novel

# Example call
df_novel_with_counts = score_properties_for_singletons(df_preds)
```

```
=== Scores for diseases with 1 vs >1 novel prediction ===
n_single_disease = 1396, median = 0.771
n_multi_disease  = 121716, median = 0.879
Mann-Whitney p-value: 1.528e-282

=== Scores for targets with 1 vs >1 novel disease ===
n_single_target = 541, median = 0.936
n_multi_target  = 122571, median = 0.878
Mann-Whitney p-value: 6.273e-17
```

**4. Targets that are "rare-specialists" vs "common-generalists"** Questions:

```
For each target, what fraction of its novel predictions are for rare diseases?

Are there "rare-specialist" targets (most predictions on Orphanet/ORPHA) vs targets mostly for
```

The merged df_preds already carries disease_rare from disease_df.

[28]:
```
def target_rare_vs_nonrare_mixing(df_preds):
    """
```

```python
    For each target, compute how many of its novel predictions are for rare vs non-rare diseases.
    """
    df_novel = df_preds[df_preds['label'] < 1].copy()
    if 'disease_rare' not in df_novel.columns:
        print("Column 'disease_rare' not found on df_preds; cannot run this analysis.")
        return None

    # Safety: convert to bool
    df_novel['disease_rare'] = df_novel['disease_rare'].astype(bool)

    # Aggregate
    agg = df_novel.groupby('targetId').agg(
        n_pairs=('diseaseId', 'size'),
        n_rare=('disease_rare', 'sum')
    ).reset_index().round(4)
    agg['frac_rare'] = agg['n_rare'] / agg['n_pairs']

    # Attach targetSymbol for readability
    names = df_novel[['targetId', 'targetSymbol']].drop_duplicates()
    agg = agg.merge(names, on='targetId', how='left')

    print("=== Fraction of rare diseases among novel predictions per target ===")
    print(agg['frac_rare'].describe().round(2))

    rare_specialists = agg[(agg['n_pairs'] >= 5) & (agg['frac_rare'] >= 0.8)]
    common_specialists = agg[(agg['n_pairs'] >= 5) & (agg['frac_rare'] <= 0.2)]

    print("\nTop 'rare-specialist' targets ( 80% rare diseases,  5 novel pairs):")
    print(rare_specialists.sort_values('frac_rare', ascending=False)[
        ['targetSymbol', 'targetId', 'n_pairs', 'n_rare', 'frac_rare']
    ].head(20))

    print("\nTop 'common-specialist' targets ( 20% rare diseases,  5 novel pairs):")
    print(common_specialists.sort_values('frac_rare', ascending=True)[
        ['targetSymbol', 'targetId', 'n_pairs', 'n_rare', 'frac_rare']
    ].head(20))

    return agg

# Example call
target_rare_mix = target_rare_vs_nonrare_mixing(df_preds)
```

```
=== Fraction of rare diseases among novel predictions per target ===
count    2115.00
mean        0.33
std         0.32
min         0.00
25%         0.00
50%         0.26
75%         0.50
max         1.00
Name: frac_rare, dtype: float64

Top 'rare-specialist' targets ( 80% rare diseases,  5 novel pairs):
     targetSymbol         targetId  n_pairs  n_rare  frac_rare
922          RPS6  ENSG00000137154        6       6   1.000000
666          SV2C  ENSG00000122012        5       5   1.000000
1627        KCNA3  ENSG00000177272        7       7   1.000000
1104         UGCG  ENSG00000148154        7       7   1.000000
1727        UTP11  ENSG00000183520       16      15   0.937500
1779         SV2B  ENSG00000185518       14      13   0.928571
1117         RPS3  ENSG00000149273       13      12   0.923077
1538         RPS7  ENSG00000171863       13      12   0.923077
509         RPS12  ENSG00000112306       11      10   0.909091
570          RTN4  ENSG00000115310        9       8   0.888889
547         RPL24  ENSG00000114391        9       8   0.888889
781        RPS4Y1  ENSG00000129824        9       8   0.888889
1013       RPL13A  ENSG00000142541        8       7   0.875000
1637        RPS27  ENSG00000177954        8       7   0.875000
110         KCNQ1  ENSG00000053918        8       7   0.875000
940         RPLP1  ENSG00000137818        8       7   0.875000
698          MC3R  ENSG00000124089        7       6   0.857143
388         RPS19  ENSG00000105372        7       6   0.857143
29          CD79B  ENSG00000007312        7       6   0.857143
252         RPLP0  ENSG00000089157        7       6   0.857143

Top 'common-specialist' targets ( 20% rare diseases,  5 novel pairs):
     targetSymbol         targetId  n_pairs  n_rare  frac_rare
1390        GALR1  ENSG00000166573        7       0        0.0
1407         CD3D  ENSG00000167286        8       0        0.0
1414          MVD  ENSG00000167508        6       0        0.0
322       SLCO4A1  ENSG00000101187        8       0        0.0
327        PROKR2  ENSG00000101292        5       0        0.0
419        PCOLCE  ENSG00000106333        5       0        0.0
1560        RXFP4  ENSG00000173080       10       0        0.0
1493        CRTAP  ENSG00000170275        5       0        0.0
1503        GPR27  ENSG00000170837        5       0        0.0
1534          PAH  ENSG00000171759        9       0        0.0
718        PTGER2  ENSG00000125384        8       0        0.0
382          AKT2  ENSG00000105221        8       0        0.0
```

| | | | | | |
|---|---|---|---|---|---|
| 1154 | DST | ENSG00000151914 | 6 | 0 | 0.0 |
| 1055 | NR1I2 | ENSG00000144852 | 6 | 0 | 0.0 |
| 1621 | SLCO3A1 | ENSG00000176463 | 10 | 0 | 0.0 |
| 856 | TRPM1 | ENSG00000134160 | 14 | 0 | 0.0 |
| 762 | GALR3 | ENSG00000128310 | 6 | 0 | 0.0 |
| 766 | SMO | ENSG00000128602 | 5 | 0 | 0.0 |
| 774 | SHBG | ENSG00000129214 | 5 | 0 | 0.0 |
| 639 | GPR68 | ENSG00000119714 | 5 | 0 | 0.0 |

**5. Clinical-phase / "repurposability" of novel targets** Questions:

```
Among novel predictions, how many involve targets that have already been in clinical trials (fo

Are your novel predictions enriched for "already drugged" targets (high maxClinicalTrialPhase)
```

[29]:
```
def target_clinical_phase_vs_novelty(df_preds, target_priority):
    """
    Look at max clinical trial phase for targets involved in novel vs known␣
↪associations.
    """
    df = df_preds.copy()
    df['is_novel'] = df['label'] < 1
    df['is_known'] = df['label'] > 0

    if 'maxClinicalTrialPhase' not in target_priority.columns:
        print("maxClinicalTrialPhase not found in target_priority.")
        return None

    tp = target_priority[['targetId', 'maxClinicalTrialPhase']].copy()
    # collapse to per-target
    tp = tp.groupby('targetId').max().reset_index()

    merged = df.merge(tp, on='targetId', how='left')

    nov = merged.loc[merged['is_novel'], 'maxClinicalTrialPhase'].dropna()
    known = merged.loc[merged['is_known'], 'maxClinicalTrialPhase'].dropna()

    print("=== Clinical trial phase of targets in novel vs known associations␣
↪===")
    print(f"Novel pairs with non-NaN phase: {len(nov)}")
    print(f"Known pairs with non-NaN phase: {len(known)}")

    if len(nov) > 10:
        print(f"Novel: median phase={nov.median():.2f}, "
              f"fraction with phase  1 = {(nov >= 1).mean():.2%}, "
              f"phase  2 = {(nov >= 2).mean():.2%}")
    if len(known) > 10:
        print(f"Known: median phase={known.median():.2f}, "
```

```
            f"fraction with phase  1 = {(known >= 1).mean():.2%}, "
            f"phase  2 = {(known >= 2).mean():.2%}")

    if len(nov) > 10 and len(known) > 10:
        u, p = stats.mannwhitneyu(nov, known, alternative='two-sided')
        print(f"Mann-Whitney test (phase novel vs known): p={p:.3e}")

    return merged

# Example call
df_with_phase = target_clinical_phase_vs_novelty(df_preds, target_priority)
```

```
=== Clinical trial phase of targets in novel vs known associations ===
Novel pairs with non-NaN phase: 97599
Known pairs with non-NaN phase: 67532
Novel: median phase=1.00, fraction with phase  1 = 94.97%, phase  2 = 0.00%
Known: median phase=1.00, fraction with phase  1 = 93.28%, phase  2 = 0.00%
Mann-Whitney test (phase novel vs known): p=6.066e-40
```

[ ]:

#### 2.1 structure, PPI, paralogs analysis

- downloaded with build_target_annotations.py
-

[40]:

```
import numpy as np
import pandas as pd

try:
    from scipy import stats
except ImportError:  # allow running without scipy
    stats = None


def _pick_target_symbol_column(df: pd.DataFrame) -> str:
    """
    Try to find a reasonable 'symbol' column for pretty printing.

    Returns column name or None.
    """
    candidates = [
        "targetSymbol",
        "target_symbol",
        "symbol",
        "geneSymbol",
        "gene_symbol",
```

```python
        "targetSymbol_x",
        "targetSymbol_y",
    ]
    for c in candidates:
        if c in df.columns:
            return c
    return None

def summarize_target_annotation_distribution(df_annot: pd.DataFrame) -> None:
    """
    Quick overview of structural / PPI / paralog features at the *target* level.

    df_annot: disease-target level dataframe, with columns:
        - targetId
        - hasStructure (0/1, PDB coverage)
        - ppiDegree (int)
        - paralogCount (int)
        and optionally a symbol column.
    """
    label_col = _pick_target_symbol_column(df_annot)

    base_cols = ["targetId", "hasStructure", "ppiDegree", "paralogCount"]
    if label_col:
        base_cols.insert(1, label_col)

    missing = [c for c in base_cols if c not in df_annot.columns]
    if missing:
        raise KeyError(f"df_annot is missing columns needed for annotation␣
↪summary: {missing}")

    # one row per target
    t = df_annot[base_cols].drop_duplicates(subset=["targetId"])

    frac_pdb = t["hasStructure"].mean()
    n_pdb = int(t["hasStructure"].sum())
    n_targets = t["targetId"].nunique()

    mean_ppi = t["ppiDegree"].mean()
    median_ppi = t["ppiDegree"].median()

    mean_paralog = t["paralogCount"].mean()
    median_paralog = t["paralogCount"].median()

    frac_zero_ppi = (t["ppiDegree"] == 0).mean()
    frac_zero_paralog = (t["paralogCount"] == 0).mean()

    print("=== Target annotation overview ===")
```

```python
    print(f"# unique targets in df_annot: {n_targets}")
    print(f"Targets with  1 PDB structure: {n_pdb}/{n_targets} "
          f"({frac_pdb:.3f} of targets)")
    print(f"PPI degree (STRING): mean = {mean_ppi:.1f}, median = {median_ppi:.
↪1f}, "
          f"fraction with degree=0: {frac_zero_ppi:.3f}")
    print(f"Paralog count (within-species): mean = {mean_paralog:.2f}, "
          f"median = {median_paralog:.1f}, "
          f"fraction with 0 paralogs: {frac_zero_paralog:.3f}")

    # simple quantiles
    for col in ["ppiDegree", "paralogCount"]:
        q = t[col].quantile([0.0, 0.25, 0.5, 0.75, 0.99])
        print(f"\n{col} quantiles:")
        print(q.to_string())


def analyze_annotations_known_vs_novel(df_annot: pd.DataFrame) -> None:
    """
    Compare annotation features between known clinical-trial targets
    and model-predicted novel targets.

    Assumes `df_annot` has a column `source` with values like
    'known_positive' and 'model_prediction'.
    """
    if "source" not in df_annot.columns:
        raise KeyError("df_annot must have a 'source' column "
                       "(e.g. known_positive vs model_prediction).")

    cols = ["targetId", "hasStructure", "ppiDegree", "paralogCount", "source"]
    missing = [c for c in cols if c not in df_annot.columns]
    if missing:
        raise KeyError(f"df_annot is missing columns needed for known vs novel␣
↪comparison: {missing}")

    # collapse to one row per (targetId, source)
    t = (
        df_annot[cols]
        .drop_duplicates(subset=["targetId", "source"])
        .reset_index(drop=True)
    )

    print("\n=== Target-level annotations: known vs novel ===")
    summary_rows = []

    for src in sorted(t["source"].unique()):
        sub = t[t["source"] == src]
```

```
        frac_pdb = sub["hasStructure"].mean()
        mean_ppi = sub["ppiDegree"].mean()
        median_ppi = sub["ppiDegree"].median()
        mean_paralog = sub["paralogCount"].mean()
        median_paralog = sub["paralogCount"].median()

        print(f"\nSource = {src}  (n targets = {len(sub)})")
        print(f"  PDB coverage fraction: {frac_pdb:.3f}")
        print(f"  PPI degree:   mean = {mean_ppi:.1f}, median = {median_ppi:.
↪1f}")
        print(f"  Paralogs:     mean = {mean_paralog:.2f}, median =␣
↪{median_paralog:.1f}")

        summary_rows.append(
            {
                "source": src,
                "frac_pdb": frac_pdb,
                "mean_ppi": mean_ppi,
                "median_ppi": median_ppi,
                "mean_paralog": mean_paralog,
                "median_paralog": median_paralog,
                "n_targets": len(sub),
            }
        )

    # compact comparison table (nice when glancing)
    summary_df = pd.DataFrame(summary_rows).set_index("source")
    print("\nSummary (means/medians by source):")
    print(summary_df.round(3).to_string())

    if stats is None:
        print("\nscipy is not installed, skipping significance tests.")
        return

    # Mann-Whitney tests where possible
    try:
        known = t[t["source"] == "known_positive"]
        novel = t[t["source"] == "model_prediction"]

        if len(known) > 0 and len(novel) > 0:
            print("\n--- Mann-Whitney U tests (target-level) ---")
            for col, label in [
                ("hasStructure", "PDB coverage"),
                ("ppiDegree", "PPI degree"),
                ("paralogCount", "paralog count"),
            ]:
```

```
                x = known[col].astype(float)
                y = novel[col].astype(float)

                if x.nunique() <= 1 and y.nunique() <= 1:
                    print(f"{label}: both groups constant, skipping test.")
                    continue

                u, p = stats.mannwhitneyu(x, y, alternative="two-sided")
                print(f"{label}: U = {u:.1f}, p = {p:.3e}")
    except Exception as e:
        print(f"Warning: Mann-Whitney tests failed with error: {e}")

def analyze_annotations_by_disease(df_annot: pd.DataFrame) -> None:
    """
    Look at how disease-level properties relate to structural / PPI annotations
    of their associated targets.

    For each disease × source, we compute:
      * num_pairs    = # unique targets
      * frac_pdb     = fraction of targets with PDB structure
      * median_ppi = median PPI degree
      * median_paralogs = median paralog count
    """
    required = ["diseaseId", "targetId", "source", "hasStructure", "ppiDegree",␣
 ↪"paralogCount"]
    missing = [c for c in required if c not in df_annot.columns]
    if missing:
        raise KeyError(
            f"df_annot is missing columns needed for disease-level annotation␣
 ↪analysis: {missing}"
        )

    grouped = (
        df_annot
        .groupby(["diseaseId", "source"])
        .agg(
            num_pairs=("targetId", "nunique"),
            frac_pdb=("hasStructure", "mean"),
            median_ppi=("ppiDegree", "median"),
            median_paralogs=("paralogCount", "median"),
        )
        .reset_index()
    )

    disease_summary = grouped.pivot_table(
        index="diseaseId",
        columns="source",
```

```
        values=["num_pairs", "frac_pdb", "median_ppi", "median_paralogs"],
        fill_value=0,
    )

    # flatten columns
    disease_summary.columns = ["%s__%s" % (m, s) for (m, s) in disease_summary.
↪columns]
    disease_summary = disease_summary.reset_index()

    print("\n=== Disease-level annotation summary (head) ===")
    print(disease_summary.head(10).round(2).to_string(index=False))

    # Global disease-level averages (where columns exist)
    print("\nDisease-level averages (across diseases):")
    for src in ["known_positive", "model_prediction"]:
        frac_col = f"frac_pdb__{src}"
        ppi_col = f"median_ppi__{src}"
        para_col = f"median_paralogs__{src}"
        n_col = f"num_pairs__{src}"

        if frac_col in disease_summary.columns:
            print(f"  [{src}] mean frac_pdb: {disease_summary[frac_col].mean():.
↪3f}")
        if ppi_col in disease_summary.columns:
            print(f"  [{src}] mean median_ppi: {disease_summary[ppi_col].mean():
↪.1f}")
        if para_col in disease_summary.columns:
            print(f"  [{src}] mean median_paralogs: {disease_summary[para_col].
↪mean():.2f}")
        if n_col in disease_summary.columns:
            print(f"  [{src}] mean #targets per disease:␣
↪{disease_summary[n_col].mean():.1f}")

    # Correlation: #novel targets vs fraction with PDB structure (per disease)
    if "num_pairs__model_prediction" in disease_summary.columns and␣
↪"frac_pdb__model_prediction" in disease_summary.columns:
        corr_pdb = disease_summary["num_pairs__model_prediction"].corr(
            disease_summary["frac_pdb__model_prediction"]
        )
        print(
            f"\nCorrelation between #novel targets and fraction with PDB␣
↪structure "
            f"(per disease, model_prediction): {corr_pdb:.3f}"
        )

    # Optional: correlation #novel vs median PPI
```

```python
    if "median_ppi__model_prediction" in disease_summary.columns:
        corr_ppi = disease_summary["num_pairs__model_prediction"].corr(
            disease_summary["median_ppi__model_prediction"]
        )
        print(
            f"Correlation between #novel targets and median PPI degree "
            f"(per disease, model_prediction): {corr_ppi:.3f}"
        )


def run_all_annotation_analyses(df_annot: pd.DataFrame) -> None:
    """
    Convenience function to run all annotation-based analyses on df_annot.
    """
    summarize_target_annotation_distribution(df_annot)
    analyze_annotations_known_vs_novel(df_annot)
    analyze_annotations_by_disease(df_annot)
```

[41]:
```python
df_preds = pd.read_csv("DL_novel_candidates_predictions.csv")  # or your␣
 ↪combined file
target_annotations = pd.read_csv("target_annotations.csv")

df_annot = df_preds.merge(target_annotations, on="targetId", how="left")
run_all_annotation_analyses(df_annot)
```

```
=== Target annotation overview ===
# unique targets in df_annot: 2613
Targets with  1 PDB structure: 777/2613 (0.297 of targets)
PPI degree (STRING): mean = 100.8, median = 54.0, fraction with degree=0: 0.052
Paralog count (within-species): mean = 1.12, median = 0.0, fraction with 0
paralogs: 0.701

ppiDegree quantiles:
0.00      0.00
0.25     20.00
0.50     54.00
0.75    118.00
0.99    717.04

paralogCount quantiles:
0.00     0.00
0.25     0.00
0.50     0.00
0.75     1.00
0.99    15.88

=== Target-level annotations: known vs novel ===
```

```
Source = known_positive  (n targets = 1518)
  PDB coverage fraction: 0.314
  PPI degree:   mean = 134.8, median = 80.0
  Paralogs:     mean = 0.98, median = 0.0

Source = model_prediction  (n targets = 2115)
  PDB coverage fraction: 0.292
  PPI degree:   mean = 95.0, median = 46.0
  Paralogs:     mean = 1.09, median = 0.0

Summary (means/medians by source):
                  frac_pdb  mean_ppi  median_ppi  mean_paralog  median_paralog
n_targets
source
known_positive       0.314   134.794        80.0         0.982             0.0
1518
model_prediction     0.292    94.978        46.0         1.090             0.0
2115

--- Mann-Whitney U tests (target-level) ---
PDB coverage: U = 1641409.5, p = 1.448e-01
PPI degree: U = 2002085.0, p = 4.185e-37
paralog count: U = 1573180.5, p = 1.976e-01

=== Disease-level annotation summary (head) ===
  diseaseId  frac_pdb__known_positive  frac_pdb__model_prediction
median_paralogs__known_positive  median_paralogs__model_prediction
median_ppi__known_positive  median_ppi__model_prediction
num_pairs__known_positive  num_pairs__model_prediction
 DOID_10113                     0.27                        0.34
0.0                               0.0                         48.0
145.0                     11.0                         100.0
 DOID_10718                     0.00                        0.00
0.0                               1.0                          0.0
36.0                      0.0                            1.0
 DOID_13406                     0.26                        0.35
0.0                               0.0                         76.0
161.0                     34.0                          94.0
  DOID_1947                     0.00                        0.00
0.0                               0.5                          0.0
18.0                      0.0                            2.0
  DOID_7551                     0.75                        0.00
0.0                               0.0                         76.0
0.0                      4.0                            0.0
EFO_0000094                     0.26                        0.21
0.0                               0.0                        120.0
48.0                     47.0                           56.0
```

```
EFO_0000095                        0.31                         0.41
0.0                                  0.0                        192.0
84.0                       354.0                         90.0
EFO_0000096                        0.37                         0.30
0.0                                  0.0                        352.0
65.0                       177.0                        100.0
EFO_0000174                        0.35                         0.08
0.0                                  0.0                        128.0
61.0                        43.0                         12.0
EFO_0000178                        0.20                         0.00
0.0                                  3.0                        108.0
37.0                        25.0                          2.0

Disease-level averages (across diseases):
  [known_positive] mean frac_pdb: 0.126
  [known_positive] mean median_ppi: 52.3
  [known_positive] mean median_paralogs: 0.13
  [known_positive] mean #targets per disease: 12.8
  [model_prediction] mean frac_pdb: 0.124
  [model_prediction] mean median_ppi: 41.6
  [model_prediction] mean median_paralogs: 0.16
  [model_prediction] mean #targets per disease: 23.4

Correlation between #novel targets and fraction with PDB structure (per disease,
model_prediction): 0.630
Correlation between #novel targets and median PPI degree (per disease,
model_prediction): 0.485
```

[38]:

```
```

```
=== Target annotation overview ===
# unique targets in df_annot: 2613
Fraction with at least one PDB structure (hasStructure=1): 0.297
Median PPI degree in STRING: 54.0
Median paralog count (within-species): 0.0

ppiDegree quantiles:
0.00      0.00
0.25     20.00
0.50     54.00
0.75    118.00
0.90    242.00
0.99    717.04

paralogCount quantiles:
0.00     0.00
0.25     0.00
0.50     0.00
```

```
0.75     1.00
0.90     4.00
0.99    15.88

=== Target-level annotations: known vs novel ===

Source = known_positive  (n targets = 1518)
  PDB coverage fraction: 0.314
  Median PPI degree:     80.0
  Median paralog count:  0.0

Source = model_prediction  (n targets = 2115)
  PDB coverage fraction: 0.292
  Median PPI degree:     46.0
  Median paralog count:  0.0

--- Mann-Whitney U tests (target-level) ---
hasStructure: U = 1641409.5, p = 1.448e-01
ppiDegree: U = 2002085.0, p = 4.185e-37
paralogCount: U = 1573180.5, p = 1.976e-01

=== Disease-level annotation summary (head) ===
  diseaseId  frac_pdb__known_positive  frac_pdb__model_prediction
median_paralogs__known_positive  median_paralogs__model_prediction
median_ppi__known_positive  median_ppi__model_prediction
num_pairs__known_positive  num_pairs__model_prediction
 DOID_10113                     0.27                        0.34
0.0                               0.0                         48.0
145.0                      11.0                        100.0
 DOID_10718                     0.00                        0.00
0.0                               1.0                          0.0
36.0                       0.0                           1.0
 DOID_13406                     0.26                        0.35
0.0                               0.0                         76.0
161.0                      34.0                         94.0
  DOID_1947                     0.00                        0.00
0.0                               0.5                          0.0
18.0                       0.0                           2.0
  DOID_7551                     0.75                        0.00
0.0                               0.0                         76.0
0.0                       4.0                          0.0
EFO_0000094                     0.26                        0.21
0.0                               0.0                        120.0
48.0                      47.0                         56.0
EFO_0000095                     0.31                        0.41
0.0                               0.0                        192.0
84.0                     354.0                         90.0
EFO_0000096                     0.37                        0.30
```

```
0.0                                  0.0                       352.0
65.0                       177.0                      100.0
EFO_0000174                        0.35                        0.08
0.0                                  0.0                       128.0
61.0                        43.0                       12.0
EFO_0000178                        0.20                        0.00
0.0                                  3.0                       108.0
37.0                        25.0                        2.0

Correlation between #novel targets and fraction with PDB structure (per
disease): 0.630
```

##### 2.1.1 Run Analyses on novel predictions

[34]:
```
# # 2. Filter for NOVEL predictions only (label < 1) as requested for specific␣
↪analyses
# # Adjust logic if your label is strictly -1 for novel
# print("\n--- Filtering for Novel Predictions (label < 1) ---")
# df_novel = df.query("label < 0").copy()
# print(f"Novel Predictions Count: {len(df_novel)}")

# # 3. Run Requested Analyses on Novel set
# analyze_rarity(df_novel)
# analyze_omim(df_novel)
# analyze_orphan(df_novel)
# analyze_score_distributions(df_novel)

# analyze_therapeutic_areas(df_novel)
# analyze_repurposing_potential(df_novel)
# analyze_target_classes(df_novel)

# # 4. Run Strictly Deduplicated Target Analysis
# analyze_target_properties_deduplicated(df_novel)
```

- MONDO:0045024 - cancer

##### 2.1.2 Run Analyses on known positive cases

[35]:
```
# df_known = df.query("label==1")

# # analyze_rarity(df_known)
# # analyze_omim(df_known)
# # analyze_orphan(df_known)
# # analyze_score_distributions(df_known)

# analyze_therapeutic_areas(df_known)
# analyze_repurposing_potential(df_known)
```

```
# # analyze_target_classes(df_known)
```
